# Dissecting the economic impact of soybean diseases in the United States over two decades

**DOI:** 10.1101/655837

**Authors:** Ananda Y. Bandara, Dilooshi K. Weerasooriya, Carl A. Bradley, Tom W. Allen, Paul D. Esker

## Abstract

Soybean (*Glycine max* L. Merrill) is a key commodity for United States agriculture. Here we analyze the economic impacts of 23 common soybean diseases in 28 soybean-producing states in the U.S., from 1996 to 2016. From 1996 to 2016, the total estimated economic loss due to soybean diseases in the U.S. was $81.39 billion, with $68.98 billion and $12.41 billion accounting for the northern and southern U.S. losses, respectively. Across states and years, soybean cyst nematode, charcoal rot, and seedling diseases were the most economically damaging pathogens/diseases while soybean rust, bacterial blight, and southern blight were the least economically damaging. Significantly positive linear correlation of mean soybean yield loss with the mean state-wide soybean production (MT) and mean soybean yield (kg ha^−1^) indicated that high production zones are more vulnerable to soybean diseases-associated yield losses. Our findings provide useful insights into how research, policy, and educational efforts should be prioritized in soybean disease management.

## INTRODUCTION

Soybean [*Glycine max* (L.) Merr.] is a major oilseed crop produced and consumed worldwide and one of the most economically important crops in the United States, the world’s largest soybean producer (Hartman et al. 2011). Soybean is arguably the most versatile row crop on earth. The crop is cultivated on an estimated 6% of the world’s arable land, and since the 1970s, the area under soybean cultivation marked the greatest percentage increase compared to any other major crop (Hartman et al. 2011). Soybean is among the five most important food crops in the world (Savary et al. 2019). The majority of the world’s soybeans are crushed or processed into soybean oil and meal (Ali 2010). Of the oil fraction, 95% is consumed as edible oil while the rest is used for industrial products (Liu 2008). Due to its high protein concentration, approximately 98% of the soybean meal is used in livestock and aquaculture feeds (Hartman et al. 2011). An estimated 2% (= 3 MMT) of the entire global soybean production is consumed by humans as food (Goldsmith 2008). Per the United States Department of Agriculture - National Agricultural Statistics Service (USDA-NASS), the total soybean area planted in the U.S in 2018 was 36 million hectares while the total production was 123 million metric tons. Like the production of other economically important crops, soybean production is constantly challenged by various abiotic and biotic factors including unpredictable weather, diseases, insect pests, weeds, and variable soil quality (Lal 2009; Strange and Scott 2005). Therefore, addressing these issues are important to ensure soybean production profitability while ensuring global food/feed security.

Among the various biotic constraints, plant diseases are detrimental to soybean production, negatively impacting yield and quality. Cultivar selection, environmental conditions, previous disease history, previous crop, and crop management practices are some of the factors that influence the occurrence of soybean diseases (Mueller et al. 2016). The degree of economic damage due to plant diseases depends upon the type of pathogen, environmental conditions, plant tissue affected, number of plants affected, severity of the disease, host plant resistance/susceptibility, plant stress level, and stage of plant development (Hartman and Hill 2010). Losses due to soybean diseases have fluctuated in time and space (location). For instance, the total estimated metric ton lost ranged from 10.07 million in 2012 to 13.94 million in 2014, a difference of 28% (Allen et al. 2017). Despite the spatiotemporal variation, the average annual yield losses due to soybean diseases in the U.S. were estimated to be approximately 11% of the total production (Allen et al. 2017; Hartman et al. 2015).

A variety of practices can be used individually or in combination to mitigate the negative impact of diseases, including, pesticide applications, seed sanitation and cultural techniques, and deployment of resistance. Nonetheless, long-term control of soybean diseases has been challenging due to several reasons including the lack of durable resistance (e.g., soybean cyst nematode - Niblack et al. 2002), lack of resistant cultivars (e.g., soybean rust - Yorinori et al. 2005) or partially resistant cultivars (e.g., *Sclerotinia sclerotiorum -* Kim and Diers 2000), lack of highly efficacious fungicides (e.g., *S. sclerotiorum* - Peltier et al. 2012), development of fungicide resistance (e.g., *Cercospora sojina* variant isolates that are nonresponsive to quinone outside inhibitor (QoI) fungicides - Zhang et al. 2018), and extreme environmental stress (e.g., charcoal rot under drought - Mengistu et al. 2011).

Quantitative information on crop losses is difficult to acquire, sparse, rarely standardized, and a challenge to compile and compare across states, agroecosystems and regions (Esker et al, 2012; Savary et al. 2019; Savary et al. 2006). As such, a precise estimation of soybean yield losses due to diseases is complicated. For example, symptoms caused by viral pathogens often mimic those caused by herbicide injury, while it has been reported that more than 30% yield losses due to the soybean cyst nematode (*Heterodera glycines* Ichinohe) can occur without conspicuous above- ground symptoms (Mueller et al. 2016; Wang et al. 2003). Furthermore, although, both yield (quantity) loss and quality loss are included in the concept of crop losses (Savary et al. 2006), the contribution of disease-associated quality loss towards the total economic loss is frequently neglected due to inadequate information or lack of appropriate loss estimation framework. Despite those challenges/deficiencies, the analysis of historical soybean loss data due to diseases is important to understand the economic impact of diseases, to identify the most and least detrimental pathogens of soybean, and to identify spatiotemporal disease occurrence/progression patterns. Such information is critical for soybean pathologists and breeders, government and funding agencies, and educators to prioritize research, policy, and educational efforts in soybean disease management. Since 1974, soybean disease loss estimates for the southern United States have been published in the annual proceedings of the Southern Soybean Disease Workers. The estimated soybean yield losses due to diseases during the past two decades have been reported in a number of papers including Wrather et al (2001) (1996 to 1998), Wrather et al (2003) (1999 to 2002), Wrather and Koenning (2006) (2003 to 2005), Koenning and Wrather (2010) (2006 to 2009), and most recently Allen et al (2017) (2010 to 2014).

In the current study, we focused on soybean yield losses obtained since 1996 for 23 soybean diseases. The specific time period corresponds to the years just prior to the first genetically modified soybean (using transgenic technology) until current day. The objectives of our study were to (i) estimate the economic losses associated with 23 soybean diseases over a 21 year period (1996 to 2016) among 28 soybean producing states, (ii) identify the most and least damaging soybean diseases in the U.S., (iii) unveil the potential economic loss differences due to diseases among northern and southern regions of the U.S., (iv) investigate the disease-associated economic losses pre- and post- discovery of soybean rust, and (v) to investigate whether disease associated economic losses are variable based upon yield (yield zones), harvest area (harvest zones), and total production (production zones) (see methods section for zone derivation procedure). Such information would serve as the base to formulate technically sound, economically viable, and spatiotemporally sensitive disease management strategies.

## MATERIALS AND METHODS

### Data used for the study

Historical soybean loss data gathered from soybean extension specialists and researchers were used for the study. Data spanned the period from 1996 to 2016 across 28 states (AL, AR, DE, FL, GA, IA, IL, IN, KS, KY, LA, MD, MI, MN, MO, MS, NC, ND, NE, OH, OK, PA, SC, SD, TN, TX, VA, WI). Data collection and reporting methods have been previously described (Allen et al. 2017). Briefly, a spreadsheet was circulated annually to plant pathologists with soybean pathology responsibilities in the states listed above for their estimates of the losses associated with each disease. In addition, categories defined as “other diseases”, “other nematodes” and “virus diseases” were included and defined specifically based on the prevalence of diseases within each defined category observed that were not included in the disease list for each state by year. The methods employed within each state differed with regards to the estimating of losses; however, evaluation of cultivar trials, fungicide efficacy plots, specific troubleshooting or field calls, queries of Extension personnel, statewide plant disease surveys, or plant disease diagnostic laboratory databases were some of the methods employed to arrive at an estimated percentage loss value for each disease. The estimated percentage losses were converted to production losses (metric tons) based on the USDA-NASS annual production estimates for each state. In addition to the soybean yield loss data, we incorporated the following classifiers/categorical predictors: (i) Soybean rust discovery (Pre, Post): based upon the pre- and post-discovery of soybean rust in the contiguous U.S. in November 2004, (ii) Region either north or south (North = Illinois, Indiana, Iowa, Kansas, Michigan, Minnesota, Nebraska, North Dakota, Ohio, Pennsylvania, South Dakota, and Wisconsin; South = Alabama, Arkansas, Delaware, Florida, Georgia, Kentucky, Louisiana, Maryland, Mississippi, Missouri, North Carolina, Oklahoma, South Carolina, Tennessee, Texas, and Virginia.), which was classified based on the two groups collecting data (NCERA-137 and Southern Soybean Disease Workers, respectively), (iii) Yield zone (1 to 4): based upon USDA-NASS estimates at the state level comparing yield (MT/HA) with all states and all years, (iv) Harvest zone (1 to 4): based upon USDA-NASS estimates at state level comparing harvested area (HA) with all states and all years, and (v) Production zone (1 to 4): based upon USDA-NASS estimates at the state level comparing total production (MT) with all states and all years. To classify data points into a particular zone (separately for yield [kg/ha], harvest area [ha], and production [MT]), the entire data set (588 data points = 21 years × 28 states) was first numerically ordered from highest to lowest value. Then, the data points within the minimum to first quartile were classified as Zone 1. Similarly, data points from the first quartile to median, median to third quartile, and > third quartile were respectively classified as zones 2, 3, and 4. As such, note that the defined zones were not based on geography. The zone of a given data point (data point = yield/harvest value for a given state at a given year) was based upon the relative numerical position (in terms of yield, harvest area, or total production) of that data point within the data base. As yield, harvest area, and production for a given state fluctuate over time, the zone classification for a given state can be vary based on year.

Based on preliminary examination of the diseases and nematodes, the following categories were developed: anthracnose, bacterial blight, brown stem rot, Cercospora leaf blight (purple seed stain), charcoal rot, Diaporthe-Phomopsis, downy mildew, frogeye leaf spot, Fusarium wilt, other diseases (which varied by year), Phytophthora root and stem rot, pod and stem blight, Rhizoctonia aerial blight, root-knot nematode and other nematodes (which varied by state and year), Sclerotinia stem rot (white mold), seedling diseases (which varied by state and year), Septoria brown spot, southern blight, soybean cyst nematode, soybean rust, stem canker, sudden death syndrome, and virus diseases (which varied by state and year). See supplementary table 1 for the Latin binomials of each pathogen(s) causing these diseases. We used this approach to accommodate some changes that occurred over the years of data collection, for example the separation of Cercospora leaf blight and purple seed stain, both caused by the same fungi, in more recent surveys.

**Table 1.** Eigenvalues, % variance, and % cumulative variance associated with the principle components (PC).

| Dimension<br>(PC) | Eigenvalue | Variance % | Cumulative<br>variance % |
| --- | --- | --- | --- |
| 1 | 2473.78 | 48.04 | 48.04 |
| 2 | 598.21 | 11.62 | 59.66 |
| 3 | 484.82 | 9.42 | 69.08 |
| 4 | 274.41 | 5.33 | 74.41 |
| 5 | 238.61 | 4.63 | 79.04 |
| 6 | 180.51 | 3.51 | 82.55 |
| 7 | 172.93 | 3.36 | 85.91 |
| 8 | 145.06 | 2.82 | 88.72 |
| 9 | 113.12 | 2.20 | 90.92 |
| 10 | 94.70 | 1.84 | 92.76 |
| 11 | 77.26 | 1.50 | 94.26 |
| 12 | 72.56 | 1.41 | 95.67 |
| 13 | 52.45 | 1.02 | 96.69 |
| 14 | 35.95 | 0.70 | 97.39 |
| 15 | 27.53 | 0.53 | 97.92 |
| 16 | 24.95 | 0.48 | 98.40 |
| 17 | 21.88 | 0.43 | 98.83 |
| 18 | 21.20 | 0.41 | 99.24 |
| 19 | 14.48 | 0.28 | 99.52 |
| 20 | 10.02 | 0.19 | 99.72 |
| 21 | 8.19 | 0.16 | 99.88 |
| 22 | 4.12 | 0.08 | 99.96 |
| 23 | 2.24 | 0.04 | 100.00 |

In addition to the loss data, data on annual values of the Palmer Drought Severity Index (averaged over the entire area of the contiguous 48 states), annual average temperature in the contiguous 48 states, and total annual precipitation in the contiguous 48 states were obtained from the National Oceanic and Atmospheric Administration’s National Centers for Environmental Information (www7.ncdc.noaa.gov.) for the graphical representation of said data. The objective here was to investigate fluctuation of important environmental parameters during the course of this study and to unravel potential relationships between such parameter fluctuations and disease losses over time.

### Estimating economic yield losses per disease

Given that the historical data were provided in the form of losses in terms of metric tons of production, to calculate the economic loss per soybean disease, we first calculated the loss as a percentage based on overall production (in metric tons) per state and year using USDA-NASS data (https://www.nass.usda.gov/Data_and_Statistics/). We then calculated the overall loss (as a percentage) using Padwick’s calculation (Padwick 1956), which is:

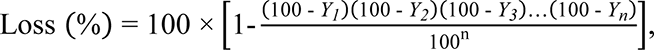

where Y_1_, Y_2_, Y_3_, Y_n_, represent the percentage loss due to disease 1, 2, 3, through n, respectively. To estimate the loss due to diseases in terms of yield, we used the average soybean yield per state and year, from which we estimated the yield in the absence of diseases (the percentage loss estimated using Padwick’s calculation). The difference between the state average yield and the estimated yield in the absence of diseases was considered the loss, from which the estimated loss in dollars per hectare was calculated based on the annual soybean price that was obtained from USDA-NASS. Losses in dollars per hectare per disease was finally estimated by multiplying the losses in dollars per hectare by the percentage of the overall loss due to an individual disease.

### Data imputation

The original data (= computed loss values associated with diseases for states and years) set contained approximately 2.8% missing data. To address missingness, data were imputed using the MICE package in R (Buuren and Groothuis-Oudshoorn 2010). The MICE algorithm deploys the multiple imputation procedure, Multivariate Imputation by Chained Equations (MICE), often referred to as fully conditional specification (FCS) (van Buuren et al. 1999) or sequential regression multiple imputation (SRMI) (Raghunathan et al. 2001). MICE is the key approach for imputing complex incomplete multivariate data. In this approach, multiple imputations on a variable-by-variable basis are created by using a separate model for each incomplete variable where each incomplete variable is regressed on the rest of the variables in the data set.

### Principle components analysis (PCA)

The principal component analysis (PCA) was performed using the FactoMineR and factoextra packages (version 1.41) in R (version 3.5.1) to visualize the principle components (Sebastien Le et al. 2008). The use of PCA allowed the visualization of the distribution pattern of 588 data points (21 years × 28 states) in the principle component space. Since all disease-related variables were standardized and in the same units (dollars), data were not scaled to unit variance (scale.unit = False). As such, non-scaling helped to preserve the inherent variance associated with the data set while avoiding the over penalization of variance inflation control. Therefore, non-scaling ensured the variance maximizing distribution of the data points in principle component space.

### Factor Analysis with Mixed Data (FAMD)

FAMD is a principal component method deployed to analyze a data set containing both quantitative and qualitative variables (Pagès 2004). FAMD makes it possible to analyze the similarity between individuals by taking into account mixed-variable types. With this analysis, quantitative and qualitative variables are normalized in order to balance the impact of each set of variables. The packages FactoMineR (version 1.41, for the analysis) and factoextra (version 1.41, for data visualization) in R (version 9.4, SAS Institute, 2017, Cary, NC) were used for FAMD analysis. Here, total cost due to all diseases was used as the quantitative variable while the year, state, region, soybean rust, yield zone, and harvest zone variables, were incorporated as qualitative variables. Production zone was not included as it was directly derived from yield and harvested area values (thus a linear combination of the yield and harvested area).

### Analysis of variance (ANOVA) and correlation analysis

To investigate the main effects of state, year, pre- and post-discovery of soybean rust, region, yield zone, harvest zone, and production zone on the total economic loss (per hectare basis) due to all diseases, ANOVA was conducted using the PROC GLIMMIX procedure in SAS (version 9.4, SAS Institute, Cary, NC) at the 5% significance level. Restricted maximum likelihood (REML) was used to compute the variance components. Degrees of freedom for the denominator of F tests were computed using the Kenward-Roger option. Assumptions of identical and independent distribution of residuals, and normality of residuals were tested using studentized residual plots and Q-Q plots, respectively. Appropriate heterogeneous variance models were fitted whenever residuals were not homogenously distributed by specifying a “*random residual/group =*” statement (where group = factor under consideration, ex: state). The Bayesian information criterion (model with the lowest BIC) was used to select the best fitting model (between homogenous variance vs heterogeneous variance). Mean separation (all pairwise comparisons of the levels within each factor) was performed with adjustments for multiple comparisons using the Tukey-Kramer test.

The PROC CORR procedure of SAS was used to compute the Pearson correlation coefficients and significance levels between mean soybean yield (kg ha^−1^)/harvest area (ha) and mean soybean yield losses (kg ha^−1^) due to all diseases. Moreover, the correlation coefficient and significance level between mean statewide soybean production (metric tons) and mean state-wide soybean yield losses (metric tons) due to all diseases were computed. Mean values of individual states across 21 years (1996 to 2016) were used for the correlation analysis.

### Grouping diseases for economic loss analysis

In addition to investigating the economic impacts of individual diseases, the 23 diseases considered for this study was grouped into several disease categories in order to investigate their relative economic importance at national, regional, yield/harvest/production zone, and state levels. Categories included were, **Bacterial** (Bacterial blight), **Foliar** (Anthracnose, Cercospora leaf blight (purple seed stain), Diaporthe-Phomopsis, Downy mildew, Frogeye leaf spot, Pod and stem blight, Rhizoctonia aerial blight, Septoria leaf spot, and Soybean rust), **Nematodes** (*Heterodera glycine* (soybean cyst nematode), *Meloidogyne* spp. (root-knot nematodes), *Rotylenchulus reniformis* (reniform nematode), *Belonolaimus longicaudatus* (sting nematode), *Helicotylenchus* spp. (spiral nematodes), *Hoplolaimus* spp. (lance nematodes), *Paratrichodorus* spp. (stubby root nematodes), and *Pratylenchus* spp. (lesion nematodes)), **Stem/Root** (Brown stem rot, Charcoal rot, Fusarium wilt, Phytophthora root and stem rot, Sclerotinia stem rot (white mold), Seedling diseases (caused by a complex of organisms such as multiple species of *Fusarium*, *Pythium*, *Phomopsis*, and/or *Rhizoctonia solani*), Southern blight, Stem canker, and Sudden death syndrome), **Virus** (*Alfalfa mosaic virus, Bean pod mottle virus, Bean yellow mosaic virus, Peanut mottle virus, Soybean dwarf virus, Soybean mosaic virus, Soybean vein necrosis virus, Tobacco ringspot virus, Tobacco streak virus, and Tomato spotted wilt virus*), and **Other diseases** (black root rot, Cercospora leaf blight, *Cylindrocladium parasticum* (red crown rot), green stem syndrome, Neocosmospora root rot, Pythium root rot, target spot, and Texas root rot). Note that all foliar and stem/root categories represent diseases caused by fungi.

## RESULTS

### Identification of the most important soybean diseases and categorical predictors using multivariate approaches

The principle components analysis (PCA) revealed that the percent contribution of the first five principle components (dimensions) to the total explained variance was 48.04, 11.62, 9.42, 5.33, and 4.63, respectively (Table 1), explaining approximately 80% of the total variation. The ten disease variables that contributed most to the first two principle components were brown stem rot, charcoal rot, Diaporthe-Phomopsis, Fusarium wilt, Phytophthora root and stem rot, Sclerotinia stem rot (white mold), seedling diseases, Septoria brown spot, soybean cyst nematode, and sudden death syndrome (Table 2). Among these, the soybean cyst nematode contributed the most (66%) to the first principle component while the second principle component was predominantly represented by charcoal rot (65%).

**Table 2.** Percent contribution of variables to the first five principle components (dimensions = dim.) as revealed via principle component analysis (PCA) and factor analysis with mixed data (FAMD) approaches.

| Variable | PCA (dimensions) |  |  |  |  |
| --- | --- | --- | --- | --- | --- |
|  | Dim.1 | Dim.2 | Dim.3 | Dim.4 | Dim.5 |
| Anthraxnose | 0.01 | 0.18 | 1.18 | 5.14 | 0.26 |
| Bacterial blight | 0.13 | 0.03 | 0.70 | 0.03 | 0.14 |
| Brown stem rot | 1.86 | 1.23 | 0.16 | 0.39 | 5.31 |
| Cercospora leaf blight (purple seed stain) | 0.07 | 0.28 | 2.09 | 6.23 | 2.73 |
| Charcoal rot | 7.36 | 65.91 | 11.40 | 0.14 | 0.02 |
| Diaporthe-Phomopsis | 0.02 | 0.96 | 3.45 | 20.84 | 0.73 |
| Downy mildew | 0.01 | 0.06 | 0.67 | 0.19 | 0.02 |
| Frogeye leaf spot | 0.02 | 0.21 | 0.79 | 14.92 | 4.25 |
| Fusarium wilt | 3.21 | 4.10 | 0.12 | 0.06 | 4.18 |
| Other diseases<sup>a</sup> | 0.03 | 0.02 | 0.02 | 0.20 | 11.74 |
| Phytophthora root and stem rot | 5.13 | 1.21 | 0.80 | 25.26 | 7.04 |
| Pod and stem blight | 0.26 | 0.11 | 1.53 | 1.15 | 0.55 |
| Rhizoctonia aerial blight | 0.00 | 0.10 | 0.08 | 0.43 | 0.12 |
| Root-knot and other nematodes<sup>b</sup> | 0.12 | 0.26 | 0.03 | 12.96 | 1.08 |
| Sclerotinia stem rot (white mold) | 2.52 | 0.20 | 28.63 | 0.90 | 19.46 |
| Seedling diseases<sup>c</sup> | 4.43 | 1.77 | 20.49 | 6.03 | 24.98 |
| Septoria brown spot | 1.90 | 0.44 | 16.44 | 1.37 | 0.46 |
| Southern blight | 0.00 | 0.00 | 0.00 | 0.02 | 0.00 |
| Soybean cyst nematode | 65.93 | 20.68 | 8.17 | 2.33 | 0.55 |
| Soybean rust | 0.01 | 0.05 | 0.01 | 0.14 | 0.06 |
| Stem canker | 0.16 | 0.14 | 0.21 | 0.01 | 0.56 |
| Sudden death syndrome | 6.61 | 1.90 | 2.77 | 1.05 | 15.52 |
| Virus diseases<sup>d</sup> | 0.19 | 0.16 | 0.24 | 0.21 | 0.24 |
| Variable | FAMD (dimensions) |  |  |  |  |
|  | Dim.1 | Dim.2 | Dim.3 | Dim.4 | Dim.5 |
| Harvest zone<sup>e</sup> | 21.06 | 4.84 | 43.50 | 43.39 | 10.01 |
| Region<sup>f</sup> | 21.15 | 6.86 | 0.08 | 1.13 | 0.59 |
| Soybean rust<sup>g</sup> | 3.31 | 35.63 | 1.80 | 1.70 | 1.73 |
| State<sup>h</sup> | 25.70 | 10.28 | 44.75 | 45.41 | 43.52 |
| Total cost due to all diseases | 7.24 | 2.46 | 1.47 | 1.55 | 13.97 |
| Year<sup>i</sup> | 4.16 | 35.95 | 2.77 | 2.08 | 11.73 |
| Yield zone<sup>j</sup> | 17.39 | 3.97 | 5.64 | 4.74 | 18.46 |

<sup>a</sup> Includes: black root rot, Cercospora leaf blight, *Cylindrocladium parasticum* (red crown rot), green stem syndrome, Neocosmospora root rot, Pythium root rot, target spot, and Texas root rot.
<sup>b</sup> Includes: *Rotylenchulus reniformis* (reniform nematode), *Belonolaimus longicaudatus* (sting nematode), and *Meloidogyne* (root-knot nematodes), *Helicotylenchus* (spiral nematodes), *Hoplolaimus* (lance nematodes), *Paratrichodorus* (stubby root nematodes), and *Pratylenchus* spp. (lesion nematodes).
<sup>c</sup> Includes: seedling diseases caused by a complex of organisms such as multiple species of *Fusarium*, *Pythium*, *Phomopsis*, and/or *Rhizoctonia solani*.
<sup>d</sup> Includes: *Alfalfa mosaic virus*, *Bean pod mottle virus*, *Bean yellow mosaic virus*, *Peanut mottle virus*, *Soybean dwarf virus*, *Soybean mosaic virus*, *Soybean vein necrosis virus*, *Tobacco ringspot virus*, *Tobacco streak virus*, and *Tomato spotted wilt virus*.
<sup>e</sup> Represent four levels (zone 1-4) based on the quartiles within a data base containing 588 harvest area (ha) data points (21 years × 28 states). The data points within the minimum to first quartile are classified as harvest zone 1. Similarly, data points from the first quartile to median, median to third quartile, and > third quartile were respectively classified as harvest zones 2, 3, and 4.
<sup>f</sup> Region relates to states contained in each region as either the northern (consisting of IA, IL, IN, KS, MI, MN, ND, NE, OH, PA, SD, and WI) or southern (consisting of AL, AR, DE, FL, GA, KY, LA, MD, MO, MS, NC, OK, SC, TN, TX, and VA) United States.
<sup>g</sup> Soybean rust relates to the years that preceded the discovery of the disease in the contiguous U.S. (1996 to 2004) and years since the discovery (2005 to 2016).
<sup>h</sup> Consisting of the states contained within the study
<sup>i</sup> Consisting of the years contained within the study (from 1996 to 2016).
<sup>j</sup> Represent four levels (zone 1-4) based on the quartiles within a data base containing 588 yield (kg/ha) data points (21 years × 28 states). The data points within the minimum to first quartile are classified as yield zone 1. Similarly, data points from the first quartile to median, median to third quartile, and > third quartile were respectively classified as yield zones 2, 3, and 4.

As revealed via the factor analysis with the mixed data (FAMD) approach, the variables state, region, and harvest zone, respectively contributed the most to the first dimension while year and pre- and post-discovery of soybean rust were the two most contributing factors to the second dimension (Table 2). Although the data points (a data point = total economic loss associated with all disease in a particular year for a given state) did not show a clear clustering pattern based on state, a clear separation was observed with year, where eight years (from 1996 to 2003) were clustered together while the remaining years (from 2004 to 2016) formed a separate cluster (Fig. 1 A, B). Similarly, data points were clearly clustered based upon the region (south and north) as well as the pre- and post-discovery of soybean rust (Fig. 2 A).

**Fig. 1.**
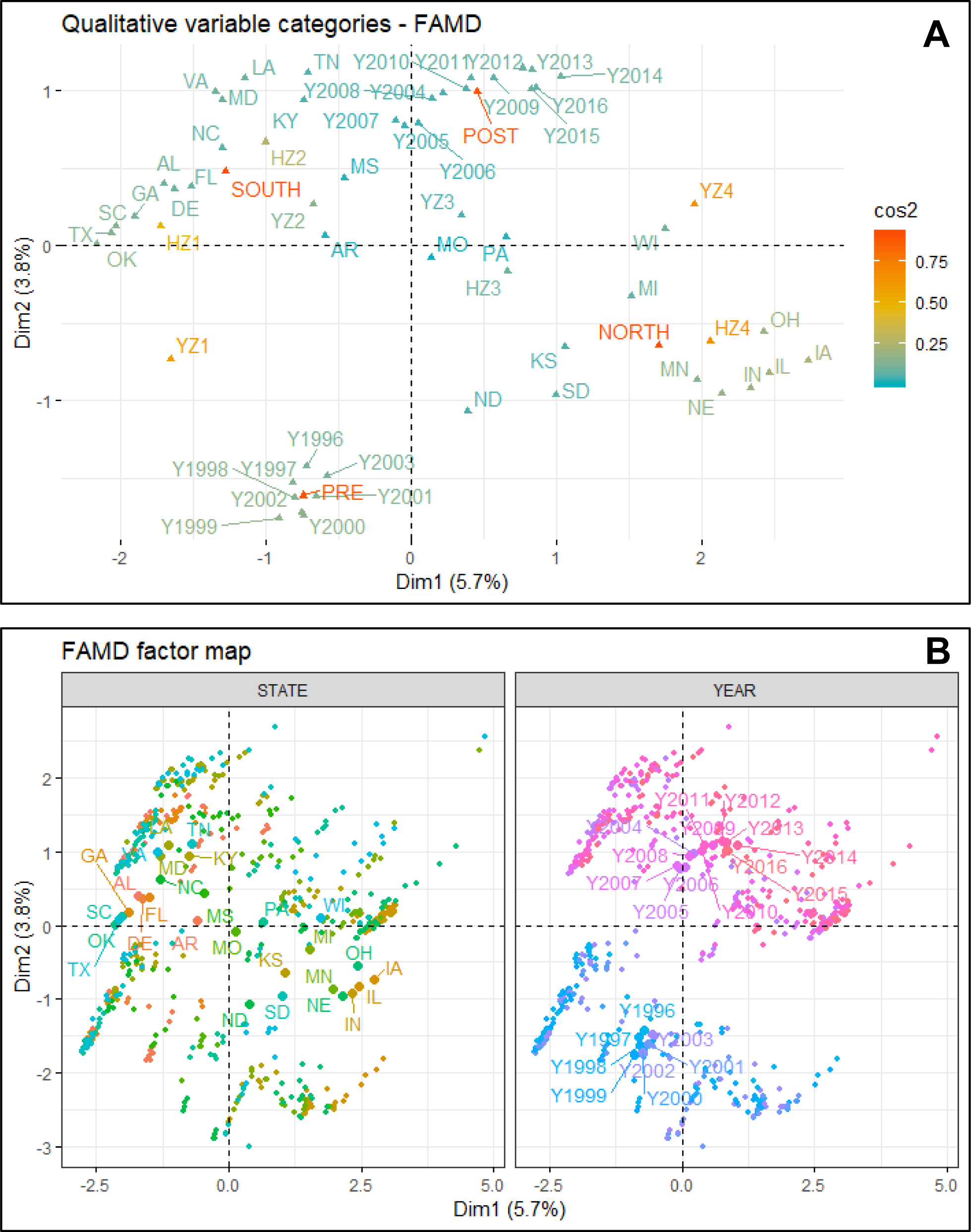
(**A**) The variance maximizing distribution pattern of the qualitative variables (years, states, region, pre- and post-discovery of soybean rust, yield zone, and harvest zone) and (**B**) the FAMD factor maps showing the variance maximizing distribution patterns of the total per hectare economic loss (due to 23 diseases) data points in the map space and their clustering patterns based on states and years. Figures were obtained from the factor analysis with mixed data approach (FAMD analysis). **Diseases** = Anthracnose, bacterial blight, brown stem rot, Cercospora leaf blight (purple seed stain), charcoal rot, Diaporthe-Phomopsis, downy mildew, frogeye leaf spot, Fusarium wilt, other diseases, Phytophthora root and stem rot, pod and stem blight, Rhizoctonia aerial blight, root-knot and other nematodes, Sclerotinia stem rot (white mold), seedling diseases, Septoria brown spot, soybean cyst nematode, soybean rust, southern blight, stem canker, sudden death syndrome, and virus diseases; **Years** = from 1996 to 2016; **States** = Alabama, Arkansas, Delaware, Florida, Georgia, Kentucky, Louisiana, Maryland, Mississippi, Missouri, North Carolina, Oklahoma, South Carolina, Tennessee, Texas, and Virginia, Illinois, Indiana, Iowa, Kansas, Michigan, Minnesota, Nebraska, North Dakota, Ohio, Pennsylvania, South Dakota, and Wisconsin; **Southern region** = Alabama, Arkansas, Delaware, Florida, Georgia, Kentucky, Louisiana, Maryland, Mississippi, Missouri, North Carolina, Oklahoma, South Carolina, Tennessee, Texas, and Virginia; **Northern region** = Illinois, Indiana, Iowa, Kansas, Michigan, Minnesota, Nebraska, North Dakota, Ohio, Pennsylvania, South Dakota, and Wisconsin; **Post-discovery of soybean rust** = from 2004 to 2016; **Pre-discovery of soybean rust** = from 1996 to 2003; **Yield and harvest zones** = represent four levels (zone 1-4) based on the quartiles within a data base containing 588 yield (kg/ha)/harvest area (ha) data points (588 = 21 years × 28 states). Within this data base, data points from the minimum to the first quartile were classified as zone 1. Similarly, data points from the first quartile to median, median to the third quartile, and > third quartile were respectively classified as zones 2, 3, and 4.

**Fig. 2.**
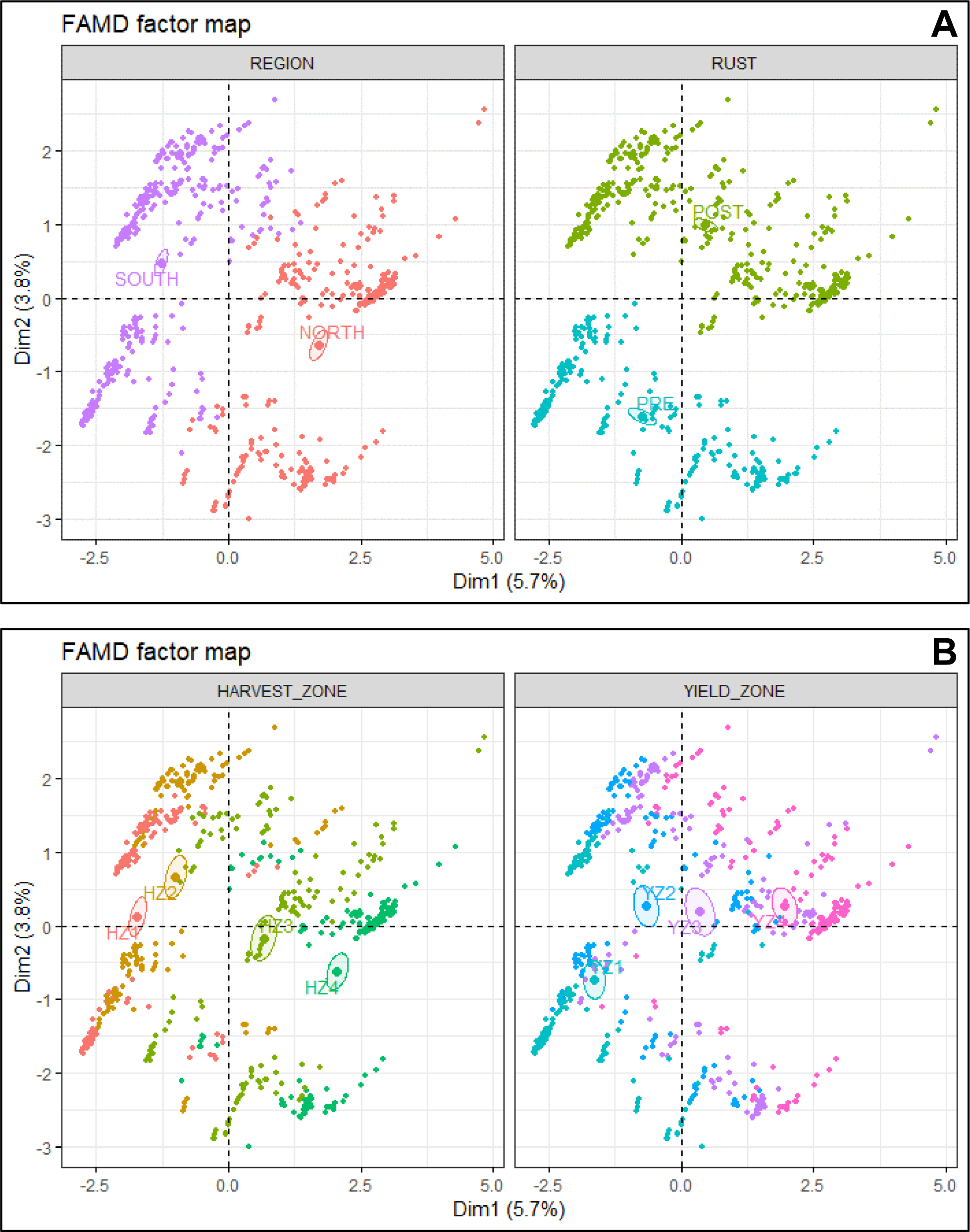
FAMD factor maps obtained from the factor analysis with mixed data approach (FAMD analysis), showing the variance maximizing distribution patterns of the total per hectare economic loss (due to 23 diseases) data points (=588) in the map space and their clustering patterns based on (**A**) region and pre- and post-discovery of rust and (**B**) harvest and yield zones. **Diseases** = Anthracnose, bacterial blight, brown stem rot, Cercospora leaf blight (purple seed stain), charcoal rot, Diaporthe-Phomopsis, downy mildew, frogeye leaf spot, Fusarium wilt, other diseases, Phytophthora root and stem rot, pod and stem blight, Rhizoctonia aerial blight, root-knot and other nematodes, Sclerotinia stem rot (white mold), seedling diseases, Septoria brown spot, soybean cyst nematode, soybean rust, southern blight, stem canker, sudden death syndrome, and virus diseases; **Southern region** = Alabama, Arkansas, Delaware, Florida, Georgia, Kentucky, Louisiana, Maryland, Mississippi, Missouri, North Carolina, Oklahoma, South Carolina, Tennessee, Texas, and Virginia; **Northern region** = Illinois, Indiana, Iowa, Kansas, Michigan, Minnesota, Nebraska, North Dakota, Ohio, Pennsylvania, South Dakota, and Wisconsin; **Post-discovery of soybean rust** = from 2004 to 2016; **Pre-discovery of soybean rust** = from 1996 to 2003; **Yield and harvest zones** = represent four levels (zone 1-4) based on the quartiles within a data base containing 588 yield (kg/ha)/harvest area (ha) data points (588 = 21 years × 28 states). Within this data base, data points from the minimum to the first quartile were classified as zone 1. Similarly, data points from the first quartile to median, median to the third quartile, and > third quartile were respectively classified as zones 2, 3, and 4.

### Analysis of variance (ANOVA) and correlation analysis

The main effects of year, region, state, pre- and post-discovery of soybean rust, yield zone, harvest zone, and production zone were significant on the total economic losses due to all diseases. With respect to the year, the greatest mean losses were observed in 2012 while the lowest were in 2000, a 77% increase between the two years (Fig. 3A). The mean loss in 2012 was significantly greater compared to mean losses in 1996, 1997, 1998, 1999, 2000, 2001, 2002, 2004, 2005, and 2006 (Fig. 3A). The loss in 2011 was significantly greater than that of 2000. Mean losses associated with the rest of the years were not significantly different from each other.

**Fig. 3.**
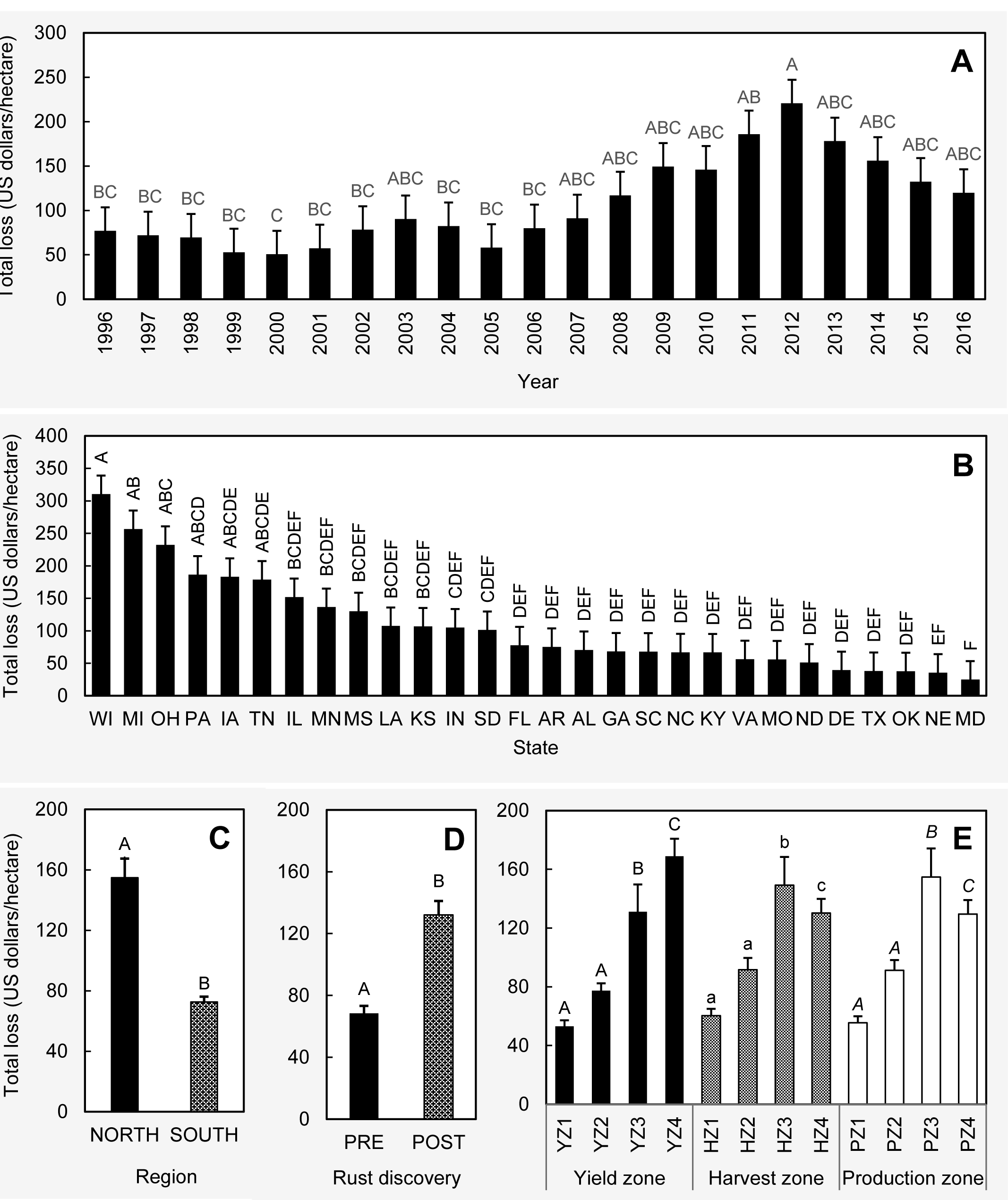
Comparison of the mean total economic losses (within a state, during a given year) due to all 23 diseases considered for this paper among (**A**) years, (**B**) states, (**C**) regions, (**D**) pre- and post-discovery of soybean rust, and (**E**) yield/harvest/production zones. Treatment means with different letter designations (within each sub-figure) are significantly different at α = 0.05 based on the adjustment for multiple comparison using Tukey-Kramer test. Error bars represent standard errors. **Diseases** = Anthracnose, bacterial blight, brown stem rot, Cercospora leaf blight (purple seed stain), charcoal rot, Diaporthe-Phomopsis, downy mildew, frogeye leaf spot, Fusarium wilt, other diseases, Phytophthora root and stem rot, pod and stem blight, Rhizoctonia aerial blight, root-knot and other nematodes, Sclerotinia stem rot (white mold), seedling diseases, Septoria brown spot, soybean cyst nematode, soybean rust, southern blight, stem canker, sudden death syndrome, and virus diseases. **Southern region** = Alabama, Arkansas, Delaware, Florida, Georgia, Kentucky, Louisiana, Maryland, Mississippi, Missouri, North Carolina, Oklahoma, South Carolina, Tennessee, Texas, and Virginia; **Northern region** = Illinois, Indiana, Iowa, Kansas, Michigan, Minnesota, Nebraska, North Dakota, Ohio, Pennsylvania, South Dakota, and Wisconsin; **Post-discovery of soybean rust** = from 2004 to 2016; **Pre-discovery of soybean rust** = from 1996 to 2003; **Yield/Harvest/Production zones** = represent four levels (zone 1-4) based on the quartiles within a data base containing 588 yield (kg/ha)/harvest area (ha)/production (MT) data points (588 = 21 years × 28 states). Within this data base, data points from the minimum to the first quartile were classified as zone 1. Similarly, data points from the first quartile to median, median to the third quartile, and > third quartile were respectively classified as zones 2, 3, and 4.

Wisconsin had the greatest mean losses while the lowest were recorded in Maryland (Fig. 3B). The losses in Wisconsin were 12.5 times greater than the losses observed in Maryland. The mean losses in Wisconsin, Michigan, and Ohio were significantly greater than those observed in Nebraska and Maryland (Fig. 3B). Except for Iowa, Michigan, Ohio, Pennsylvania, and Tennessee, the mean losses of all the other (22) states were significantly lower than that of Wisconsin.

Comparing the two regions across all years, the northern losses were significantly greater in comparison to the south (Fig. 3C). As a percentage, the north reported a 113% greater loss compared to south.

A significantly greater mean loss (93%) was observed in the years following the discovery of soybean rust (2004 to 2016) as compared to the years prior to the discovery (1996 to 2003) (Fig. 3D).

The mean losses associated with yield zone 3 and 4 were significantly greater than that of yield zones 1 and 2, while yield zone 2 had significantly greater losses compared to yield zone 1 (Fig. 3 E). The same scenario was observed with harvest and production zones (Fig. 3E). For yield zone, the greatest losses were observed in yield zone 4, which was 218% greater than the losses observed in yield zone 1 (yield zone with the lowest loss). For harvest zone, the greatest and lowest losses were observed in harvest zones 3 and 1, respectively (147% difference). The same trend was observed with the production zone where zone 3 had a 178% greater mean loss as compared to the loss observed in zone 1.

The mean soybean yield losses in each state, due to all diseases considered in the current study and across the 21 year time period, was positively correlated with the mean state soybean production (in MT, R^2^= 0.88, *P* < 0.0001, Fig. 4A), mean soybean yield (in kg ha^−1^, R^2^ = 0.26, *P* = 0.0052, Fig. 4B), and mean soybean harvest area (in ha, R^2^= 0.15, *P* = 0.0395, Fig. 4C).

**Fig. 4.**
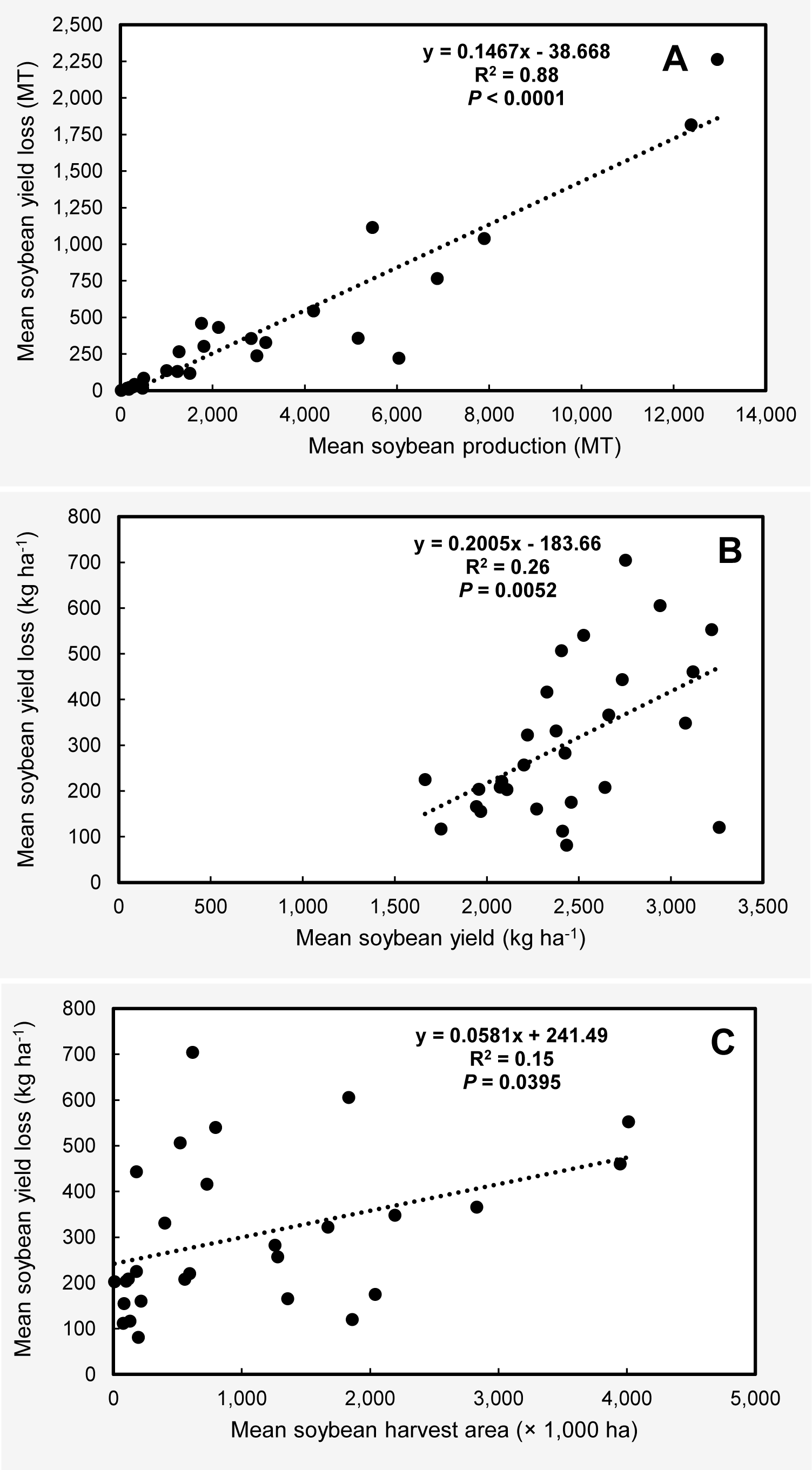
Scatter plots showing the linear relationship between (**A**) mean soybean yield losses and mean state soybean production (both in MT), (**B**) mean soybean yield losses (kg ha^−1^) and mean soybean yield (kg ha^−1^), and (**C**) mean soybean yield loss (kg ha^−1^) and mean soybean harvest area (ha). Each dot represents a state. Mean soybean yield losses represent the losses due all diseases considered in this study (n=23) for individual states between 1996 and 2016.

### Losses associated with soybean diseases

From 1996 to 2016, the total estimated economic loss as a result of soybean diseases in the U.S. was $81.39 billion (Supplementary Tables 2 and 3). Loss in northern states was $68.98 billion while southern states reported a $12.41 billion loss. Therefore, the total economic losses due to soybean diseases in the northern region of the U.S. was 5.6 times greater than that of the southern region. Among all states, Iowa reported the greatest total loss while Maryland reported the lowest. Yearly estimated economic losses (across states) were greatest in 2012 and least in 2001, a reduction in losses between the two years of 70%. Over the 21-year period, the average annual economic loss due to soybean diseases in the U.S. reached nearly $3.88 billion. The same for northern and southern regions were $3.28 and 0.59 billion, respectively. Estimated annual (from 1996 to 2016) soybean economic losses (in million USD) due to individual diseases across 28 states within the United States are presented in the Supplementary Table 4. Estimated cumulative soybean economic losses from 1996 to 2016 (in million USD) as a result of the individual diseases in 12 northern and 16 southern states in the United States are respectively presented in Supplementary Tables 5 and 6. When considered in temporal scale, the total economic loss (USD millions) as well as per hectare loss (USD) due to 23 soybean diseases showed general increasing trend from 1996 to 2012 while both showed a decreasing trend from 2012 to 2016 (Fig. 5A).

**Fig. 5.**
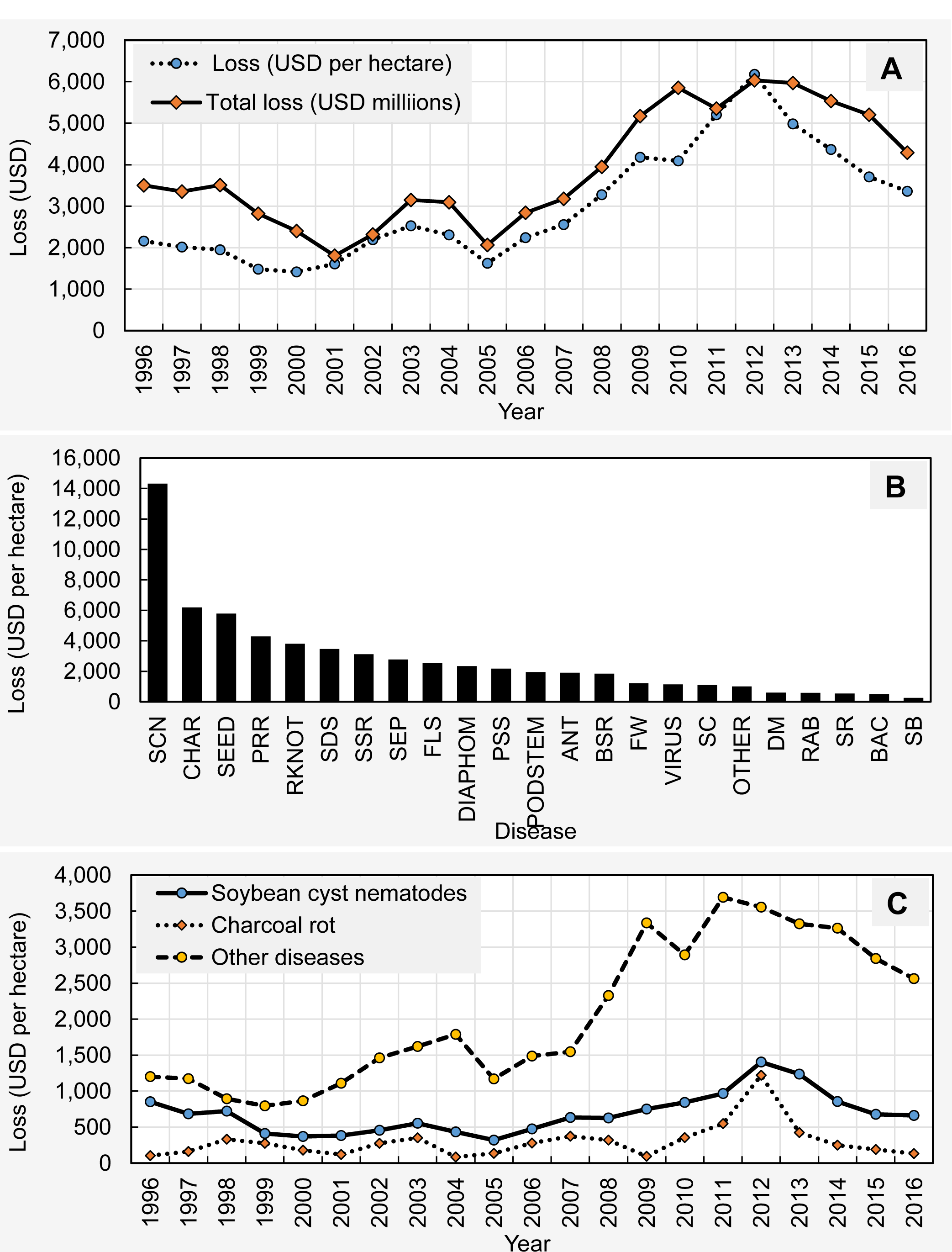
(**A**) Fluctuation of the per hectare (USD) and total (USD millions) economic losses from 1996 to 2016 due to 23 diseases across 28 states (AL, AR, DE, FL, GA, IA, IL, IN, KS, KY, LA, MD, MI, MN, MO, MS, NC, ND, NE, OH, OK, PA, SC, SD, TN, TX, VA, and WI). (**B**) The estimated total economic loss (USD per hectare) associated with 23 diseases across 21 years (1996 to 2016) and 28 states in descending order: SCN = soybean cyst nematodes, CHAR = charcoal rot, SEED = seedling diseases, PRR = Phytophthora root and stem rot, RKNOT = root-knot and other nematodes, SDS = sudden death syndrome, SSR = Sclerotinia stem rot (white mold), SEP = Septoria brown spot, FLS = Frogeye leaf spot, DIAPHOM = Diaporthe-Phomopsis, CLBPSS = Cercospora leaf blight (purple seed stain), PODSTEM = pod and stem blight, ANT = Anthracnose, BSR = Brown stem rot, FW = Fusarium wilt, VIRUS = virus diseases, SC = stem canker, OTHER = other diseases, DM = downy mildew, RAB = Rhizoctonia aerial blight, SR = soybean rust, BAC = bacterial blight, SB = southern blight. (**C**) Fluctuation of the total economic losses (across 28 states) associated with soybean cyst nematode, charcoal rot, and all the other diseases together during 1996 to 2016 period.

On a per hectare basis, the cumulative economic loss due to 23 diseases from 1996 to 2016 across 28 states was $63,409 (Supplementary Table 7). Loss in northern states was $39,006 while southern states reported a $24,002 loss (Supplementary Tables 8, 9; rounding errors may occur). Therefore, the per hectare total economic losses due to soybean diseases in the northern region of the U.S. was 1.6 times greater than that of the southern region. When arranged in descending order, the soybean cyst nematode, charcoal rot, seedling diseases, Phytophthora root and stem rot, root-knot and other nematodes, sudden death syndrome, Sclerotinia stem rot (white mold), Septoria brown spot, frogeye leaf spot, Diaporthe-Phomopsis, Cercospora leaf blight (purple seed stain), pod and stem blight, anthracnose, brown stem rot, Fusarium wilt, virus diseases, stem canker, other diseases, downy mildew, Rhizoctonia aerial blight, soybean rust, bacterial blight, and southern blight were estimated to have the greatest to least economic losses per hectare (across 21 years and 28 states), respectively (Fig. 5B). The total economic losses (per hectare basis, across 28 states) associated with soybean cyst nematode and charcoal rot during the 1996 to 2016 period showed similar fluctuation patterns (Fig. 5C). Losses due to both diseases showed a general increasing trend from 1996 to 2012, when peak economic losses were observed. The heatmap presented in Fig. 6A shows the estimated cumulative economic losses due to each disease from 1996 to 2016 (in U.S. dollars per hectare) for each state. Based on these loss profiles, Wisconsin, Michigan, Iowa, and Ohio clustered separately from the rest of the states. Within this big cluster (consisting of 24 states), no clear sub-clustering was apparent based upon region. Southern and northern states were mixed within this big cluster.

**Fig. 6.**
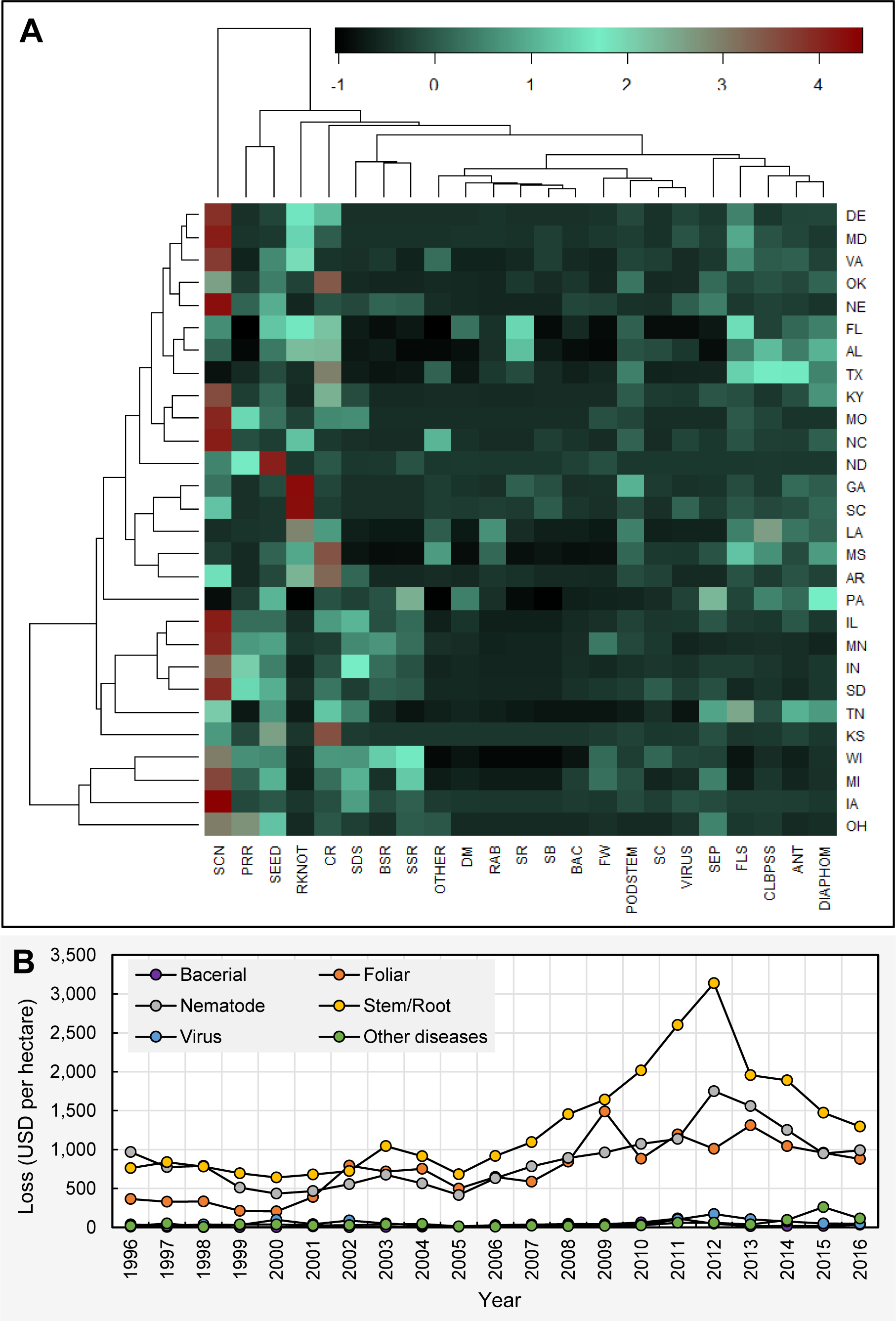
(**A**) Heat map showing the estimated cumulative economic losses (USD per hectare) from 1996 to 2016 associated with 23 diseases for each state. ANT = Anthracnose, BAC = bacterial blight, BSR = Brown stem rot, CHAR = charcoal rot, CLBPSS = Cercospora leaf blight (purple seed stain), DIAPHOM = Diaporthe-Phomopsis, DM = downy mildew, FLS = Frogeye leaf spot, FW = Fusarium wilt, OTHER = other diseases, PODSTEM = pod and stem blight, PRR = Phytophthora root and stem rot, RAB = Rhizoctonia aerial blight, RKNOT = root-knot and other nematodes, SB = southern blight, SC = stem canker, SCN = soybean cyst nematodes, SDS = sudden death syndrome, SEED = seedling diseases, SEP = Septoria brown spot, SR = soybean rust, SSR = Sclerotinia stem rot (white mold), VIRUS = virus diseases. (**B**) Fluctuation of the cumulative economic losses (USD per hectare) of 6 disease categories from 1996 to 2016 across 28 states. **Bacterial** = Bacterial blight; **Foliar** = Anthracnose, Cercospora leaf blight (purple seed stain), Diaporthe-Phomopsis, Downy mildew, Frogeye leaf spot, Pod and stem blight, Rhizoctonia aerial blight, Septoria leaf spot, and Soybean rust; **Nematodes** = *Heterodera glycine* (soybean cyst nematode), *Meloidogyne* spp. (root-knot nematodes), *Rotylenchulus reniformis* (reniform nematode), *Belonolaimus longicaudatus* (sting nematode), *Helicotylenchus* spp. (spiral nematodes), *Hoplolaimus* spp. (lance nematodes), *Paratrichodorus* spp. (stubby root nematodes), and *Pratylenchus* spp. (lesion nematodes); **Stem/Root** = Brown stem rot, Charcoal rot, Fusarium wilt, Phytophthora root and stem rot, Sclerotinia stem rot (white mold), Seedling diseases (caused by a complex of organisms such as multiple species of Fusarium, Pythium, Phomopsis, and/or Rhizoctonia solani), Southern blight, Stem canker, and Sudden death syndrome; **Virus** = *Alfalfa mosaic virus, Bean pod mottle virus, Bean yellow mosaic virus, Peanut mottle virus, Soybean dwarf virus, Soybean mosaic virus, Soybean vein necrosis virus, Tobacco ringspot virus, Tobacco streak virus, and Tomato spotted wilt virus*; **Other diseases** = black root rot, Cercospora leaf blight, *Cylindrocladium parasticum* (red crown rot), green stem syndrome, Neocosmospora root rot, Pythium root rot, target spot, and Texas root rot.

As indicated above, the soybean cyst nematodes caused the greatest total dollar loss (on a per hectare basis) during the period of 1996 to 2016 across the compiled states while southern blight caused the least economic loss in dollars. Furthermore, soybean cyst nematode was estimated to have caused more than twice as much yield loss than any other disease. The total economic losses associated with individual diseases during the time period varied among states/region as well as yield/harvest/production zones. Similarly, the total economic loss in soybean due to diseases in the U.S. varied among years (Supplementary Tables 7, 8, 9, 10). Despite the variation among years, soybean cyst nematode recorded the greatest loss on an annual basis followed by charcoal rot. When considered as regions, losses due to bacterial blight, brown stem rot, downy mildew, Fusarium wilt, Phytophthora root and stem rot, Sclerotinia stem rot (white mold), seedling diseases, Septoria brown spot, soybean cyst nematode, stem canker, sudden death syndrome, and virus diseases were greater in the north compared to south (Supplementary Table 8, 9). However, losses due to anthracnose, Cercospora leaf blight (purple seed stain), charcoal rot, Diaporthe-Phomopsis, frogeye leaf spot, other diseases, pod and stem blight, Rhizoctonia aerial blight, root-knot and other nematodes, southern blight, and soybean rust, were greater in the south as compared to the north. Interestingly, all diseases together recorded a greater loss (across years and states) following the discovery of soybean rust (2004 to 2016) in comparison to pre-discovery (1996 to 2003).

Grouping diseases into categories showed that stem/root, nematode, foliar, virus, other diseases, and bacterial categories caused $27,232, 18,124, 15,427, 1136, 1,004, and 487 per hectare loss at national level (across 28 states and 21 years), respectively. The temporal fluctuation of per hectare economic losses (USD) due to these categories across 28 states is presented in Fig. 6B (see supplementary Table 11 for detailed information). Figure shows that the economic losses due to stem/root diseases have been almost always greater compared to other categories from 1996 to 2016. While nematode diseases caused more losses in certain years, foliar diseases tended to cause more economic damage in certain other years. Virus, other diseases, and bacterial categories showed relatively less economic damages compared to stem/root, nematode, and foliar diseases throughout the 21-year period.

Analysis of the economic damages (USD per hectare) due to disease categories at regional level (across 21 years) showed that stem/root, nematodes, virus, and bacterial disease caused greater losses in northern US compared to southern US (232, 34, 136, and 283 % respectively for each category) (Fig. 7A). However, foliar and other disease categories were greater in south compared to north (49 and 139 % respectively for two categories). The per hectare loss (USD) caused by each disease category is presented in the Supplementary Table 12.

**Fig. 7.**
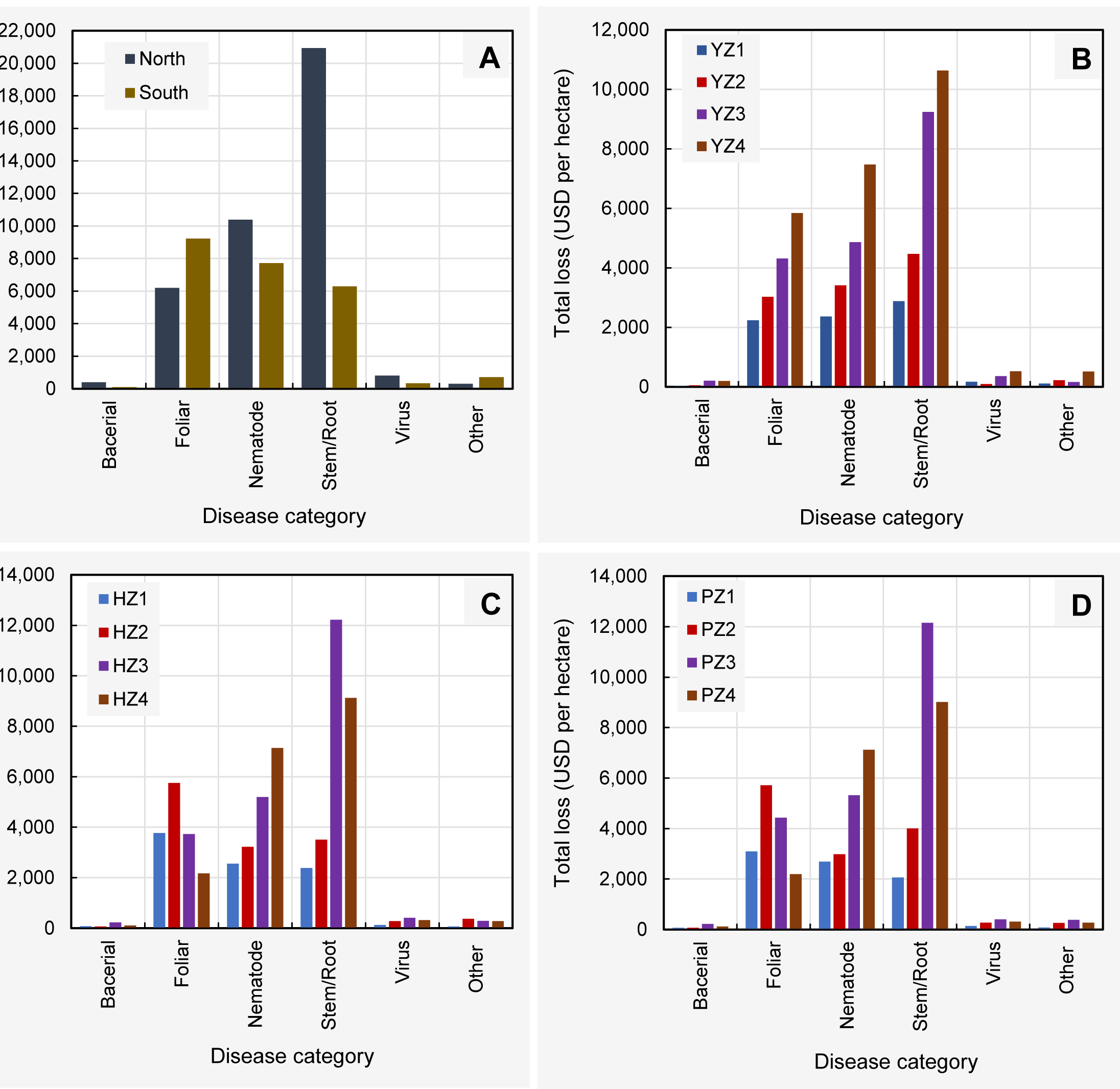
Comparison of the cumulative total per hectare economic loss (USD) from 1996 to 2016 due to disease categories at (**A**) northern and southern US. (**B**) different yield zones (1-4). (**C**) different harvest zones (1-4). (**D**) different production zones (1-4). **Northern region** = Illinois, Indiana, Iowa, Kansas, Michigan, Minnesota, Nebraska, North Dakota, Ohio, Pennsylvania, South Dakota, and Wisconsin; **Southern region** = Alabama, Arkansas, Delaware, Florida, Georgia, Kentucky, Louisiana, Maryland, Mississippi, Missouri, North Carolina, Oklahoma, South Carolina, Tennessee, Texas, and Virginia; **Yield/Harvest/Production zones** = represent four levels (zone 1-4) based on the quartiles within a data base containing 588 yield (kg/ha)/harvest area (ha)/production (MT) data points (588 = 21 years × 28 states). Within this data base, data points from the minimum to the first quartile were classified as zone 1. Similarly, data points from the first quartile to median, median to the third quartile, and > third quartile were respectively classified as zones 2, 3, and 4. **Bacterial** = Bacterial blight; **Foliar** = Anthracnose, Cercospora leaf blight (purple seed stain), Diaporthe-Phomopsis, Downy mildew, Frogeye leaf spot, Pod and stem blight, Rhizoctonia aerial blight, Septoria leaf spot, and Soybean rust; **Nematodes** = *Heterodera glycine* (soybean cyst nematode), *Meloidogyne* spp. (root-knot nematodes), *Rotylenchulus reniformis* (reniform nematode), *Belonolaimus longicaudatus* (sting nematode), *Helicotylenchus* spp. (spiral nematodes), *Hoplolaimus* spp. (lance nematodes), *Paratrichodorus* spp. (stubby root nematodes), and *Pratylenchus* spp. (lesion nematodes); **Stem/Root** = Brown stem rot, Charcoal rot, Fusarium wilt, Phytophthora root and stem rot, Sclerotinia stem rot (white mold), Seedling diseases (caused by a complex of organisms such as multiple species of Fusarium, Pythium, Phomopsis, and/or Rhizoctonia solani), Southern blight, Stem canker, and Sudden death syndrome; **Virus** = *Alfalfa mosaic virus, Bean pod mottle virus, Bean yellow mosaic virus, Peanut mottle virus, Soybean dwarf virus, Soybean mosaic virus, Soybean vein necrosis virus, Tobacco ringspot virus, Tobacco streak virus, and Tomato spotted wilt virus*; **Other diseases** = black root rot, Cercospora leaf blight, *Cylindrocladium parasticum* (red crown rot), green stem syndrome, Neocosmospora root rot, Pythium root rot, target spot, and Texas root rot.

Analysis of the economic damages (USD per hectare) due to disease categories in different yield zones showed that stem/root, nematode, and foliar diseases caused greater losses in yield zone 4 followed by yield zones 3, 2, and 1 (Fig. 7B). The losses caused by bacterial, other diseases, stem/root, nematode, virus, and foliar diseases in yield zone 4 were 450, 389, 269, 215, 204, and 161% greater than those of in yield zone 1. Losses associated with disease categories showed similar patterns between harvest and production zones (Fig. 7 C, D). The foliar diseases appeared to cause more damage in harvest/production zone 2, followed by zones 3, 1, and 4 respectively. The loss due to nematode diseases was greatest in harvest/production zone 4, followed by zones 3, 2, and 1 respectively. The loss associated with stem/root diseases was highest in harvest/production zone 3, followed by zones 4, 2, and 1 respectively (see supplementary Table 13 for detailed information).

## DISCUSSION

Soybean diseases are caused by a complex set of organisms. Previously, Yang and Feng (2001) reported that soybean diseases tend to be more diverse in the northern U.S. than in the southern U.S. with increasing latitude. Soybean diseases can be regional in their occurrence. For instance, in the current study, the losses associated with diseases such as brown stem rot were more pronounced in the northern U.S. while the losses associated with Rhizoctonia aerial blight were more pronounced in the southern U.S. In general, the losses (on a per hectare basis) as a composite within each category were greater in the north than in the south except in the foliar disease category which is comprised of visually elegant diseases that infect the leaves, upper stems and pods and “other diseases” category. In fact, the losses caused by foliar diseases were approximately 50% greater in the southern U.S. whereas the other diseases based on categories ranged from 34% greater in the north (nematodes) to 283% greater in the north than the south (bacterial diseases).

In the current study, the principle component analysis and general analysis of losses associated with soybean diseases revealed the soybean cyst nematode (*Heterodera glycines* Ichinohe) and charcoal rot as important disease barriers for soybean production and profitability in the U.S. In addition, a recent report by Savary et al. (2019) revealed that soybean cyst nematode and charcoal rot are among the top yield loss causing diseases in soybean on a global scale. In the current study, these two diseases accounted for approximately 32% of the total economic losses resulting from all soybean diseases considered (on a per hectare basis). Previous reports by Wrather and Koenning (2006) and Koenning and Wrather (2010) emphasized the importance of these diseases with soybean yield losses. In fact, these two diseases were recently ranked as among the most damaging soybean diseases in the U.S. and Canada (Allen et al. 2017). Our results for the period in question (1996 to 2016) confirmed that losses due to soybean cyst nematode were more prominent in northern states (with the exception of Pennsylvania) while losses associated with charcoal rot were more prominent in southern states, even though both pathogens were reported from every state considered in this research (Allen et al. 2017; Koenning 2005; Koenning 2009). Moreover, the actual economic losses associated with soybean cyst nematode could be understated due to the reliance on aboveground symptom expression for loss assessment as well as the need for soil samples to diagnose the issue. More than 30% yield loss can occur in soybean cyst nematode infested fields without noticeable aboveground symptoms (Mueller et al. 2016; Wang et al. 2003).

Although the charcoal rot pathogen *Macrophomina phaseolina* (Tassi) Goid. likely infects soybean under a broad range of environmental conditions, high disease severity associated yield losses are more pronounced in areas where soils are dry or under persistent drought throughout the soybean reproductive growth stages, generally considered to be periods of stress (Mengistu et al. 2011). The estimated Palmer Drought Severity Index (Fig. 8A), annual average temperature anomaly (Fig. 8B), and total annual precipitation anomaly (Fig. 8C) across the contiguous 48 states showed that 2012 was a year with severe drought, higher temperatures, and lower precipitation. Therefore, environmental conditions that occurred during 2012, as related to drought severity, could, in part explain the severe effect observed, especially when combined with high soybean prices in 2012 and greater than average yield losses as a result of charcoal rot (Fig. 8D).

**Fig. 8.**
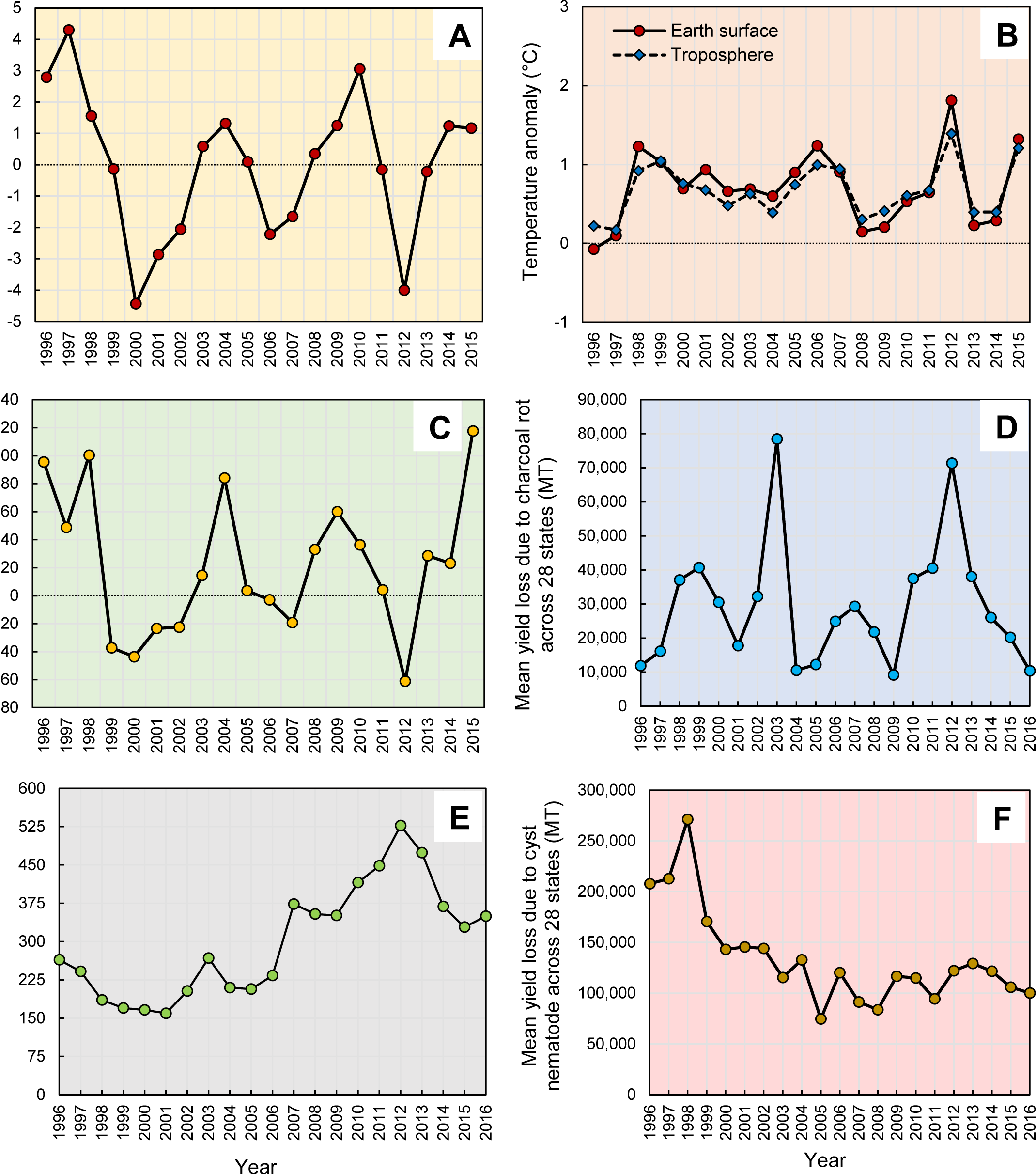
(A) Annual values of the Palmer Drought Severity Index, averaged over the entire area of the contiguous 48 states. Positive values represent wetter-than-average conditions, while negative values represent drier-than-average conditions. A value between −2 and −3 indicates moderate drought, −3 to −4 is severe drought, and −4 or below indicates extreme drought. (**B**) Annual average temperature in the contiguous 48 states. (**C**) Total annual precipitation in the contiguous 48 states. Graph uses the 1901 to 2000 average as a baseline for depicting change. Selecting a different baseline period would not change the shape of the data over time. Data for figures were obtained from the National Oceanic and Atmospheric Administration’s National Centers for Environmental Information (www7.ncdc.noaa.gov.). (**D**) The annual soybean price fluctuation in the United States from 1996 t0 2016 (price for a given year is the mean value across 28 soybean growing states (AL, AR, DE, FL, GA, IA, IL, IN, KS, KY, LA, MD, MI, MN, MO, MS, NC, ND, NE, OH, OK, PA, SC, SD, TN, TX, VA, and WI). (**E**) Mean soybean yield loss due to charcoal rot disease across 28 states from 1996 to 2016. (**F**) Mean soybean yield loss due to soybean cyst nematode across 28 states from 1996 to 2016.

The rationale behind the peak economic losses due to soybean cyst nematode in the drought year 2012 appeared to be the high soybean price (Fig. 8E), despite the reduced yield losses (Fig. 8F) observed as a result of the soybean cyst nematode. The literature on the effect of drought on soybean cyst nematode mediated yield losses is inconsistent. For example, the numbers of soybean cyst nematode eggs, infective juveniles, and cysts were reported to not be affected by soil moisture in the presence of soybean (Barker and Koenning 1989). Moreover, soybean plants respond to moisture stress by increasing root biomass, which would in turn promote soybean cyst nematode reproduction, leading to increased population densities under drought conditions (Barker and Koenning 1989; Koenning et al. 1988; Koenning and Barker 1995). These results support the increased yield losses due to the soybean cyst nematode in years exacerbated by drought years/locations. However, in the absence of soybean (host), survival of the soybean cyst nematode in the soil is related to soil moisture, being greatest at field capacity, followed by dry soil, and least by flooded conditions (Slack et al. 1972). As the nematode migration and population size within fields are restricted by drought, a reduced incidence of disease and subsequent yield losses associated with the pest are possible due to soybean cyst nematode. This might elucidate the reduced soybean cyst nematode-associated yield losses observed in the drought year 2012.

Following the peak losses observed during 2012, economic losses due to charcoal rot as well as soybean cyst nematode declined from 2012 to 2016. The same trend was observed with the losses due to other diseases. Other than the declining soybean prices, the declining economic losses trend could be contributed by less stressful growth conditions, improved management practices and/or use of resistant cultivars or simply as a result of improved soybean genetics. Previous reports, along with the findings of the current study showed the importance of devising spatially sensitive disease management packages for controlling soybean cyst nematode and charcoal rot.

The results of the factor analysis with mixed data (clear and separate clustering of factor levels) agreed with the ANOVA results where significant differences were observed between levels of most factors when the total loss due to all diseases was considered as the response variable. For instance, intuitively the total economic loss due to all diseases (per hectare as well as cumulative total loss) was significantly greater in the northern U.S. as compared to the southern U.S. Although the disparities in climatic conditions may have contributed to such economic loss differences, it would be useful to conduct a comprehensive study to look at the management practices such as planting dates, plant population densities, use of resistant/tolerant soybean cultivars, crop rotation, tillage practices, fungicide application (timing, active ingredients, frequency, etc.), fertilizer application (timing, type of nutrients, etc.), to see if said factors vary between two regions and yield/harvest/production zones. Unveiling such differences would help elucidating the disparities in losses due to diseases. More importantly, identification of associated soybean management practices in the southern U.S. would be a good starting point to investigate the feasibility of introducing such practices to the northern U.S. For instance, we are currently exploring the relationship between fungicide use and yield losses due to foliar diseases across 28 states from 1996 to 2016. Investigating fungicide use patterns may help us to identify whether fungicide use is a potential contributing factor towards the observed economic loss differences due to foliar diseases among states/regions.

Soybean rust [*Phakopsora pachyrhizi* (Syd & P. Syd.)] has previously been reported to cause moderate to severe yield losses globally (Hartman et al. 1991; Pivonia et al. 2005). Soybean rust was first confirmed on soybean in nine states (Alabama, Arkansas, Florida, Georgia, Louisiana, Missouri, Mississippi, South Carolina, and Tennessee) in the continental U.S. during November and December 2004 (Christiano et al. 2006; Schneider et al. 2005). Since then, soybean rust epidemics have been primarily restricted to states bordering the Gulf of Mexico and the disease has shown a sluggish northward progression into major soybean production regions in the U.S. (Christiano and Scherm 2007). Del Ponte and Esker (2008) unveiled the general lack of reports on negative economic effects due to soybean rust in the U.S. and the extent to which such losses may occur on a regional or continental scale across the U.S in the future. The results of the current investigation showed that soybean rust was among the least three economically damaging diseases (with bacterial blight and southern blight) over the period of 1996 to 2016. In fact, other than a few southern states such as Alabama, Florida, Georgia, Louisiana, and Texas, the losses due to soybean rust were not as conspicuous in other states. Therefore, soybean rust had not contributed to a considerable economic damage compared to diseases such as soybean cyst nematode and charcoal rot. The coordinated effort implemented since 2003 through the North Central Regional Association (NCRA, a committee designated NC-504 “Soybean Rust: A New Pest of Soybean Production”) to manage soybean rust in North America could have contributed to reducing the economic losses attributed to soybean rust through carefully implemented management practices as well as the education of individuals that deal with soybean in the allied agriculture industry (Sikora et al. 2014)

A significantly increased mean total economic loss due to all studied diseases was observed after the discovery of soybean rust, compared to the pre-discovery. This might partly be due to the concerted effort at scouting (Sikora et al. 2014), disease-monitoring network consists of soybean sentinel plots established in multiple locations within cooperating states and provinces (Giesler et al. 2007; Hershman et al. 2011; Hershman, 2009; Isard and Russo, 2007) and publicly available monitoring and prediction system known as the Soybean Rust-Pest Information Platform for Extension and Education (SBR-PIPE) (Goellner et al. 2010; Hershman et al. 2009; Isard et al. 2006; VanKirk et al. 2012). Moreover, additional factors are also responsible for the increased observation/identification of diseases following the initial observation of soybean rust including improved tools for visual identification of foliar diseases (Al-Hiary et al. 2011; Camargo and Smith, 2009; Pertot et al. 2012), improved molecular tools for disease diagnosis (Boonham et al. 2007; Niessen and Vogel 2010; Schaad et al. 2002), and development of national disease diagnostic networks during the last two decades (Stack et al. 2006). Plant disease diagnostic networks, in particular have been developed to address the problems of efficient and effective disease diagnosis and pathogen detection (Miller et al. 2009). These national and global networks, along with improved disease diagnostic tools, can in turn contribute to increased reporting of the occurrence of plant diseases. Furthermore, precise disease identification contributes to accurate estimation of yield losses associated with different diseases, which otherwise could have resulted in underestimation of losses.

Mean economic losses varied as a function of yield, harvest or production zone. Intuitively, low yield/harvest/production zones had significantly reduced mean economic losses due to diseases in comparison to high yield/harvest/production zones. Observations of a significant and positive linear correlation of mean soybean yield loss (in each state, due to all diseases) with the mean state soybean production (MT), mean soybean yield (kg ha^−1^), and mean soybean harvest area (ha) also solidified these observations. The positive relationship observed between harvest area and yield losses due to diseases may be explained by the differences in resource utilization efficiency and other related management practices. For instance, due to low farming area, farmers in low soybean growing zones can monitor their cultivation more meticulously. This in turn can potentially aid deploying more efficient fungicide application strategies in comparison to farmers in high production zones. For example, low farming hectarage allow the farmers in low harvest zones (1 and 2 for instance) to apply a proper fungicide product within the effective application window whereas farmers with extensively large soybean hectarage (harvest zones 3 and 4 for instance) would find it difficult to apply a fungicide to the whole area within the most efficacious application window. This would result in lower yield losses due to diseases in low harvest zones and vice versa. The observed positive relationship between obtained yield and yield losses (per ha basis) due to diseases may be due to several reasons. One possible explanation could be that the environmental conditions (geographic location, climatic factors, and edaphic factors) in low yield zones are not conducive for maximizing soybean yield. On the other hand, the same environmental conditions may not be conducive for disease occurrence/spread either. Therefore, less disease pressure may occur in low yield producing zones while high disease pressure may occur in high yield producing zones. For example, Oerke (2006) reported that an increase in attainable yield was often associated with an increased vulnerability to damage inflicted by insect pests and diseases. Moreover, it may also be possible that farmers in low yield zones used soybean cultivars with inherently low yield potential yet with greater level of resistance/tolerance to common soybean diseases while farmers in high yield zones used soybean cultivars with inherently high yield potential yet with lower level of resistance/tolerance to common soybean diseases. Consequently, since farmers are aware of the historically low yield potential in their zones, they may be trying to protect their yield through efficient disease management practices such as proper fungicide application. In fact, our unpublished data showed that farmers in low yield zones (particularly in yield zone 1) utilize more foliar fungicides compared to those in high yield zones. This may contribute to the reduced yield losses due to diseases in low yield producing zones.

Despite the observed differences among regions, yield/harvest/production zones, the cumulative economic losses (loss per hectare as well as total state loss) due to all diseases considered in this study showed a general increasing trend from 2005 to 2012. Interestingly, both loss types showed a decreasing trend from 2012 to 2016. These trends appear to be directly related to the soybean price fluctuation. Other than the price, these trends can be associated with climate-related patterns (e.g., less drought, lower temperatures, and greater precipitation as revealed), and the success of nationwide efforts to manage soybean diseases. Therefore, we believe it is the responsibility of policy makers, funding agencies, research institutes, and researchers to maintain/accelerate these loss declining trends. We hope that the findings of this investigation will be useful for plant pathologists, soybean breeders, federal and state governments, and educators to prioritize soybean disease management research, policy, and educational efforts.

## ACKNOWLEDGEMENTS

We thank the United Soybean Board for the support of this project, as well as the USDA National Institute of Food and Federal Appropriations under Project PEN04660 and Accession number 1016474. We also would like to express our thanks to all Extension plant pathologist and soybean disease experts who contributed observations on soybean losses in their respective states over the years.

---

**Supplementary table 1.** Common names of the diseases considered in the study and Latin binomial of the pathogen(s) causing each disease.

| Disease common name | Pathogen/s causing the disease |
| --- | --- |
| Anthracnose | *Colletotrichum truncatum* and several related species |
| Bacterial diseases | *Pseudomonas savastanoi* pv. *glycines*; *P. syringae* pv. *tabaci*;<br>*Xanthomonas axonopodis* pv. *glycines*; others |
| Brown stem rot | *Phialophora gregata* |
| Cercospora leaf blight and/or purple seed stain | *Cercospora flagellaris*, *C. kikuchii*, *C. sigesbeckiae* |
| Charcoal rot | *Macrophomina phaseolina* |
| Downy mildew | *Peronospora manshurica* |
| Frogeye leaf spot | *Cercospora sojina* |
| Fusarium wilt and root rot | *Fusarium* spp. |
| Other diseases | See footnote<sup>a</sup> |
| Diaporthe-Phomopsis | *Diaporthe longicolla* |
| Phytophthora root and stem rot | *Phytophthora sojae* |
| Pod and stem blight | *Diaporthe sojae* |
| Rhizoctonia aerial blight | *Rhizoctonia solani* |
| Root-knot and other nematodes | See footnote<sup>b</sup> |
| Sclerotinia stem rot (White mold) | *Sclerotinia sclerotiorum* |
| Seedling diseases | See footnote<sup>c</sup> |
| Septoria brown spot | *Septoria glycines* |
| Southern blight | *Sclerotium rolfsii* |
| Soybean cyst nematode | *Heterodera glycines* |
| Soybean rust | *Phakopsora pachyrhizi* |
| Stem canker | *Diaporthe aspalathi*; *D. caulivora* |
| Sudden death syndrome | *Fusarium virguliforme* |
| Virus diseases | See footnote<sup>d</sup> |

<sup>a</sup> Includes: black root rot, *Cercospora* leaf blight, *Cylindrocladium parasticum* (red crown rot), green stem syndrome, *Neocosmospora* root rot, *Pythium* root rot, target spot, and Texas root rot.
<sup>b</sup> Includes: *Rotylenchulus reniformis* (reniform nematode), *Belonolaimus longicaudatus* (sting nematode), and *Meloidogyne* (root-knot nematodes), *Helicotylenchus* (spiral nematodes), *Hoplolaimus* (lance nematodes), *Paratrichodorus* (stubby root nematodes), and *Pratylenchus* spp. (lesion nematodes).
<sup>c</sup> Includes: seedling diseases caused by a complex of organisms such as multiple species of *Fusarium*, *Pythium*, *Phomopsis*, and/or *Rhizoctonia solani*.
<sup>d</sup> Includes: *Alfalfa mosaic virus*, *Bean pod mottle virus*, *Bean yellow mosaic virus*, *Peanut mottle virus*, *Soybean dwarf virus*, *Soybean mosaic virus*, *Soybean vein necrosis virus*, *Tobacco ringspot virus*, *Tobacco streak virus*, and *Tomato spotted wilt virus*.

**Supplementary table 2.** Total state-wide economic losses due to soybean diseases (in million USD) from 12 northern states in the United States from 1996 to 2016.

| Year | State (northern United States) |  |  |  |  |  |  |  |  |  |  |  | Total |
| --- | --- | --- | --- | --- | --- | --- | --- | --- | --- | --- | --- | --- | --- |
|  | IA | IL | IN | KS | MI | MN | ND | NE | OH | PA | SD | WI |  |
| 1996 | 1,973.51 | 289.17 | 218.39 | 21.72 | 59.67 | 110.27 | 7.88 | 54.93 | 274.27 | 15.94 | 51.82 | 12.83 | **3,090.40** |
| 1997 | 1,436.53 | 257.15 | 234.40 | 54.27 | 48.70 | 329.33 | 9.79 | 65.18 | 312.78 | 2.90 | 92.69 | 32.56 | **2,876.28** |
| 1998 | 1,927.99 | 371.33 | 211.12 | 225.19 | 52.55 | 52.31 | 7.34 | 47.37 | 80.57 | 2.37 | 77.28 | 18.24 | **3,073.66** |
| 1999 | 1,409.53 | 237.99 | 160.91 | 95.77 | 43.72 | 423.05 | 9.06 | 45.24 | 35.79 | 0.50 | 53.28 | 29.07 | **2,543.91** |
| 2000 | 498.00 | 651.62 | 188.75 | 62.92 | 39.96 | 306.78 | 25.58 | 72.58 | 190.31 | 7.78 | 65.31 | 42.72 | **2,152.31** |
| 2001 | 526.87 | 210.08 | 155.96 | 26.03 | 13.84 | 186.54 | 19.67 | 39.70 | 136.81 | 38.17 | 52.39 | 42.68 | **1,448.74** |
| 2002 | 456.41 | 476.78 | 144.03 | 79.35 | 33.67 | 296.53 | 25.81 | 63.19 | 146.59 | 64.13 | 65.30 | 54.29 | **1,906.08** |
| 2003 | 625.75 | 928.31 | 185.49 | 208.68 | 32.02 | 56.14 | 22.27 | 37.04 | 373.62 | 15.61 | 78.94 | 62.83 | **2,626.70** |
| 2004 | 869.72 | 672.42 | 189.01 | 38.70 | 20.79 | 299.75 | 22.55 | 28.93 | 248.91 | 10.70 | 53.76 | 65.65 | **2,520.89** |
| 2005 | 551.48 | 359.04 | 180.02 | 75.10 | 32.91 | 194.56 | 31.75 | 29.31 | 205.02 | 23.17 | 58.91 | 84.47 | **1,825.74** |
| 2006 | 366.52 | 584.34 | 235.73 | 124.64 | 36.52 | 438.82 | 47.11 | 31.79 | 420.26 | 12.04 | 76.55 | 118.24 | **2,492.56** |
| 2007 | 314.19 | 994.69 | 260.71 | 61.02 | 39.17 | 508.02 | 84.63 | 32.98 | 221.68 | 31.62 | 185.81 | 145.10 | **2,879.62** |
| 2008 | 371.27 | 698.68 | 388.63 | 150.20 | 145.14 | 455.86 | 56.55 | 40.89 | 659.57 | 22.41 | 240.53 | 210.84 | **3,440.57** |
| 2009 | 583.84 | 969.79 | 342.45 | 260.92 | 39.55 | 498.32 | 127.66 | 44.67 | 680.22 | 15.82 | 219.22 | 455.51 | **4,237.97** |
| 2010 | 861.65 | 1,046.72 | 300.15 | 311.63 | 402.13 | 836.18 | 214.74 | 56.72 | 344.80 | 23.61 | 392.83 | 422.16 | **5,213.32** |
| 2011 | 318.25 | 581.41 | 175.49 | 181.40 | 1,420.83 | 622.64 | 291.80 | 144.44 | 304.21 | 98.01 | 311.92 | 331.86 | **4,782.26** |
| 2012 | 700.58 | 414.13 | 171.13 | 442.20 | 707.54 | 526.79 | 131.58 | 144.55 | 319.77 | 105.41 | 398.50 | 1,216.49 | **5,278.67** |
| 2013 | 331.35 | 543.21 | 185.25 | 227.00 | 295.61 | 568.35 | 135.98 | 111.46 | 1,515.99 | 69.72 | 452.09 | 267.82 | **4,703.83** |
| 2014 | 489.50 | 568.05 | 379.80 | 165.53 | 364.93 | 534.33 | 141.56 | 115.27 | 800.78 | 72.45 | 310.51 | 298.86 | **4,241.57** |
| 2015 | 483.94 | 805.97 | 242.86 | 48.55 | 278.59 | 460.49 | 121.09 | 101.88 | 1,301.42 | 70.40 | 209.40 | 174.89 | **4,299.48** |
| 2016 | 439.35 | 704.81 | 277.76 | 80.48 | 198.41 | 436.07 | 181.46 | 87.89 | 422.07 | 56.67 | 217.33 | 239.26 | **3,341.56** |
| **Total** | **15,536.25** | **12,365.67** | **4,828.06** | **2,941.30** | **4,306.25** | **8,141.15** | **1,715.86** | **1,396.01** | **8,995.43** | **759.44** | **3,664.36** | **4,326.38** | **68,976.12** |

**Supplementary table 3.** Total state-wide economic losses due to soybean diseases (in million USD) from 16 southern states in the United States from 1996 to 2016.

| Year | State (southern United States) |  |  |  |  |  |  |  |  |  |  |  |  |  |  |  | Total |
| --- | --- | --- | --- | --- | --- | --- | --- | --- | --- | --- | --- | --- | --- | --- | --- | --- | --- |
|  | AL | AR | DE | FL | GA | KY | LA | MD | MO | MS | NC | OK | SC | TN | TX | VA |  |
| 1996 | 5.31 | 88.53 | 1.35 | 1.03 | 7.97 | 28.76 | 42.31 | 3.02 | 97.74 | 61.28 | 23.82 | 3.28 | 19.62 | 16.79 | 8.99 | 1.55 | **411.35** |
| 1997 | 7.87 | 81.27 | 2.12 | 1.18 | 5.02 | 28.88 | 54.22 | 3.81 | 133.53 | 70.59 | 33.87 | 4.77 | 14.21 | 20.41 | 10.68 | 2.02 | **474.45** |
| 1998 | 5.56 | 50.70 | 1.95 | 0.32 | 3.76 | 28.97 | 27.96 | 2.33 | 85.57 | 151.20 | 16.98 | 3.98 | 10.97 | 36.31 | 6.34 | 2.27 | **435.17** |
| 1999 | 3.10 | 44.03 | 2.52 | 0.25 | 1.77 | 17.45 | 24.90 | 6.20 | 62.42 | 66.47 | 17.30 | 4.83 | 10.06 | 9.77 | 3.98 | 2.23 | **277.28** |
| 2000 | 0.90 | 44.27 | 0.14 | 0.09 | 2.01 | 22.53 | 20.95 | 3.75 | 62.55 | 24.61 | 31.66 | 1.99 | 5.70 | 17.64 | 4.58 | 4.25 | **247.62** |
| 2001 | 1.52 | 63.92 | 2.38 | 0.13 | 1.51 | 22.24 | 10.30 | 2.17 | 47.08 | 68.23 | 28.12 | 2.31 | 9.00 | 91.40 | 1.16 | 3.21 | **354.68** |
| 2002 | 1.43 | 82.67 | 1.45 | . | 1.72 | 23.47 | 19.15 | 2.31 | 48.56 | 76.07 | 33.16 | 1.62 | 8.80 | 104.11 | 7.15 | 2.13 | **413.80** |
| 2003 | 3.86 | 90.69 | 2.13 | . | 5.52 | 23.56 | 36.64 | 3.95 | 59.98 | 68.70 | 60.79 | 2.16 | 16.87 | 136.54 | 5.83 | 8.02 | **525.24** |
| 2004 | 5.12 | 105.76 | 1.35 | 0.80 | 5.06 | 16.81 | 41.78 | 4.01 | 50.22 | 109.17 | 61.72 | 2.28 | 20.90 | 128.85 | 9.83 | 11.92 | **575.58** |
| 2005 | 2.71 | 34.35 | 1.74 | 0.13 | 4.84 | 19.55 | 16.22 | 2.82 | 40.44 | 49.08 | 31.97 | 4.02 | 7.92 | 5.07 | 6.72 | 11.03 | **238.61** |
| 2006 | 14.50 | 44.70 | 0.83 | 0.05 | 3.55 | 20.13 | 18.43 | 2.27 | 56.25 | 32.91 | 32.75 | 3.31 | 9.05 | 94.49 | 5.40 | 8.85 | **347.47** |
| 2007 | 3.80 | 65.07 | 0.98 | 0.69 | 8.75 | 30.71 | 20.54 | 1.96 | 61.27 | 22.35 | 20.61 | 5.01 | 6.94 | 35.88 | 1.93 | 11.51 | **298.00** |
| 2008 | 14.86 | 60.44 | 1.64 | 3.96 | 17.22 | 31.05 | 44.89 | 2.44 | 104.18 | 41.06 | 52.09 | 7.19 | 14.67 | 96.22 | 0.67 | 13.76 | **506.34** |
| 2009 | 19.45 | 95.69 | 2.51 | 2.07 | 32.87 | 37.05 | 57.04 | 1.36 | 119.15 | 108.04 | 51.93 | 10.05 | 15.48 | 361.94 | 0.84 | 16.01 | **931.48** |
| 2010 | 9.75 | 82.84 | 3.57 | 0.36 | 10.36 | 28.93 | 25.33 | 3.20 | 144.47 | 131.89 | 42.90 | 9.11 | 11.89 | 117.53 | 0.37 | 16.78 | **639.28** |
| 2011 | 8.71 | 122.91 | 8.53 | 0.16 | 3.35 | 35.50 | 24.90 | 3.84 | 129.80 | 78.52 | 29.97 | 3.10 | 9.15 | 91.37 | 0.00 | 22.25 | **572.06** |
| 2012 | 14.27 | 170.98 | 7.70 | 0.64 | 10.82 | 67.07 | 64.43 | 5.74 | 72.74 | 98.93 | 55.75 | 4.75 | 17.52 | 133.66 | 0.35 | 29.51 | **754.86** |
| 2013 | 18.65 | 196.60 | 9.39 | 1.18 | 5.49 | 92.83 | 56.87 | 26.68 | 466.62 | 135.89 | 44.92 | 8.43 | 8.12 | 166.26 | 0.09 | 28.48 | **1,266.50** |
| 2014 | 13.20 | 179.61 | 3.38 | 0.69 | 9.62 | 101.35 | 156.74 | 7.84 | 257.61 | 309.83 | 79.19 | 7.09 | 13.96 | 123.43 | 0.31 | 28.60 | **1,292.45** |
| 2015 | 15.46 | 156.80 | 2.86 | 0.21 | 7.17 | 69.53 | 89.95 | 7.39 | 193.60 | 167.09 | 50.09 | 6.64 | 13.12 | 100.15 | 0.10 | 22.69 | **902.85** |
| 2016 | 8.71 | 163.27 | 0.98 | 0.72 | 8.89 | 73.72 | 121.25 | 4.87 | 137.44 | 191.96 | 49.23 | 8.34 | 12.93 | 142.04 | 0.35 | 22.04 | **946.74** |
| **Total** | **178.73** | **2,025.10** | **59.51** | **14.67** | **157.27** | **820.09** | **974.79** | **101.96** | **2,431.22** | **2,063.88** | **848.86** | **104.26** | **256.88** | **2,029.87** | **75.68** | **269.11** | **12,411.81** |

**Supplementary table 4.**
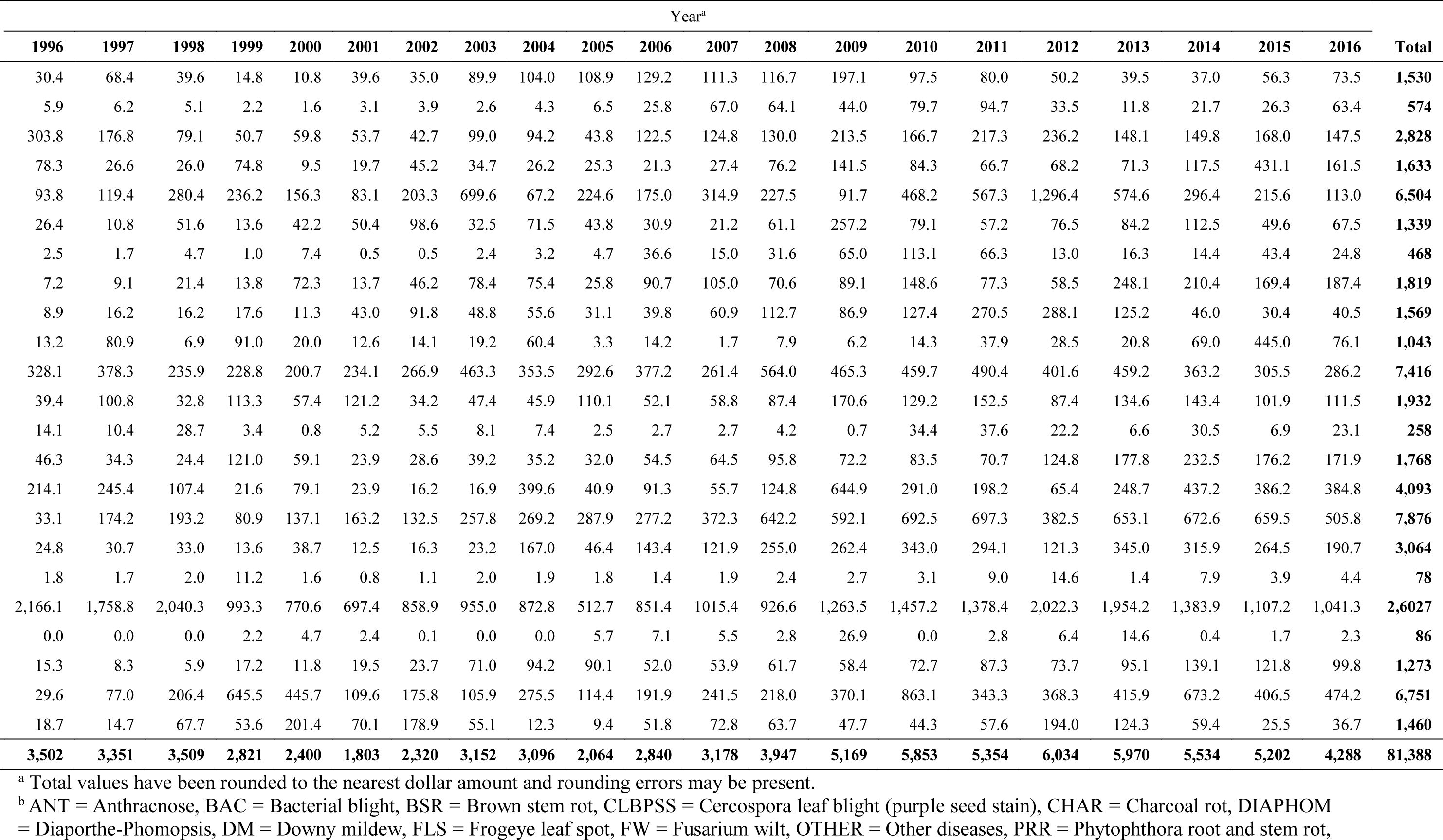
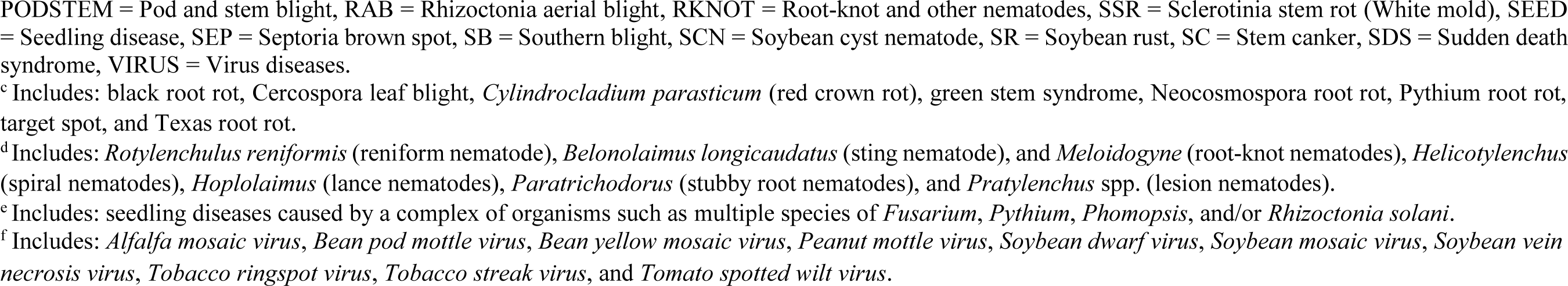
Estimated annual (from 1996 to 2016) soybean economic losses (in million USD) due to diseases across 28 states within the United States (AL, AR, DE, FL, GA, IA, IL, IN, KS, KY, LA, MD, MI, MN, MO, NC, ND, NE, OH, OK, PA, SC, SD, TN, TX, VA, and WI).

| Disease<sup>b</sup> | Year<sup>a</sup> |  |  |  |  |  |  |  |  |  |  |  |  |  |  |  |  |  |  |  |  | Total |
| --- | --- | --- | --- | --- | --- | --- | --- | --- | --- | --- | --- | --- | --- | --- | --- | --- | --- | --- | --- | --- | --- | --- |
|  | 1996 | 1997 | 1998 | 1999 | 2000 | 2001 | 2002 | 2003 | 2004 | 2005 | 2006 | 2007 | 2008 | 2009 | 2010 | 2011 | 2012 | 2013 | 2014 | 2015 | 2016 |  |
| ANT | 30.4 | 68.4 | 39.6 | 14.8 | 10.8 | 39.6 | 35.0 | 89.9 | 104.0 | 108.9 | 129.2 | 111.3 | 116.7 | 197.1 | 97.5 | 80.0 | 50.2 | 39.5 | 37.0 | 56.3 | 73.5 | **1,530** |
| BAC | 5.9 | 6.2 | 5.1 | 2.2 | 1.6 | 3.1 | 3.9 | 2.6 | 4.3 | 6.5 | 25.8 | 67.0 | 64.1 | 44.0 | 79.7 | 94.7 | 33.5 | 11.8 | 21.7 | 26.3 | 63.4 | **574** |
| BSR | 303.8 | 176.8 | 79.1 | 50.7 | 59.8 | 53.7 | 42.7 | 99.0 | 94.2 | 43.8 | 122.5 | 124.8 | 130.0 | 213.5 | 166.7 | 217.3 | 236.2 | 148.1 | 149.8 | 168.0 | 147.5 | **2,828** |
| CLBPSS | 78.3 | 26.6 | 26.0 | 74.8 | 9.5 | 19.7 | 45.2 | 34.7 | 26.2 | 25.3 | 21.3 | 27.4 | 76.2 | 141.5 | 84.3 | 66.7 | 68.2 | 71.3 | 117.5 | 431.1 | 161.5 | **1,633** |
| CHAR | 93.8 | 119.4 | 280.4 | 236.2 | 156.3 | 83.1 | 203.3 | 699.6 | 67.2 | 224.6 | 175.0 | 314.9 | 227.5 | 91.7 | 468.2 | 567.3 | 1,296.4 | 574.6 | 296.4 | 215.6 | 113.0 | **6,504** |
| DIAPHOM | 26.4 | 10.8 | 51.6 | 13.6 | 42.2 | 50.4 | 98.6 | 32.5 | 71.5 | 43.8 | 30.9 | 21.2 | 61.1 | 257.2 | 79.1 | 57.2 | 76.5 | 84.2 | 112.5 | 49.6 | 67.5 | **1,339** |
| DM | 2.5 | 1.7 | 4.7 | 1.0 | 7.4 | 0.5 | 0.5 | 2.4 | 3.2 | 4.7 | 36.6 | 15.0 | 31.6 | 65.0 | 113.1 | 66.3 | 13.0 | 16.3 | 14.4 | 43.4 | 24.8 | **468** |
| FLS | 7.2 | 9.1 | 21.4 | 13.8 | 72.3 | 13.7 | 46.2 | 78.4 | 75.4 | 25.8 | 90.7 | 105.0 | 70.6 | 89.1 | 148.6 | 77.3 | 58.5 | 248.1 | 210.4 | 169.4 | 187.4 | **1,819** |
| FW | 8.9 | 16.2 | 16.2 | 17.6 | 11.3 | 43.0 | 91.8 | 48.8 | 55.6 | 31.1 | 39.8 | 60.9 | 112.7 | 86.9 | 127.4 | 270.5 | 288.1 | 125.2 | 46.0 | 30.4 | 40.5 | **1,569** |
| OTHER<sup>c</sup> | 13.2 | 80.9 | 6.9 | 91.0 | 20.0 | 12.6 | 14.1 | 19.2 | 60.4 | 3.3 | 14.2 | 1.7 | 7.9 | 6.2 | 14.3 | 37.9 | 28.5 | 20.8 | 69.0 | 445.0 | 76.1 | **1,043** |
| PRR | 328.1 | 378.3 | 235.9 | 228.8 | 200.7 | 234.1 | 266.9 | 463.3 | 353.5 | 292.6 | 377.2 | 261.4 | 564.0 | 465.3 | 459.7 | 490.4 | 401.6 | 459.2 | 363.2 | 305.5 | 286.2 | **7,416** |
| PODSTEM | 39.4 | 100.8 | 32.8 | 113.3 | 57.4 | 121.2 | 34.2 | 47.4 | 45.9 | 110.1 | 52.1 | 58.8 | 87.4 | 170.6 | 129.2 | 152.5 | 87.4 | 134.6 | 143.4 | 101.9 | 111.5 | **1,932** |
| RAB | 14.1 | 10.4 | 28.7 | 3.4 | 0.8 | 5.2 | 5.5 | 8.1 | 7.4 | 2.5 | 2.7 | 2.7 | 4.2 | 0.7 | 34.4 | 37.6 | 22.2 | 6.6 | 30.5 | 6.9 | 23.1 | **258** |
| RKNOT<sup>d</sup> | 46.3 | 34.3 | 24.4 | 121.0 | 59.1 | 23.9 | 28.6 | 39.2 | 35.2 | 32.0 | 54.5 | 64.5 | 95.8 | 72.2 | 83.5 | 70.7 | 124.8 | 177.8 | 232.5 | 176.2 | 171.9 | **1,768** |
| SSR | 214.1 | 245.4 | 107.4 | 21.6 | 79.1 | 23.9 | 16.2 | 16.9 | 399.6 | 40.9 | 91.3 | 55.7 | 124.8 | 644.9 | 291.0 | 198.2 | 65.4 | 248.7 | 437.2 | 386.2 | 384.8 | **4,093** |
| SEED<sup>e</sup> | 33.1 | 174.2 | 193.2 | 80.9 | 137.1 | 163.2 | 132.5 | 257.8 | 269.2 | 287.9 | 277.2 | 372.3 | 642.2 | 592.1 | 692.5 | 697.3 | 382.5 | 653.1 | 672.6 | 659.5 | 505.8 | **7,876** |
| SEP | 24.8 | 30.7 | 33.0 | 13.6 | 38.7 | 12.5 | 16.3 | 23.2 | 167.0 | 46.4 | 143.4 | 121.9 | 255.0 | 262.4 | 343.0 | 294.1 | 121.3 | 345.0 | 315.9 | 264.5 | 190.7 | **3,064** |
| SB | 1.8 | 1.7 | 2.0 | 11.2 | 1.6 | 0.8 | 1.1 | 2.0 | 1.9 | 1.8 | 1.4 | 1.9 | 2.4 | 2.7 | 3.1 | 9.0 | 14.6 | 1.4 | 7.9 | 3.9 | 4.4 | **78** |
| SCN | 2,166.1 | 1,758.8 | 2,040.3 | 993.3 | 770.6 | 697.4 | 858.9 | 955.0 | 872.8 | 512.7 | 851.4 | 1015.4 | 926.6 | 1,263.5 | 1,457.2 | 1,378.4 | 2,022.3 | 1,954.2 | 1,383.9 | 1,107.2 | 1,041.3 | **2,6027** |
| SR | 0.0 | 0.0 | 0.0 | 2.2 | 4.7 | 2.4 | 0.1 | 0.0 | 0.0 | 5.7 | 7.1 | 5.5 | 2.8 | 26.9 | 0.0 | 2.8 | 6.4 | 14.6 | 0.4 | 1.7 | 2.3 | **86** |
| SC | 15.3 | 8.3 | 5.9 | 17.2 | 11.8 | 19.5 | 23.7 | 71.0 | 94.2 | 90.1 | 52.0 | 53.9 | 61.7 | 58.4 | 72.7 | 87.3 | 73.7 | 95.1 | 139.1 | 121.8 | 99.8 | **1,273** |
| SDS | 29.6 | 77.0 | 206.4 | 645.5 | 445.7 | 109.6 | 175.8 | 105.9 | 275.5 | 114.4 | 191.9 | 241.5 | 218.0 | 370.1 | 863.1 | 343.3 | 368.3 | 415.9 | 673.2 | 406.5 | 474.2 | **6,751** |
| VIRUS<sup>f</sup> | 18.7 | 14.7 | 67.7 | 53.6 | 201.4 | 70.1 | 178.9 | 55.1 | 12.3 | 9.4 | 51.8 | 72.8 | 63.7 | 47.7 | 44.3 | 57.6 | 194.0 | 124.3 | 59.4 | 25.5 | 36.7 | **1,460** |
| **Total** | **3,502** | **3,351** | **3,509** | **2,821** | **2,400** | **1,803** | **2,320** | **3,152** | **3,096** | **2,064** | **2,840** | **3,178** | **3,947** | **5,169** | **5,853** | **5,354** | **6,034** | **5,970** | **5,534** | **5,202** | **4,288** | **81,388** |

<sup>a</sup> Total values have been rounded to the nearest dollar amount and rounding errors may be present.
<sup>b</sup> ANT = Anthracnose, BAC = Bacterial blight, BSR = Brown stem rot, CLBPSS = Cercospora leaf blight (purple seed stain), CHAR = Charcoal rot, DIAPHOM = Diaporthe-Phomopsis, DM = Downy mildew, FLS = Frogeye leaf spot, FW = Fusarium wilt, OTHER = Other diseases, PRR = Phytophthora root and stem rot,
PODSTEM = Pod and stem blight, RAB = Rhizoctonia aerial blight, RKNOT = Root-knot and other nematodes, SSR = Sclerotinia stem rot (White mold), SEED = Seedling disease, SEP = Septoria brown spot, SB = Southern blight, SCN = Soybean cyst nematode, SR = Soybean rust, SC = Stem canker, SDS = Sudden death syndrome, VIRUS = Virus diseases. <sup>c</sup> Includes: black root rot, Cercospora leaf blight, *Cylindrocladium parasticum* (red crown rot), green stem syndrome, Neocosmospora root rot, Pythium root rot, target spot, and Texas root rot. <sup>d</sup> Includes: *Rotylenchulus reniformis* (reniform nematode), *Belonolaimus longicaudatus* (sting nematode), and *Meloidogyne* (root-knot nematodes), *Helicotylenchus* (spiral nematodes), *Hoplolaimus* (lance nematodes), *Paratrichodorus* (stubby root nematodes), and *Pratylenchus* spp. (lesion nematodes). <sup>e</sup> Includes: seedling diseases caused by a complex of organisms such as multiple species of *Fusarium*, *Pythium*, *Phomopsis*, and/or *Rhizoctonia solani*. <sup>f</sup> Includes: *Alfalfa mosaic virus*, *Bean pod mottle virus*, *Bean yellow mosaic virus*, *Peanut mottle virus*, *Soybean dwarf virus*, *Soybean mosaic virus*, *Soybean vein* *necrosis virus*, *Tobacco ringspot virus*, *Tobacco streak virus*, and *Tomato spotted wilt virus*.

**Supplementary table 5.**
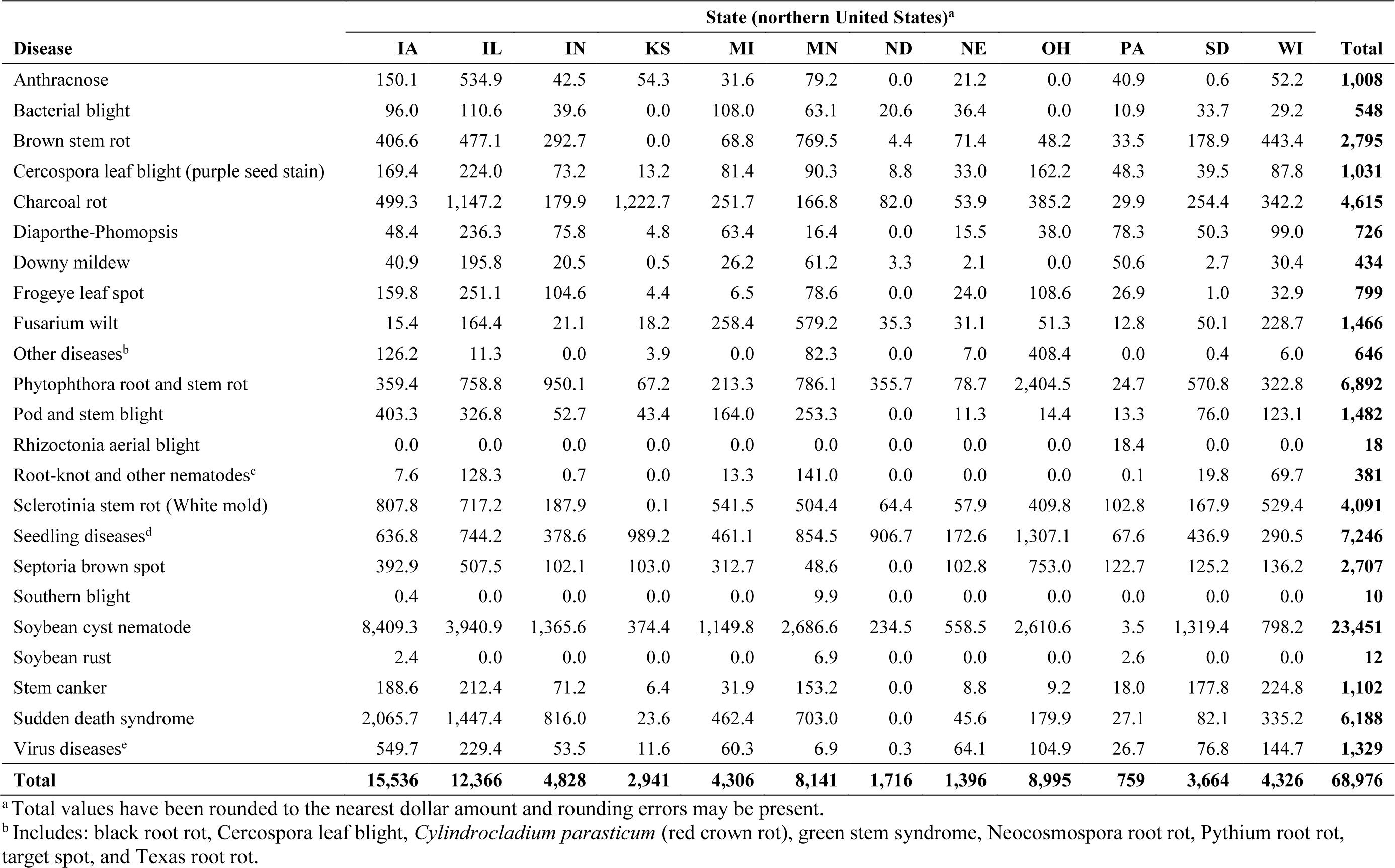
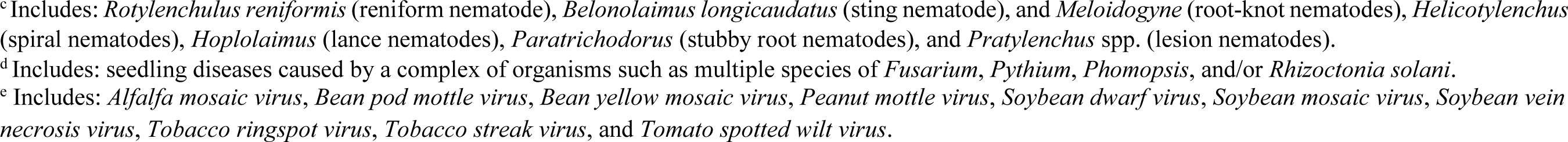
Estimated cumulative soybean economic losses from 1996 to 2016 (in million USD) as a result of diseases affecting soybean from 12 states in the northern United States.

| Disease | State (northern United States)<sup>a</sup> |  |  |  |  |  |  |  |  |  |  |  | Total |
| --- | --- | --- | --- | --- | --- | --- | --- | --- | --- | --- | --- | --- | --- |
|  | IA | IL | IN | KS | MI | MN | ND | NE | OH | PA | SD | WI |  |
| Anthrachnose | 150.1 | 534.9 | 42.5 | 54.3 | 31.6 | 79.2 | 0.0 | 21.2 | 0.0 | 40.9 | 0.6 | 52.2 | **1,008** |
| Bacterial blight | 96.0 | 110.6 | 39.6 | 0.0 | 108.0 | 63.1 | 20.6 | 36.4 | 0.0 | 10.9 | 33.7 | 29.2 | **548** |
| Brown stem rot | 406.6 | 477.1 | 292.7 | 0.0 | 68.8 | 769.5 | 4.4 | 71.4 | 48.2 | 33.5 | 178.9 | 443.4 | **2,795** |
| Cercospora leaf blight (purple seed stain) | 169.4 | 224.0 | 73.2 | 13.2 | 81.4 | 90.3 | 8.8 | 33.0 | 162.2 | 48.3 | 39.5 | 87.8 | **1,031** |
| Charcoal rot | 499.3 | 1,147.2 | 179.9 | 1,222.7 | 251.7 | 166.8 | 82.0 | 53.9 | 385.2 | 29.9 | 254.4 | 342.2 | **4,615** |
| Diaporthe-Phomopsis | 48.4 | 236.3 | 75.8 | 4.8 | 63.4 | 16.4 | 0.0 | 15.5 | 38.0 | 78.3 | 50.3 | 99.0 | **726** |
| Downy mildew | 40.9 | 195.8 | 20.5 | 0.5 | 26.2 | 61.2 | 3.3 | 2.1 | 0.0 | 50.6 | 2.7 | 30.4 | **434** |
| Frogeye leaf spot | 159.8 | 251.1 | 104.6 | 4.4 | 6.5 | 78.6 | 0.0 | 24.0 | 108.6 | 26.9 | 1.0 | 32.9 | **799** |
| Fusarium wilt | 15.4 | 164.4 | 21.1 | 18.2 | 258.4 | 579.2 | 35.3 | 31.1 | 51.3 | 12.8 | 50.1 | 228.7 | **1,466** |
| Other diseases<sup>b</sup> | 126.2 | 11.3 | 0.0 | 3.9 | 0.0 | 82.3 | 0.0 | 7.0 | 408.4 | 0.0 | 0.4 | 6.0 | **646** |
| Phytophthora root and stem rot | 359.4 | 758.8 | 950.1 | 67.2 | 213.3 | 786.1 | 355.7 | 78.7 | 2,404.5 | 24.7 | 570.8 | 322.8 | **6,892** |
| Pod and stem blight | 403.3 | 326.8 | 52.7 | 43.4 | 164.0 | 253.3 | 0.0 | 11.3 | 14.4 | 13.3 | 76.0 | 123.1 | **1,482** |
| Rhizoctonia aerial blight | 0.0 | 0.0 | 0.0 | 0.0 | 0.0 | 0.0 | 0.0 | 0.0 | 0.0 | 18.4 | 0.0 | 0.0 | **18** |
| Root-knot and other nematodes<sup>c</sup> | 7.6 | 128.3 | 0.7 | 0.0 | 13.3 | 141.0 | 0.0 | 0.0 | 0.0 | 0.1 | 19.8 | 69.7 | **381** |
| Sclerotinia stem rot (White mold) | 807.8 | 717.2 | 187.9 | 0.1 | 541.5 | 504.4 | 64.4 | 57.9 | 409.8 | 102.8 | 167.9 | 529.4 | **4,091** |
| Seedling diseases<sup>d</sup> | 636.8 | 744.2 | 378.6 | 989.2 | 461.1 | 854.5 | 906.7 | 172.6 | 1,307.1 | 67.6 | 436.9 | 290.5 | **7,246** |
| Septoria brown spot | 392.9 | 507.5 | 102.1 | 103.0 | 312.7 | 48.6 | 0.0 | 102.8 | 753.0 | 122.7 | 125.2 | 136.2 | **2,707** |
| Southern blight | 0.4 | 0.0 | 0.0 | 0.0 | 0.0 | 9.9 | 0.0 | 0.0 | 0.0 | 0.0 | 0.0 | 0.0 | **10** |
| Soybean cyst nematode | 8,409.3 | 3,940.9 | 1,365.6 | 374.4 | 1,149.8 | 2,686.6 | 234.5 | 558.5 | 2,610.6 | 3.5 | 1,319.4 | 798.2 | **23,451** |
| Soybean rust | 2.4 | 0.0 | 0.0 | 0.0 | 0.0 | 6.9 | 0.0 | 0.0 | 0.0 | 2.6 | 0.0 | 0.0 | **12** |
| Stem canker | 188.6 | 212.4 | 71.2 | 6.4 | 31.9 | 153.2 | 0.0 | 8.8 | 9.2 | 18.0 | 177.8 | 224.8 | **1,102** |
| Sudden death syndrome | 2,065.7 | 1,447.4 | 816.0 | 23.6 | 462.4 | 703.0 | 0.0 | 45.6 | 179.9 | 27.1 | 82.1 | 335.2 | **6,188** |
| Virus diseases<sup>e</sup> | 549.7 | 229.4 | 53.5 | 11.6 | 60.3 | 6.9 | 0.3 | 64.1 | 104.9 | 26.7 | 76.8 | 144.7 | **1,329** |
| **Total** | **15,536** | **12,366** | **4,828** | **2,941** | **4,306** | **8,141** | **1,716** | **1,396** | **8,995** | **759** | **3,664** | **4,326** | **68,976** |

<sup>a</sup> Total values have been rounded to the nearest dollar amount and rounding errors may be present.
<sup>b</sup> Includes: black root rot, Cercospora leaf blight, *Cylindrocladium parasticum* (red crown rot), green stem syndrome, Neocosmospora root rot, Pythium root rot, target spot, and Texas root rot.
<sup>c</sup>Includes: *Rotylenchulus reniformis* (reniform nematode), *Belonolaimus longicaudatus* (sting nematode), and *Meloidogyne* (root-knot nematodes), *Helicotylenchus* (spiral nematodes), *Hoplolaimus* (lance nematodes), *Paratrichodorus* (stubby root nematodes), and *Pratylenchus* spp. (lesion nematodes). <sup>d</sup>Includes: seedling diseases caused by a complex of organisms such as multiple species of *Fusarium*, *Pythium*, *Phomopsis*, and/or *Rhizoctonia solani*. <sup>e</sup>Includes: *Alfalfa mosaic virus*, *Bean pod mottle virus*, *Bean yellow mosaic virus*, *Peanut mottle virus*, *Soybean dwarf virus*, *Soybean mosaic virus*, *Soybean vein* *necrosis virus*, *Tobacco ringspot virus*, *Tobacco streak virus*, and *Tomato spotted wilt virus*.

**Supplementary table 6.**
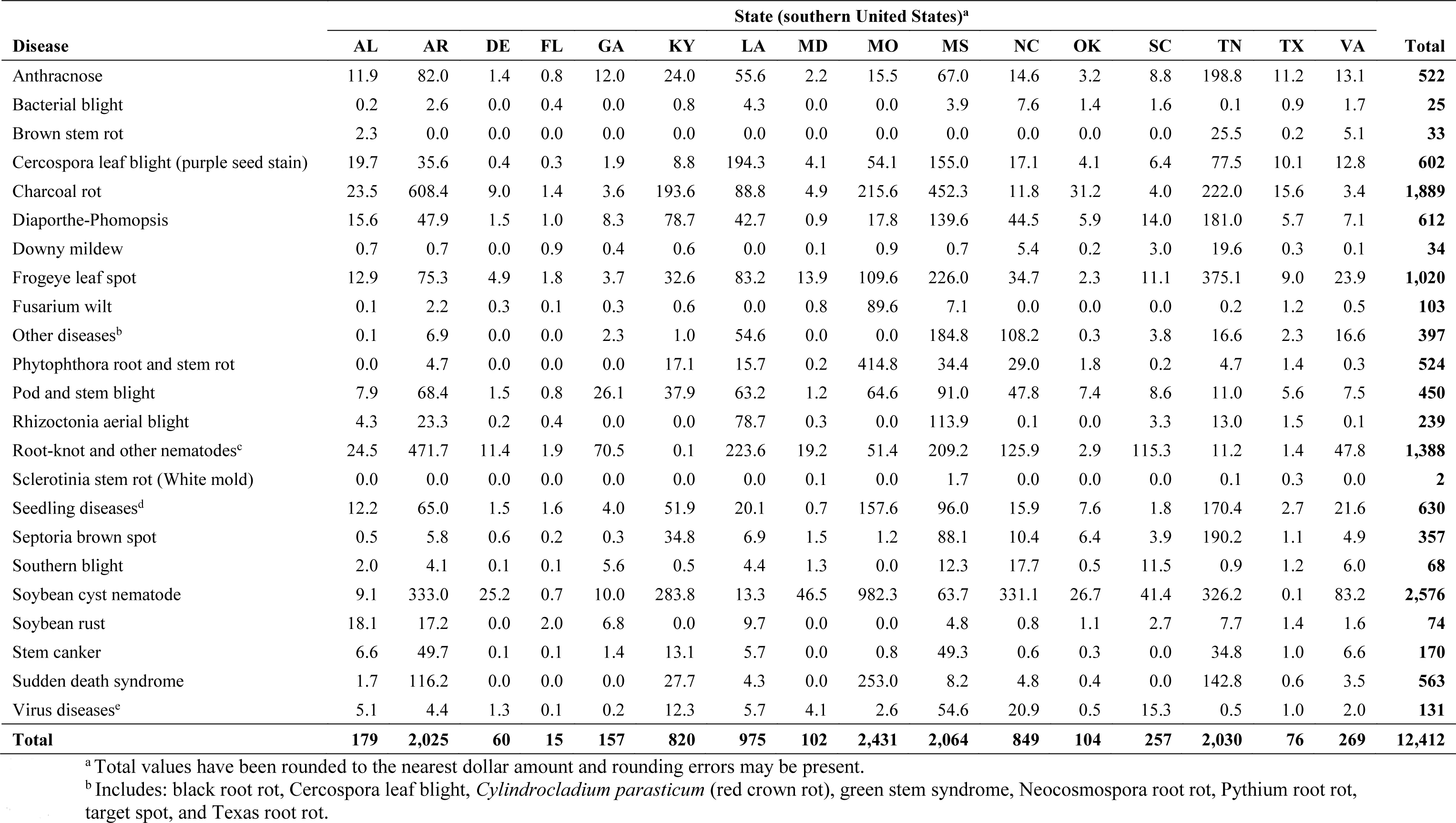
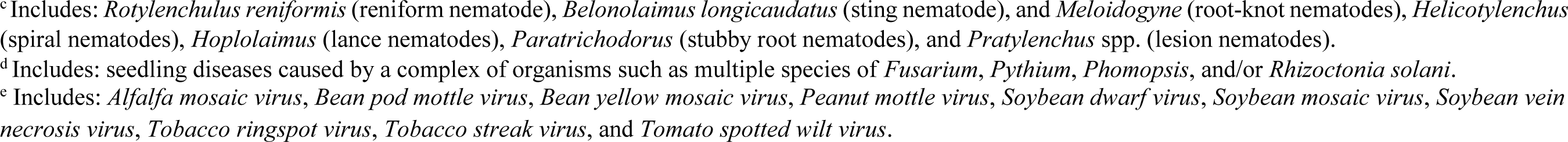
Estimated cumulative soybean economic losses from 1996 to 2016 (in million USD) as a result of diseases affecting soybean from 16 states in the southern United States.

| Disease | State (southern United States)<sup>a</sup> |  |  |  |  |  |  |  |  |  |  |  |  |  |  |  | Total |
| --- | --- | --- | --- | --- | --- | --- | --- | --- | --- | --- | --- | --- | --- | --- | --- | --- | --- |
|  | AL | AR | DE | FL | GA | KY | LA | MD | MO | MS | NC | OK | SC | TN | TX | VA |  |
| Anthrachnose | 11.9 | 82.0 | 1.4 | 0.8 | 12.0 | 24.0 | 55.6 | 2.2 | 15.5 | 67.0 | 14.6 | 3.2 | 8.8 | 198.8 | 11.2 | 13.1 | **522** |
| Bacterial blight | 0.2 | 2.6 | 0.0 | 0.4 | 0.0 | 0.8 | 4.3 | 0.0 | 0.0 | 3.9 | 7.6 | 1.4 | 1.6 | 0.1 | 0.9 | 1.7 | **25** |
| Brown stem rot | 2.3 | 0.0 | 0.0 | 0.0 | 0.0 | 0.0 | 0.0 | 0.0 | 0.0 | 0.0 | 0.0 | 0.0 | 0.0 | 25.5 | 0.2 | 5.1 | **33** |
| Cercospora leaf blight (purple seed stain) | 19.7 | 35.6 | 0.4 | 0.3 | 1.9 | 8.8 | 194.3 | 4.1 | 54.1 | 155.0 | 17.1 | 4.1 | 6.4 | 77.5 | 10.1 | 12.8 | **602** |
| Charcoal rot | 23.5 | 608.4 | 9.0 | 1.4 | 3.6 | 193.6 | 88.8 | 4.9 | 215.6 | 452.3 | 11.8 | 31.2 | 4.0 | 222.0 | 15.6 | 3.4 | **1,889** |
| Diaporthe-Phomopsis | 15.6 | 47.9 | 1.5 | 1.0 | 8.3 | 78.7 | 42.7 | 0.9 | 17.8 | 139.6 | 44.5 | 5.9 | 14.0 | 181.0 | 5.7 | 7.1 | **612** |
| Downy mildew | 0.7 | 0.7 | 0.0 | 0.9 | 0.4 | 0.6 | 0.0 | 0.1 | 0.9 | 0.7 | 5.4 | 0.2 | 3.0 | 19.6 | 0.3 | 0.1 | **34** |
| Frogeye leaf spot | 12.9 | 75.3 | 4.9 | 1.8 | 3.7 | 32.6 | 83.2 | 13.9 | 109.6 | 226.0 | 34.7 | 2.3 | 11.1 | 375.1 | 9.0 | 23.9 | **1,020** |
| Fusarium wilt | 0.1 | 2.2 | 0.3 | 0.1 | 0.3 | 0.6 | 0.0 | 0.8 | 89.6 | 7.1 | 0.0 | 0.0 | 0.0 | 0.2 | 1.2 | 0.5 | **103** |
| Other diseases<sup>b</sup> | 0.1 | 6.9 | 0.0 | 0.0 | 2.3 | 1.0 | 54.6 | 0.0 | 0.0 | 184.8 | 108.2 | 0.3 | 3.8 | 16.6 | 2.3 | 16.6 | **397** |
| Phytophthora root and stem rot | 0.0 | 4.7 | 0.0 | 0.0 | 0.0 | 17.1 | 15.7 | 0.2 | 414.8 | 34.4 | 29.0 | 1.8 | 0.2 | 4.7 | 1.4 | 0.3 | **524** |
| Pod and stem blight | 7.9 | 68.4 | 1.5 | 0.8 | 26.1 | 37.9 | 63.2 | 1.2 | 64.6 | 91.0 | 47.8 | 7.4 | 8.6 | 11.0 | 5.6 | 7.5 | **450** |
| Rhizoctonia aerial blight | 4.3 | 23.3 | 0.2 | 0.4 | 0.0 | 0.0 | 78.7 | 0.3 | 0.0 | 113.9 | 0.1 | 0.0 | 3.3 | 13.0 | 1.5 | 0.1 | **239** |
| Root-knot and other nematodes<sup>c</sup> | 24.5 | 471.7 | 11.4 | 1.9 | 70.5 | 0.1 | 223.6 | 19.2 | 51.4 | 209.2 | 125.9 | 2.9 | 115.3 | 11.2 | 1.4 | 47.8 | **1,388** |
| Sclerotinia stem rot (White mold) | 0.0 | 0.0 | 0.0 | 0.0 | 0.0 | 0.0 | 0.0 | 0.1 | 0.0 | 1.7 | 0.0 | 0.0 | 0.0 | 0.1 | 0.3 | 0.0 | **2** |
| Seedling diseases<sup>d</sup> | 12.2 | 65.0 | 1.5 | 1.6 | 4.0 | 51.9 | 20.1 | 0.7 | 157.6 | 96.0 | 15.9 | 7.6 | 1.8 | 170.4 | 2.7 | 21.6 | **630** |
| Septoria brown spot | 0.5 | 5.8 | 0.6 | 0.2 | 0.3 | 34.8 | 6.9 | 1.5 | 1.2 | 88.1 | 10.4 | 6.4 | 3.9 | 190.2 | 1.1 | 4.9 | **357** |
| Southern blight | 2.0 | 4.1 | 0.1 | 0.1 | 5.6 | 0.5 | 4.4 | 1.3 | 0.0 | 12.3 | 17.7 | 0.5 | 11.5 | 0.9 | 1.2 | 6.0 | **68** |
| Soybean cyst nematode | 9.1 | 333.0 | 25.2 | 0.7 | 10.0 | 283.8 | 13.3 | 46.5 | 982.3 | 63.7 | 331.1 | 26.7 | 41.4 | 326.2 | 0.1 | 83.2 | **2,576** |
| Soybean rust | 18.1 | 17.2 | 0.0 | 2.0 | 6.8 | 0.0 | 9.7 | 0.0 | 0.0 | 4.8 | 0.8 | 1.1 | 2.7 | 7.7 | 1.4 | 1.6 | **74** |
| Stem canker | 6.6 | 49.7 | 0.1 | 0.1 | 1.4 | 13.1 | 5.7 | 0.0 | 0.8 | 49.3 | 0.6 | 0.3 | 0.0 | 34.8 | 1.0 | 6.6 | **170** |
| Sudden death syndrome | 1.7 | 116.2 | 0.0 | 0.0 | 0.0 | 27.7 | 4.3 | 0.0 | 253.0 | 8.2 | 4.8 | 0.4 | 0.0 | 142.8 | 0.6 | 3.5 | **563** |
| Virus diseases<sup>e</sup> | 5.1 | 4.4 | 1.3 | 0.1 | 0.2 | 12.3 | 5.7 | 4.1 | 2.6 | 54.6 | 20.9 | 0.5 | 15.3 | 0.5 | 1.0 | 2.0 | **131** |
| **Total** | **179** | **2,025** | **60** | **15** | **157** | **820** | **975** | **102** | **2,431** | **2,064** | **849** | **104** | **257** | **2,030** | **76** | **269** | **12,412** |

**1136** <sup>a</sup> Total values have been rounded to the nearest dollar amount and rounding errors may be present.
**1137** <sup>b</sup> Includes: black root rot, Cercospora leaf blight, *Cylindrocladium parasticum* (red crown rot), green stem syndrome, Neocosmospora root rot, Pythium root rot, **1138** target spot, and Texas root rot.
<sup>c</sup>Includes: *Rotylenchulus reniformis* (reniform nematode), *Belonolaimus longicaudatus* (sting nematode), and *Meloidogyne* (root-knot nematodes), *Helicotylenchus* (spiral nematodes), *Hoplolaimus* (lance nematodes), *Paratrichodorus* (stubby root nematodes), and *Pratylenchus* spp. (lesion nematodes). <sup>d</sup>Includes: seedling diseases caused by a complex of organisms such as multiple species of *Fusarium*, *Pythium*, *Phomopsis*, and/or *Rhizoctonia solani*. <sup>e</sup>Includes: *Alfalfa mosaic virus*, *Bean pod mottle virus*, *Bean yellow mosaic virus*, *Peanut mottle virus*, *Soybean dwarf virus*, *Soybean mosaic virus*, *Soybean vein* *necrosis virus*, *Tobacco ringspot virus*, *Tobacco streak virus*, and *Tomato spotted wilt virus*.

**Supplementary table 7.**
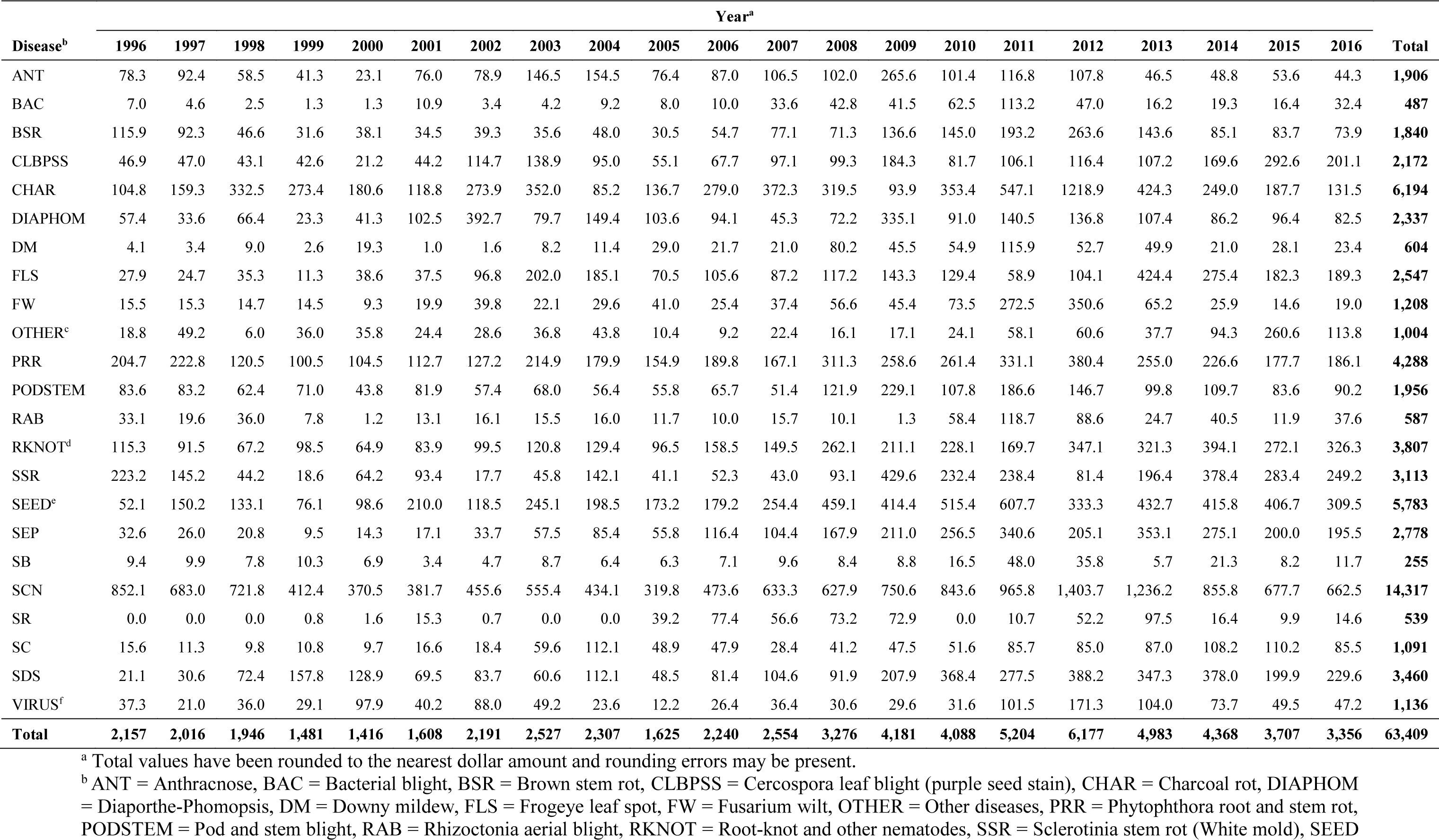
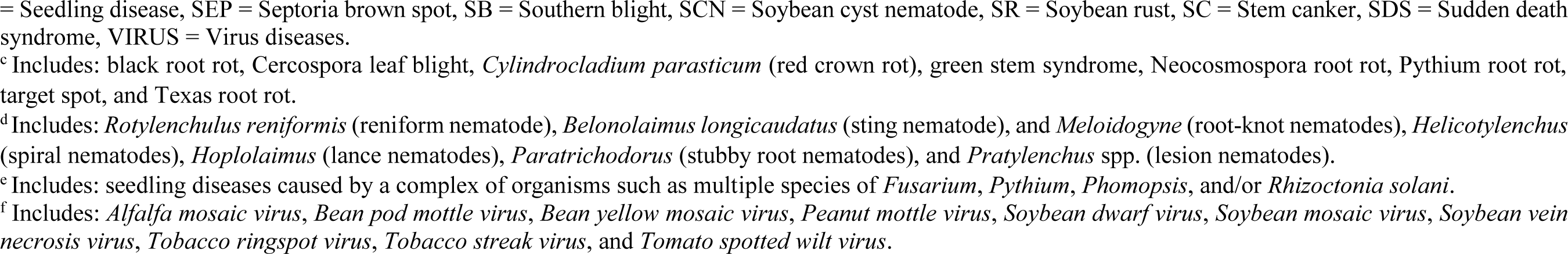
Estimated cumulative soybean economic losses (in US dollars per hectare) due to diseases in 28 states within the United States (AL, AR, DE, FL, GA, IA, IL, IN, KS, KY, LA, MD, MI, MN, MO, NC, ND, NE, OH, OK, PA, SC, SD, TN, TX, VA, and WI) from 1996 to 2016.

| Disease<sup>b</sup> | Year<sup>a</sup> |  |  |  |  |  |  |  |  |  |  |  |  |  |  |  |  |  |  |  |  | Total |
| --- | --- | --- | --- | --- | --- | --- | --- | --- | --- | --- | --- | --- | --- | --- | --- | --- | --- | --- | --- | --- | --- | --- |
|  | 1996 | 1997 | 1998 | 1999 | 2000 | 2001 | 2002 | 2003 | 2004 | 2005 | 2006 | 2007 | 2008 | 2009 | 2010 | 2011 | 2012 | 2013 | 2014 | 2015 | 2016 |  |
| ANT | 78.3 | 92.4 | 58.5 | 41.3 | 23.1 | 76.0 | 78.9 | 146.5 | 154.5 | 76.4 | 87.0 | 106.5 | 102.0 | 265.6 | 101.4 | 116.8 | 107.8 | 46.5 | 48.8 | 53.6 | 44.3 | **1,906** |
| BAC | 7.0 | 4.6 | 2.5 | 1.3 | 1.3 | 10.9 | 3.4 | 4.2 | 9.2 | 8.0 | 10.0 | 33.6 | 42.8 | 41.5 | 62.5 | 113.2 | 47.0 | 16.2 | 19.3 | 16.4 | 32.4 | **487** |
| BSR | 115.9 | 92.3 | 46.6 | 31.6 | 38.1 | 34.5 | 39.3 | 35.6 | 48.0 | 30.5 | 54.7 | 77.1 | 71.3 | 136.6 | 145.0 | 193.2 | 263.6 | 143.6 | 85.1 | 83.7 | 73.9 | **1,840** |
| CLBPSS | 46.9 | 47.0 | 43.1 | 42.6 | 21.2 | 44.2 | 114.7 | 138.9 | 95.0 | 55.1 | 67.7 | 97.1 | 99.3 | 184.3 | 81.7 | 106.1 | 116.4 | 107.2 | 169.6 | 292.6 | 201.1 | **2,172** |
| CHAR | 104.8 | 159.3 | 332.5 | 273.4 | 180.6 | 118.8 | 273.9 | 352.0 | 85.2 | 136.7 | 279.0 | 372.3 | 319.5 | 93.9 | 353.4 | 547.1 | 1218.9 | 424.3 | 249.0 | 187.7 | 131.5 | **6,194** |
| DIAPHOM | 57.4 | 33.6 | 66.4 | 23.3 | 41.3 | 102.5 | 392.7 | 79.7 | 149.4 | 103.6 | 94.1 | 45.3 | 72.2 | 335.1 | 91.0 | 140.5 | 136.8 | 107.4 | 86.2 | 96.4 | 82.5 | **2,337** |
| DM | 4.1 | 3.4 | 9.0 | 2.6 | 19.3 | 1.0 | 1.6 | 8.2 | 11.4 | 29.0 | 21.7 | 21.0 | 80.2 | 45.5 | 54.9 | 115.9 | 52.7 | 49.9 | 21.0 | 28.1 | 23.4 | **604** |
| FLS | 27.9 | 24.7 | 35.3 | 11.3 | 38.6 | 37.5 | 96.8 | 202.0 | 185.1 | 70.5 | 105.6 | 87.2 | 117.2 | 143.3 | 129.4 | 58.9 | 104.1 | 424.4 | 275.4 | 182.3 | 189.3 | **2,547** |
| FW | 15.5 | 15.3 | 14.7 | 14.5 | 9.3 | 19.9 | 39.8 | 22.1 | 29.6 | 41.0 | 25.4 | 37.4 | 56.6 | 45.4 | 73.5 | 272.5 | 350.6 | 65.2 | 25.9 | 14.6 | 19.0 | **1,208** |
| OTHER<sup>c</sup> | 18.8 | 49.2 | 6.0 | 36.0 | 35.8 | 24.4 | 28.6 | 36.8 | 43.8 | 10.4 | 9.2 | 22.4 | 16.1 | 17.1 | 24.1 | 58.1 | 60.6 | 37.7 | 94.3 | 260.6 | 113.8 | **1,004** |
| PRR | 204.7 | 222.8 | 120.5 | 100.5 | 104.5 | 112.7 | 127.2 | 214.9 | 179.9 | 154.9 | 189.8 | 167.1 | 311.3 | 258.6 | 261.4 | 331.1 | 380.4 | 255.0 | 226.6 | 177.7 | 186.1 | **4,288** |
| PODSTEM | 83.6 | 83.2 | 62.4 | 71.0 | 43.8 | 81.9 | 57.4 | 68.0 | 56.4 | 55.8 | 65.7 | 51.4 | 121.9 | 229.1 | 107.8 | 186.6 | 146.7 | 99.8 | 109.7 | 83.6 | 90.2 | **1,956** |
| RAB | 33.1 | 19.6 | 36.0 | 7.8 | 1.2 | 13.1 | 16.1 | 15.5 | 16.0 | 11.7 | 10.0 | 15.7 | 10.1 | 1.3 | 58.4 | 118.7 | 88.6 | 24.7 | 40.5 | 11.9 | 37.6 | **587** |
| RKNOT<sup>d</sup> | 115.3 | 91.5 | 67.2 | 98.5 | 64.9 | 83.9 | 99.5 | 120.8 | 129.4 | 96.5 | 158.5 | 149.5 | 262.1 | 211.1 | 228.1 | 169.7 | 347.1 | 321.3 | 394.1 | 272.1 | 326.3 | **3,807** |
| SSR | 223.2 | 145.2 | 44.2 | 18.6 | 64.2 | 93.4 | 17.7 | 45.8 | 142.1 | 41.1 | 52.3 | 43.0 | 93.1 | 429.6 | 232.4 | 238.4 | 81.4 | 196.4 | 378.4 | 283.4 | 249.2 | **3,113** |
| SEED<sup>e</sup> | 52.1 | 150.2 | 133.1 | 76.1 | 98.6 | 210.0 | 118.5 | 245.1 | 198.5 | 173.2 | 179.2 | 254.4 | 459.1 | 414.4 | 515.4 | 607.7 | 333.3 | 432.7 | 415.8 | 406.7 | 309.5 | **5,783** |
| SEP | 32.6 | 26.0 | 20.8 | 9.5 | 14.3 | 17.1 | 33.7 | 57.5 | 85.4 | 55.8 | 116.4 | 104.4 | 167.9 | 211.0 | 256.5 | 340.6 | 205.1 | 353.1 | 275.1 | 200.0 | 195.5 | **2,778** |
| SB | 9.4 | 9.9 | 7.8 | 10.3 | 6.9 | 3.4 | 4.7 | 8.7 | 6.4 | 6.3 | 7.1 | 9.6 | 8.4 | 8.8 | 16.5 | 48.0 | 35.8 | 5.7 | 21.3 | 8.2 | 11.7 | **255** |
| SCN | 852.1 | 683.0 | 721.8 | 412.4 | 370.5 | 381.7 | 455.6 | 555.4 | 434.1 | 319.8 | 473.6 | 633.3 | 627.9 | 750.6 | 843.6 | 965.8 | 1,403.7 | 1,236.2 | 855.8 | 677.7 | 662.5 | **14,317** |
| SR | 0.0 | 0.0 | 0.0 | 0.8 | 1.6 | 15.3 | 0.7 | 0.0 | 0.0 | 39.2 | 77.4 | 56.6 | 73.2 | 72.9 | 0.0 | 10.7 | 52.2 | 97.5 | 16.4 | 9.9 | 14.6 | **539** |
| SC | 15.6 | 11.3 | 9.8 | 10.8 | 9.7 | 16.6 | 18.4 | 59.6 | 112.1 | 48.9 | 47.9 | 28.4 | 41.2 | 47.5 | 51.6 | 85.7 | 85.0 | 87.0 | 108.2 | 110.2 | 85.5 | **1,091** |
| SDS | 21.1 | 30.6 | 72.4 | 157.8 | 128.9 | 69.5 | 83.7 | 60.6 | 112.1 | 48.5 | 81.4 | 104.6 | 91.9 | 207.9 | 368.4 | 277.5 | 388.2 | 347.3 | 378.0 | 199.9 | 229.6 | **3,460** |
| VIRUS<sup>f</sup> | 37.3 | 21.0 | 36.0 | 29.1 | 97.9 | 40.2 | 88.0 | 49.2 | 23.6 | 12.2 | 26.4 | 36.4 | 30.6 | 29.6 | 31.6 | 101.5 | 171.3 | 104.0 | 73.7 | 49.5 | 47.2 | **1,136** |
| **Total** | **2,157** | **2,016** | **1,946** | **1,481** | **1,416** | **1,608** | **2,191** | **2,527** | **2,307** | **1,625** | **2,240** | **2,554** | **3,276** | **4,181** | **4,088** | **5,204** | **6,177** | **4,983** | **4,368** | **3,707** | **3,356** | **63,409** |

<sup>a</sup> Total values have been rounded to the nearest dollar amount and rounding errors may be present.
<sup>b</sup> ANT = Anthracnose, BAC = Bacterial blight, BSR = Brown stem rot, CLBPSS = Cercospora leaf blight (purple seed stain), CHAR = Charcoal rot, DIAPHOM = Diaporthe-Phomopsis, DM = Downy mildew, FLS = Frogeye leaf spot, FW = Fusarium wilt, OTHER = Other diseases, PRR = Phytophthora root and stem rot, PODSTEM = Pod and stem blight, RAB = Rhizoctonia aerial blight, RKNOT = Root-knot and other nematodes, SSR = Sclerotinia stem rot (White mold), SEED
= Seedling disease, SEP = Septoria brown spot, SB = Southern blight, SCN = Soybean cyst nematode, SR = Soybean rust, SC = Stem canker, SDS = Sudden death syndrome, VIRUS = Virus diseases.
<sup>c</sup> Includes: black root rot, Cercospora leaf blight, *Cylindrocladium parasticum* (red crown rot), green stem syndrome, Neocosmospora root rot, Pythium root rot, target spot, and Texas root rot.
<sup>d</sup> Includes: *Rotylenchulus reniformis* (reniform nematode), *Belonolaimus longicaudatus* (sting nematode), and *Meloidogyne* (root-knot nematodes), *Helicotylenchus* (spiral nematodes), *Hoplolaimus* (lance nematodes), *Paratrichodorus* (stubby root nematodes), and *Pratylenchus* spp. (lesion nematodes).
<sup>e</sup> Includes: seedling diseases caused by a complex of organisms such as multiple species of *Fusarium*, *Pythium*, *Phomopsis*, and/or *Rhizoctonia solani*.
<sup>f</sup> Includes: Alfalfa mosaic virus, Bean pod mottle virus, Bean yellow mosaic virus, Peanut mottle virus, Soybean dwarf virus, Soybean mosaic virus, Soybean vein necrosis virus, Tobacco ringspot virus, Tobacco streak virus, and Tomato spotted wilt virus.

**Supplementary table 8.**
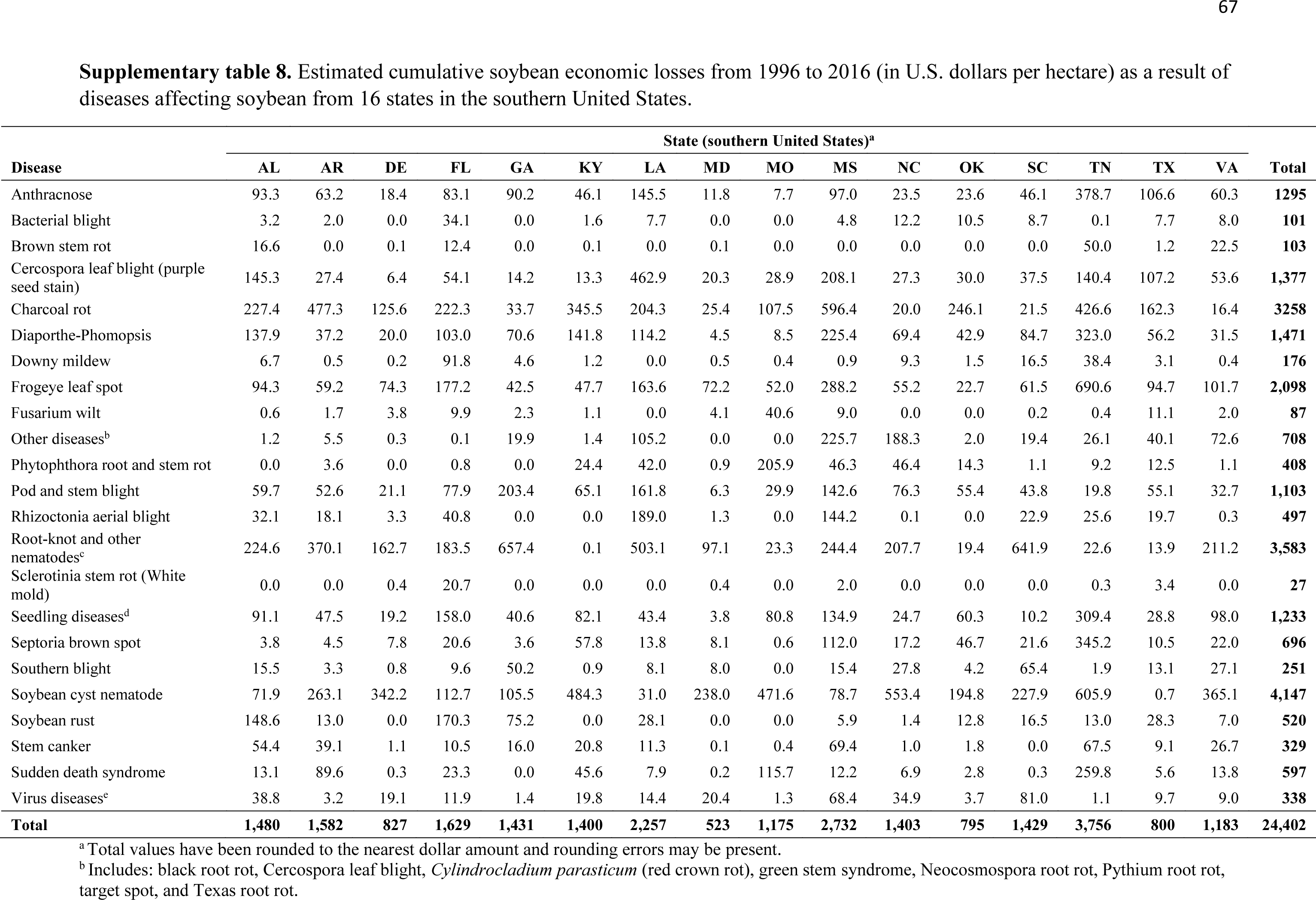
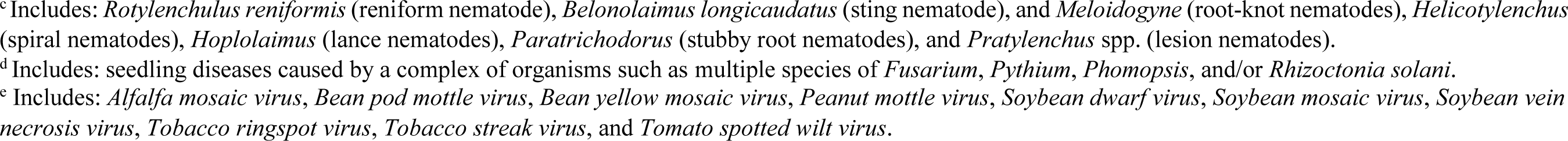
Estimated cumulative soybean economic losses from 1996 to 2016 (in U.S. dollars per hectare) as a result of diseases affecting soybean from 16 states in the southern United States.

| Disease | State (southern United States)<sup>a</sup> |  |  |  |  |  |  |  |  |  |  |  |  |  |  |  | Total |
| --- | --- | --- | --- | --- | --- | --- | --- | --- | --- | --- | --- | --- | --- | --- | --- | --- | --- |
|  | AL | AR | DE | FL | GA | KY | LA | MD | MO | MS | NC | OK | SC | TN | TX | VA |  |
| Anthracnose | 93.3 | 63.2 | 18.4 | 83.1 | 90.2 | 46.1 | 145.5 | 11.8 | 7.7 | 97.0 | 23.5 | 23.6 | 46.1 | 378.7 | 106.6 | 60.3 | **1295** |
| Bacterial blight | 3.2 | 2.0 | 0.0 | 34.1 | 0.0 | 1.6 | 7.7 | 0.0 | 0.0 | 4.8 | 12.2 | 10.5 | 8.7 | 0.1 | 7.7 | 8.0 | **101** |
| Brown stem rot | 16.6 | 0.0 | 0.1 | 12.4 | 0.0 | 0.1 | 0.0 | 0.1 | 0.0 | 0.0 | 0.0 | 0.0 | 0.0 | 50.0 | 1.2 | 22.5 | **103** |
| Cercospora leaf blight (purple seed stain) | 145.3 | 27.4 | 6.4 | 54.1 | 14.2 | 13.3 | 462.9 | 20.3 | 28.9 | 208.1 | 27.3 | 30.0 | 37.5 | 140.4 | 107.2 | 53.6 | **1,377** |
| Charcoal rot | 227.4 | 477.3 | 125.6 | 222.3 | 33.7 | 345.5 | 204.3 | 25.4 | 107.5 | 596.4 | 20.0 | 246.1 | 21.5 | 426.6 | 162.3 | 16.4 | **3258** |
| Diaporthe-Phomopsis | 137.9 | 37.2 | 20.0 | 103.0 | 70.6 | 141.8 | 114.2 | 4.5 | 8.5 | 225.4 | 69.4 | 42.9 | 84.7 | 323.0 | 56.2 | 31.5 | **1,471** |
| Downy mildew | 6.7 | 0.5 | 0.2 | 91.8 | 4.6 | 1.2 | 0.0 | 0.5 | 0.4 | 0.9 | 9.3 | 1.5 | 16.5 | 38.4 | 3.1 | 0.4 | **176** |
| Frogeye leaf spot | 94.3 | 59.2 | 74.3 | 177.2 | 42.5 | 47.7 | 163.6 | 72.2 | 52.0 | 288.2 | 55.2 | 22.7 | 61.5 | 690.6 | 94.7 | 101.7 | **2,098** |
| Fusarium wilt | 0.6 | 1.7 | 3.8 | 9.9 | 2.3 | 1.1 | 0.0 | 4.1 | 40.6 | 9.0 | 0.0 | 0.0 | 0.2 | 0.4 | 11.1 | 2.0 | **87** |
| Other diseases<sup>b</sup> | 1.2 | 5.5 | 0.3 | 0.1 | 19.9 | 1.4 | 105.2 | 0.0 | 0.0 | 225.7 | 188.3 | 2.0 | 19.4 | 26.1 | 40.1 | 72.6 | **708** |
| Phytophthora root and stem rot | 0.0 | 3.6 | 0.0 | 0.8 | 0.0 | 24.4 | 42.0 | 0.9 | 205.9 | 46.3 | 46.4 | 14.3 | 1.1 | 9.2 | 12.5 | 1.1 | **408** |
| Pod and stem blight | 59.7 | 52.6 | 21.1 | 77.9 | 203.4 | 65.1 | 161.8 | 6.3 | 29.9 | 142.6 | 76.3 | 55.4 | 43.8 | 19.8 | 55.1 | 32.7 | **1,103** |
| Rhizoctonia aerial blight | 32.1 | 18.1 | 3.3 | 40.8 | 0.0 | 0.0 | 189.0 | 1.3 | 0.0 | 144.2 | 0.1 | 0.0 | 22.9 | 25.6 | 19.7 | 0.3 | **497** |
| Root-knot and other nematodes<sup>c</sup> | 224.6 | 370.1 | 162.7 | 183.5 | 657.4 | 0.1 | 503.1 | 97.1 | 23.3 | 244.4 | 207.7 | 19.4 | 641.9 | 22.6 | 13.9 | 211.2 | **3,583** |
| Sclerotinia stem rot (White mold) | 0.0 | 0.0 | 0.4 | 20.7 | 0.0 | 0.0 | 0.0 | 0.4 | 0.0 | 2.0 | 0.0 | 0.0 | 0.0 | 0.3 | 3.4 | 0.0 | **27** |
| Seedling diseases<sup>d</sup> | 91.1 | 47.5 | 19.2 | 158.0 | 40.6 | 82.1 | 43.4 | 3.8 | 80.8 | 134.9 | 24.7 | 60.3 | 10.2 | 309.4 | 28.8 | 98.0 | **1,233** |
| Septoria brown spot | 3.8 | 4.5 | 7.8 | 20.6 | 3.6 | 57.8 | 13.8 | 8.1 | 0.6 | 112.0 | 17.2 | 46.7 | 21.6 | 345.2 | 10.5 | 22.0 | **696** |
| Southern blight | 15.5 | 3.3 | 0.8 | 9.6 | 50.2 | 0.9 | 8.1 | 8.0 | 0.0 | 15.4 | 27.8 | 4.2 | 65.4 | 1.9 | 13.1 | 27.1 | **251** |
| Soybean cyst nematode | 71.9 | 263.1 | 342.2 | 112.7 | 105.5 | 484.3 | 31.0 | 238.0 | 471.6 | 78.7 | 553.4 | 194.8 | 227.9 | 605.9 | 0.7 | 365.1 | **4,147** |
| Soybean rust | 148.6 | 13.0 | 0.0 | 170.3 | 75.2 | 0.0 | 28.1 | 0.0 | 0.0 | 5.9 | 1.4 | 12.8 | 16.5 | 13.0 | 28.3 | 7.0 | **520** |
| Stem canker | 54.4 | 39.1 | 1.1 | 10.5 | 16.0 | 20.8 | 11.3 | 0.1 | 0.4 | 69.4 | 1.0 | 1.8 | 0.0 | 67.5 | 9.1 | 26.7 | **329** |
| Sudden death syndrome | 13.1 | 89.6 | 0.3 | 23.3 | 0.0 | 45.6 | 7.9 | 0.2 | 115.7 | 12.2 | 6.9 | 2.8 | 0.3 | 259.8 | 5.6 | 13.8 | **597** |
| Virus diseases<sup>c</sup> | 38.8 | 3.2 | 19.1 | 11.9 | 1.4 | 19.8 | 14.4 | 20.4 | 1.3 | 68.4 | 34.9 | 3.7 | 81.0 | 1.1 | 9.7 | 9.0 | **338** |
| **Total** | **1,480** | **1,582** | **827** | **1,629** | **1,431** | **1,400** | **2,257** | **523** | **1,175** | **2,732** | **1,403** | **795** | **1,429** | **3,756** | **800** | **1,183** | **24,402** |

<sup>a</sup> Total values have been rounded to the nearest dollar amount and rounding errors may be present.
<sup>b</sup> Includes: black root rot, Cercospora leaf blight, *Cylindrocladium parasticum* (red crown rot), green stem syndrome, Neocosmospora root rot, Pythium root rot, target spot, and Texas root rot.
<sup>c</sup> Includes: *Rotylenchulus reniformis* (reniform nematode), *Belonolaimus longicaudatus* (sting nematode), and *Meloidogyne* (root-knot nematodes), *Helicotylenchus* (spiral nematodes), *Hoplolaimus* (lance nematodes), *Paratrichodorus* (stubby root nematodes), and *Pratylenchus* spp. (lesion nematodes).
<sup>d</sup> Includes: seedling diseases caused by a complex of organisms such as multiple species of *Fusarium*, *Pythium*, *Phomopsis*, and/or *Rhizoctonia solani*.
<sup>e</sup> Includes: *Alfalfa mosaic virus*, *Bean pod mottle virus*, *Bean yellow mosaic virus*, *Peanut mottle virus*, *Soybean dwarf virus*, *Soybean mosaic virus*, *Soybean vein necrosis virus*, *Tobacco ringspot virus*, *Tobacco streak virus*, and *Tomato spotted wilt virus*.

**Supplementary table 9.**
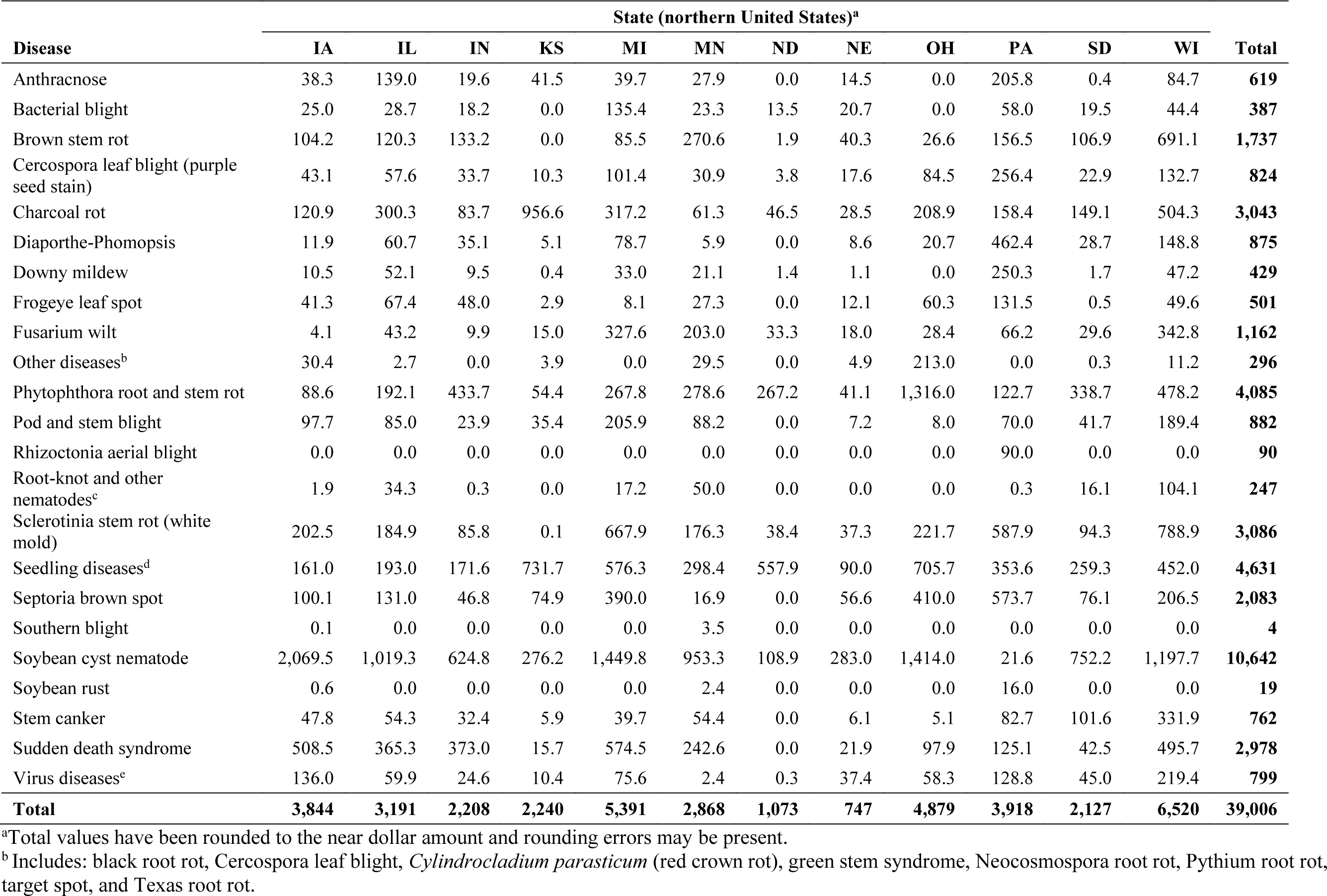
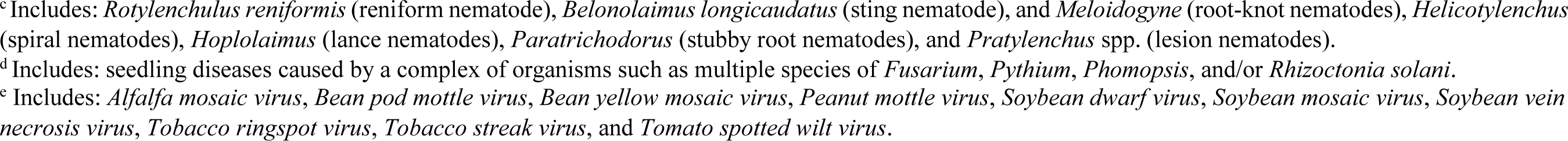
Estimated cumulative economic losses from 1996 to 2016 (in U.S. dollars per hectare) as a result of diseases affecting soybean from 12 states in the northern United States.

| Disease | State (northern United States)<sup>a</sup> |  |  |  |  |  |  |  |  |  |  |  | Total |
| --- | --- | --- | --- | --- | --- | --- | --- | --- | --- | --- | --- | --- | --- |
|  | IA | IL | IN | KS | MI | MN | ND | NE | OH | PA | SD | WI |  |
| Anthrachnose | 38.3 | 139.0 | 19.6 | 41.5 | 39.7 | 27.9 | 0.0 | 14.5 | 0.0 | 205.8 | 0.4 | 84.7 | **619** |
| Bacterial blight | 25.0 | 28.7 | 18.2 | 0.0 | 135.4 | 23.3 | 13.5 | 20.7 | 0.0 | 58.0 | 19.5 | 44.4 | **387** |
| Brown stem rot | 104.2 | 120.3 | 133.2 | 0.0 | 85.5 | 270.6 | 1.9 | 40.3 | 26.6 | 156.5 | 106.9 | 691.1 | **1,737** |
| Cercospora leaf blight (purple seed stain) | 43.1 | 57.6 | 33.7 | 10.3 | 101.4 | 30.9 | 3.8 | 17.6 | 84.5 | 256.4 | 22.9 | 132.7 | **824** |
| Charcoal rot | 120.9 | 300.3 | 83.7 | 956.6 | 317.2 | 61.3 | 46.5 | 28.5 | 208.9 | 158.4 | 149.1 | 504.3 | **3,043** |
| Diaporthe-Phomopsis | 11.9 | 60.7 | 35.1 | 5.1 | 78.7 | 5.9 | 0.0 | 8.6 | 20.7 | 462.4 | 28.7 | 148.8 | **875** |
| Downy mildew | 10.5 | 52.1 | 9.5 | 0.4 | 33.0 | 21.1 | 1.4 | 1.1 | 0.0 | 250.3 | 1.7 | 47.2 | **429** |
| Frogeye leaf spot | 41.3 | 67.4 | 48.0 | 2.9 | 8.1 | 27.3 | 0.0 | 12.1 | 60.3 | 131.5 | 0.5 | 49.6 | **501** |
| Fusarium wilt | 4.1 | 43.2 | 9.9 | 15.0 | 327.6 | 203.0 | 33.3 | 18.0 | 28.4 | 66.2 | 29.6 | 342.8 | **1,162** |
| Other diseases<sup>b</sup> | 30.4 | 2.7 | 0.0 | 3.9 | 0.0 | 29.5 | 0.0 | 4.9 | 213.0 | 0.0 | 0.3 | 11.2 | **296** |
| Phytophthora root and stem rot | 88.6 | 192.1 | 433.7 | 54.4 | 267.8 | 278.6 | 267.2 | 41.1 | 1,316.0 | 122.7 | 338.7 | 478.2 | **4,085** |
| Pod and stem blight | 97.7 | 85.0 | 23.9 | 35.4 | 205.9 | 88.2 | 0.0 | 7.2 | 8.0 | 70.0 | 41.7 | 189.4 | **882** |
| Rhizoctonia aerial blight | 0.0 | 0.0 | 0.0 | 0.0 | 0.0 | 0.0 | 0.0 | 0.0 | 0.0 | 90.0 | 0.0 | 0.0 | **90** |
| Root-knot and other nematodes<sup>c</sup> | 1.9 | 34.3 | 0.3 | 0.0 | 17.2 | 50.0 | 0.0 | 0.0 | 0.0 | 0.3 | 16.1 | 104.1 | **247** |
| Sclerotinia stem rot (white mold) | 202.5 | 184.9 | 85.8 | 0.1 | 667.9 | 176.3 | 38.4 | 37.3 | 221.7 | 587.9 | 94.3 | 788.9 | **3,086** |
| Seedling diseases<sup>d</sup> | 161.0 | 193.0 | 171.6 | 731.7 | 576.3 | 298.4 | 557.9 | 90.0 | 705.7 | 353.6 | 259.3 | 452.0 | **4,631** |
| Septoria brown spot | 100.1 | 131.0 | 46.8 | 74.9 | 390.0 | 16.9 | 0.0 | 56.6 | 410.0 | 573.7 | 76.1 | 206.5 | **2,083** |
| Southern blight | 0.1 | 0.0 | 0.0 | 0.0 | 0.0 | 3.5 | 0.0 | 0.0 | 0.0 | 0.0 | 0.0 | 0.0 | **4** |
| Soybean cyst nematode | 2,069.5 | 1,019.3 | 624.8 | 276.2 | 1,449.8 | 953.3 | 108.9 | 283.0 | 1,414.0 | 21.6 | 752.2 | 1,197.7 | **10,642** |
| Soybean rust | 0.6 | 0.0 | 0.0 | 0.0 | 0.0 | 2.4 | 0.0 | 0.0 | 0.0 | 16.0 | 0.0 | 0.0 | **19** |
| Stem canker | 47.8 | 54.3 | 32.4 | 5.9 | 39.7 | 54.4 | 0.0 | 6.1 | 5.1 | 82.7 | 101.6 | 331.9 | **762** |
| Sudden death syndrome | 508.5 | 365.3 | 373.0 | 15.7 | 574.5 | 242.6 | 0.0 | 21.9 | 97.9 | 125.1 | 42.5 | 495.7 | **2,978** |
| Virus diseases<sup>e</sup> | 136.0 | 59.9 | 24.6 | 10.4 | 75.6 | 2.4 | 0.3 | 37.4 | 58.3 | 128.8 | 45.0 | 219.4 | **799** |
| **Total** | **3,844** | **3,191** | **2,208** | **2,240** | **5,391** | **2,868** | **1,073** | **747** | **4,879** | **3,918** | **2,127** | **6,520** | **39,006** |

<sup>a</sup>Total values have been rounded to the near dollar amount and rounding errors may be present.
<sup>b</sup>Includes: black root rot, Cercospora leaf blight, *Cylindrocladium parasticum* (red crown rot), green stem syndrome, Neocosmospora root rot, Pythium root rot, target spot, and Texas root rot.
<sup>c</sup> Includes: *Rotylenchulus reniformis* (reniform nematode), *Belonolaimus longicaudatus* (sting nematode), and *Meloidogyne* (root-knot nematodes), *Helicotylenchus* (spiral nematodes), *Hoplolaimus* (lance nematodes), *Paratrichodorus* (stubby root nematodes), and *Pratylenchus* spp. (lesion nematodes).
<sup>d</sup> Includes: seedling diseases caused by a complex of organisms such as multiple species of *Fusarium*, *Pythium*, *Phomopsis*, and/or *Rhizoctonia solani*.
<sup>e</sup> Includes: *Alfalfa mosaic virus*, *Bean pod mottle virus*, *Bean yellow mosaic virus*, *Peanut mottle virus*, *Soybean dwarf virus*, *Soybean mosaic virus*, *Soybean vein necrosis virus*, *Tobacco ringspot virus*, *Tobacco streak virus*, and *Tomato spotted wilt virus*.

**Supplementary table 10.**
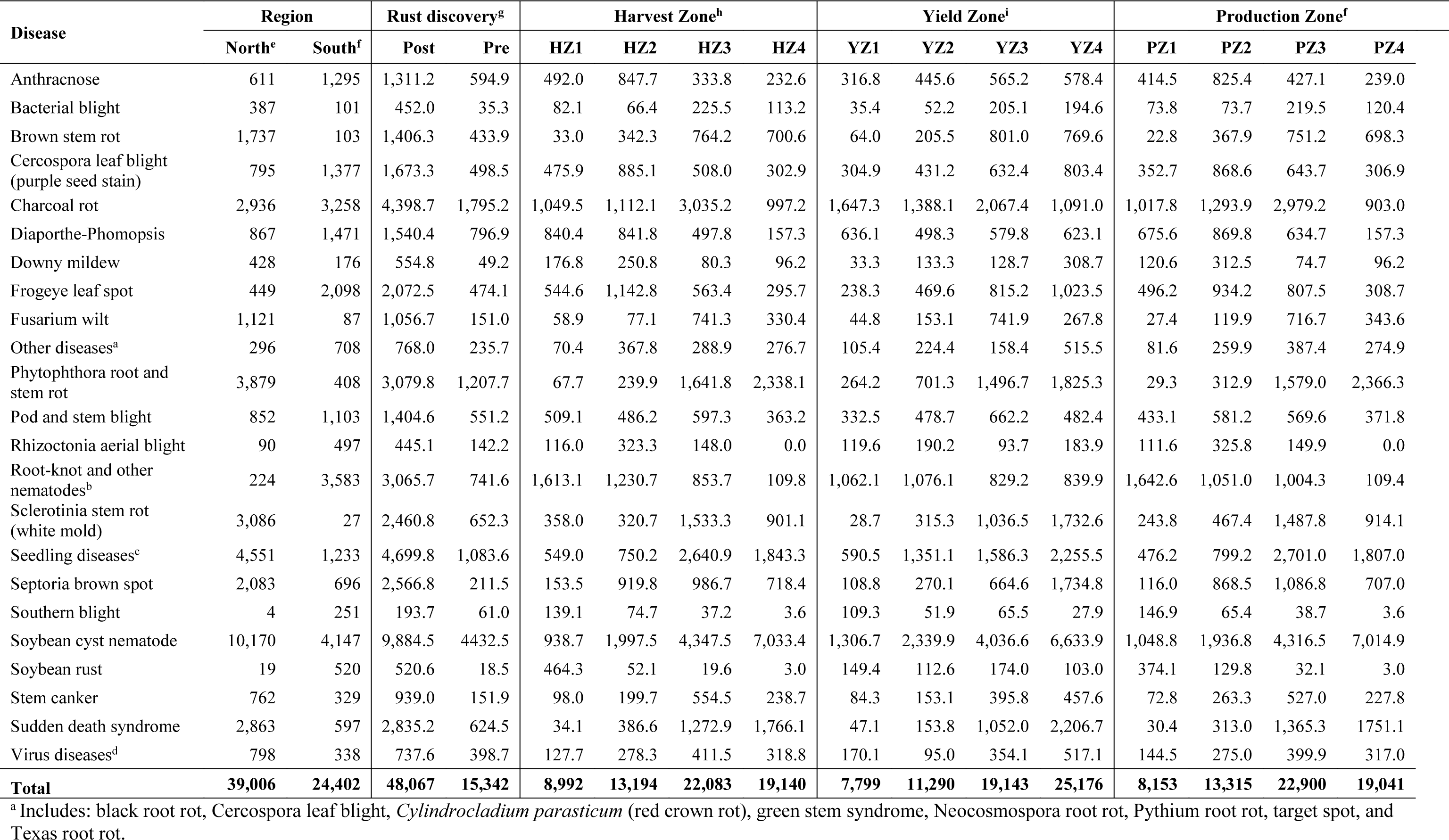
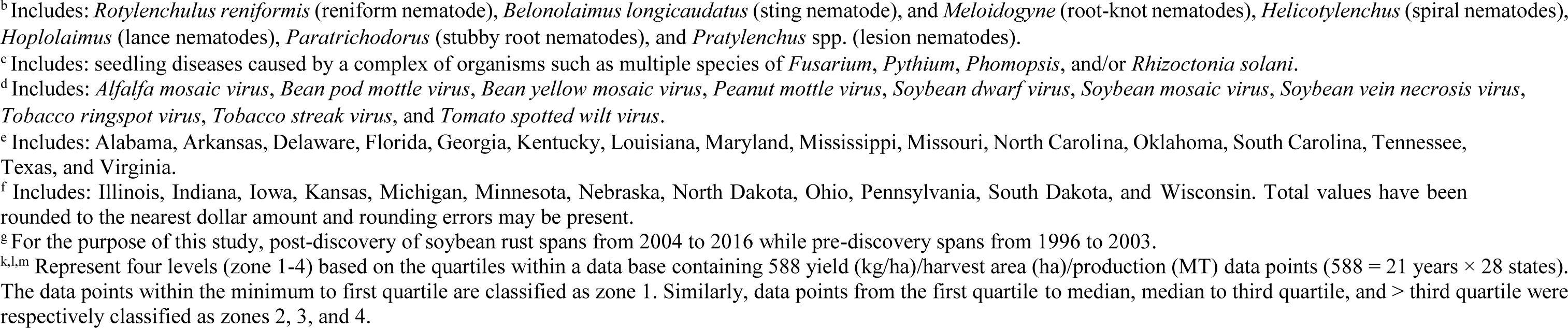
Estimated cumulative economic losses (in U.S. dollars per hectare) associated with soybean diseases (n=23) observed from a total of 28 states within each of two regions in the United States, pre- and post-discovery of soybean rust, harvest/yield/production zones (factors) of a period from 1996 to 2016.

| Disease | Region |  | Rust discovery<sup>g</sup> |  | Harvest Zone<sup>h</sup> |  |  |  | Yield Zone<sup>i</sup> |  |  |  | Production Zone<sup>f</sup> |  |  |  |
| --- | --- | --- | --- | --- | --- | --- | --- | --- | --- | --- | --- | --- | --- | --- | --- | --- |
|  | North<sup>e</sup> | South<sup>f</sup> | Post | Pre | HZ1 | HZ2 | HZ3 | HZ4 | YZ1 | YZ2 | YZ3 | YZ4 | PZ1 | PZ2 | PZ3 | PZ4 |
| Anthracnose | 611 | 1,295 | 1,311.2 | 594.9 | 492.0 | 847.7 | 333.8 | 232.6 | 316.8 | 445.6 | 565.2 | 578.4 | 414.5 | 825.4 | 427.1 | 239.0 |
| Bacterial blight | 387 | 101 | 452.0 | 35.3 | 82.1 | 66.4 | 225.5 | 113.2 | 35.4 | 52.2 | 205.1 | 194.6 | 73.8 | 73.7 | 219.5 | 120.4 |
| Brown stem rot | 1,737 | 103 | 1,406.3 | 433.9 | 33.0 | 342.3 | 764.2 | 700.6 | 64.0 | 205.5 | 801.0 | 769.6 | 22.8 | 367.9 | 751.2 | 698.3 |
| Cercospora leaf blight (purple seed stain) | 795 | 1,377 | 1,673.3 | 498.5 | 475.9 | 885.1 | 508.0 | 302.9 | 304.9 | 431.2 | 632.4 | 803.4 | 352.7 | 868.6 | 643.7 | 306.9 |
| Charcoal rot | 2,936 | 3,258 | 4,398.7 | 1,795.2 | 1,049.5 | 1,112.1 | 3,035.2 | 997.2 | 1,647.3 | 1,388.1 | 2,067.4 | 1,091.0 | 1,017.8 | 1,293.9 | 2,979.2 | 903.0 |
| Diaporthe-Phomopsis | 867 | 1,471 | 1,540.4 | 796.9 | 840.4 | 841.8 | 497.8 | 157.3 | 636.1 | 498.3 | 579.8 | 623.1 | 675.6 | 869.8 | 634.7 | 157.3 |
| Downy mildew | 428 | 176 | 554.8 | 49.2 | 176.8 | 250.8 | 80.3 | 96.2 | 33.3 | 133.3 | 128.7 | 308.7 | 120.6 | 312.5 | 74.7 | 96.2 |
| Frogeye leaf spot | 449 | 2,098 | 2,072.5 | 474.1 | 544.6 | 1,142.8 | 563.4 | 295.7 | 238.3 | 469.6 | 815.2 | 1,023.5 | 496.2 | 934.2 | 807.5 | 308.7 |
| Fusarium wilt | 1,121 | 87 | 1,056.7 | 151.0 | 58.9 | 77.1 | 741.3 | 330.4 | 44.8 | 153.1 | 741.9 | 267.8 | 27.4 | 119.9 | 716.7 | 343.6 |
| Other diseases<sup>a</sup> | 296 | 708 | 768.0 | 235.7 | 70.4 | 367.8 | 288.9 | 276.7 | 105.4 | 224.4 | 158.4 | 515.5 | 81.6 | 259.9 | 387.4 | 274.9 |
| Phytophthora root and stem rot | 3,879 | 408 | 3,079.8 | 1,207.7 | 67.7 | 239.9 | 1,641.8 | 2,338.1 | 264.2 | 701.3 | 1,496.7 | 1,825.3 | 29.3 | 312.9 | 1,579.0 | 2,366.3 |
| Pod and stem blight | 852 | 1,103 | 1,404.6 | 551.2 | 509.1 | 486.2 | 597.3 | 363.2 | 332.5 | 478.7 | 662.2 | 482.4 | 433.1 | 581.2 | 569.6 | 371.8 |
| Rhizoctonia aerial blight | 90 | 497 | 445.1 | 142.2 | 116.0 | 323.3 | 148.0 | 0.0 | 119.6 | 190.2 | 93.7 | 183.9 | 111.6 | 325.8 | 149.9 | 0.0 |
| Root-knot and other nematodes<sup>b</sup> | 224 | 3,583 | 3,065.7 | 741.6 | 1,613.1 | 1,230.7 | 853.7 | 109.8 | 1,062.1 | 1,076.1 | 829.2 | 839.9 | 1,642.6 | 1,051.0 | 1,004.3 | 109.4 |
| Sclerotinia stem rot (white mold) | 3,086 | 27 | 2,460.8 | 652.3 | 358.0 | 320.7 | 1,533.3 | 901.1 | 28.7 | 315.3 | 1,036.5 | 1,732.6 | 243.8 | 467.4 | 1,487.8 | 914.1 |
| Seedling diseases<sup>c</sup> | 4,551 | 1,233 | 4,699.8 | 1,083.6 | 549.0 | 750.2 | 2,640.9 | 1,843.3 | 590.5 | 1,351.1 | 1,586.3 | 2,255.5 | 476.2 | 799.2 | 2,701.0 | 1,807.0 |
| Septoria brown spot | 2,083 | 696 | 2,566.8 | 211.5 | 153.5 | 919.8 | 986.7 | 718.4 | 108.8 | 270.1 | 664.6 | 1,734.8 | 116.0 | 868.5 | 1,086.8 | 707.0 |
| Southern blight | 4 | 251 | 193.7 | 61.0 | 139.1 | 74.7 | 37.2 | 3.6 | 109.3 | 51.9 | 65.5 | 27.9 | 146.9 | 65.4 | 38.7 | 3.6 |
| Soybean cyst nematode | 10,170 | 4,147 | 9,884.5 | 4432.5 | 938.7 | 1,997.5 | 4,347.5 | 7,033.4 | 1,306.7 | 2,339.9 | 4,036.6 | 6,633.9 | 1,048.8 | 1,936.8 | 4,316.5 | 7,014.9 |
| Soybean rust | 19 | 520 | 520.6 | 18.5 | 464.3 | 52.1 | 19.6 | 3.0 | 149.4 | 112.6 | 174.0 | 103.0 | 374.1 | 129.8 | 32.1 | 3.0 |
| Stem canker | 762 | 329 | 939.0 | 151.9 | 98.0 | 199.7 | 554.5 | 238.7 | 84.3 | 153.1 | 395.8 | 457.6 | 72.8 | 263.3 | 527.0 | 227.8 |
| Sudden death syndrome | 2,863 | 597 | 2,835.2 | 624.5 | 34.1 | 386.6 | 1,272.9 | 1,766.1 | 47.1 | 153.8 | 1,052.0 | 2,206.7 | 30.4 | 313.0 | 1,365.3 | 1751.1 |
| Virus diseases<sup>d</sup> | 798 | 338 | 737.6 | 398.7 | 127.7 | 278.3 | 411.5 | 318.8 | 170.1 | 95.0 | 354.1 | 517.1 | 144.5 | 275.0 | 399.9 | 317.0 |
| **Total** | **39,006** | **24,402** | **48,067** | **15,342** | **8,992** | **13,194** | **22,083** | **19,140** | **7,799** | **11,290** | **19,143** | **25,176** | **8,153** | **13,315** | **22,900** | **19,041** |

<sup>a</sup> Includes: black root rot, Cercospora leaf blight, *Cylindrocladium parasticum* (red crown rot), green stem syndrome, Neocosmospora root rot, Pythium root rot, target spot, and Texas root rot.
<sup>b</sup> Includes: *Rotylenchulus reniformis* (reniform nematode), *Belonolaimus longicaudatus* (sting nematode), and *Meloidogyne* (root-knot nematodes), *Helicotylenchus* (spiral nematodes), *Hoplolaimus* (lance nematodes), *Paratrichodorus* (stubby root nematodes), and *Pratylenchus* spp. (lesion nematodes).
<sup>c</sup> Includes: seedling diseases caused by a complex of organisms such as multiple species of *Fusarium*, *Pythium*, *Phomopsis*, and/or *Rhizoctonia solani*.
<sup>d</sup> Includes: *Alfalfa mosaic virus*, *Bean pod mottle virus*, *Bean yellow mosaic virus*, *Peanut mottle virus*, *Soybean dwarf virus*, *Soybean mosaic virus*, *Soybean vein necrosis virus*, *Tobacco ringspot virus*, *Tobacco streak virus*, and *Tomato spotted wilt virus*.
<sup>e</sup> Includes: Alabama, Arkansas, Delaware, Florida, Georgia, Kentucky, Louisiana, Maryland, Mississippi, Missouri, North Carolina, Oklahoma, South Carolina, Tennessee, Texas, and Virginia.
<sup>f</sup> Includes: Illinois, Indiana, Iowa, Kansas, Michigan, Minnesota, Nebraska, North Dakota, Ohio, Pennsylvania, South Dakota, and Wisconsin. Total values have been rounded to the nearest dollar amount and rounding errors may be present.
<sup>g</sup> For the purpose of this study, post-discovery of soybean rust spans from 2004 to 2016 while pre-discovery spans from 1996 to 2003.
<sup>k,l,m</sup> Represent four levels (zone 1-4) based on the quartiles within a data base containing 588 yield (kg/ha)/harvest area (ha)/production (MT) data points ( $588 = 21 \text{ years} \times 28 \text{ states}$ ). The data points within the minimum to first quartile are classified as zone 1. Similarly, data points from the first quartile to median, median to third quartile, and > third quartile were respectively classified as zones 2, 3, and 4.

**Supplementary table 11.** Estimated cumulative soybean economic losses (in U.S. dollars per hectare) due to disease categories in 28 states within the United States (AL, AR, DE, FL, GA, IA, IL, IN, KS, KY, LA, MD, MI, MN, MO, NC, ND, NE, OH, OK, PA, SC, SD, TN, TX, VA, and WI) from 1996 to 2016.

| Category | Year<sup>a</sup> |  |  |  |  |  |  |  |  |  |  |  |  |  |  |  |  |  |  |  |  | Total |
| --- | --- | --- | --- | --- | --- | --- | --- | --- | --- | --- | --- | --- | --- | --- | --- | --- | --- | --- | --- | --- | --- | --- |
|  | 1996 | 1997 | 1998 | 1999 | 2000 | 2001 | 2002 | 2003 | 2004 | 2005 | 2006 | 2007 | 2008 | 2009 | 2010 | 2011 | 2012 | 2013 | 2014 | 2015 | 2016 |  |
| Bacterial<sup>b</sup> | 7.0 | 4.6 | 2.5 | 1.3 | 1.3 | 10.9 | 3.4 | 4.2 | 9.2 | 8.0 | 10.0 | 33.6 | 42.8 | 41.5 | 62.5 | 113.2 | 47.0 | 16.2 | 19.3 | 16.4 | 32.4 | 487 |
| Foliar<sup>c</sup> | 363.9 | 329.9 | 331.5 | 210.2 | 204.4 | 388.6 | 792.6 | 716.3 | 753.2 | 497.1 | 645.6 | 585.2 | 844.0 | 1,488.1 | 881.1 | 1,194.8 | 1,010.4 | 1,310.5 | 1,042.7 | 958.4 | 878.5 | 15,426 |
| Nematode<sup>d</sup> | 967.4 | 774.5 | 789.0 | 510.9 | 435.4 | 465.6 | 555.1 | 676.2 | 563.5 | 416.3 | 632.1 | 782.8 | 890.0 | 961.7 | 1,071.7 | 1,135.5 | 1,750.8 | 1,557.5 | 1,249.9 | 949.8 | 988.8 | 18,124 |
| Stem/Root<sup>e</sup> | 762.3 | 836.9 | 781.6 | 693.6 | 640.8 | 678.8 | 723.2 | 1,044.4 | 913.9 | 681.1 | 916.8 | 1,093.9 | 1,452.4 | 1,642.7 | 2,017.6 | 2,601.2 | 3,137.2 | 1,957.2 | 1,888.3 | 1,472.1 | 1,296.0 | 27,232 |
| Virus<sup>f</sup> | 37.3 | 21.0 | 36.0 | 29.1 | 97.9 | 40.2 | 88.0 | 49.2 | 23.6 | 12.2 | 26.4 | 36.4 | 30.6 | 29.6 | 31.6 | 101.5 | 171.3 | 104.0 | 73.7 | 49.5 | 47.2 | 1,136 |
| Other<sup>g</sup> | 18.8 | 49.2 | 6.0 | 36.0 | 35.8 | 24.4 | 28.6 | 36.8 | 43.8 | 10.4 | 9.2 | 22.4 | 16.1 | 17.1 | 24.1 | 58.1 | 60.6 | 37.7 | 94.3 | 260.6 | 113.8 | 1,004 |
| Total | 2,157 | 2,016 | 1,947 | 1,481 | 1,416 | 1,609 | 2,191 | 2,527 | 2,307 | 1,625 | 2,240 | 2,554 | 3,276 | 4,181 | 4,089 | 5,204 | 6,177 | 4,983 | 4,368 | 3,707 | 3,357 | 63,409 |

<sup>a</sup> Total values have been rounded to the nearest dollar amount and rounding errors may be present.
<sup>b</sup> Includes: Bacterial blight.
<sup>c</sup> Includes: Anthracnose, Cercospora leaf blight (purple seed stain), Diaporthe-Phomopsis, Downy mildew, Frogeye leaf spot, Pod and stem blight, Rhizoctonia aerial blight, Septoria leaf spot, and Soybean rust
<sup>d</sup> Includes: *Heterodera glycine* (soybean cyst nematode), *Meloidogyne* spp. (root-knot nematodes), *Rotylenchulus reniformis* (reniform nematode), *Belonolaimus longicaudatus* (sting nematode), *Helicotylenchus* (spiral nematodes), *Hoplolaimus* (lance nematodes), *Paratrichodorus* (stubby root nematodes), and *Pratylenchus* spp. (lesion nematodes).
<sup>e</sup> Includes: Brown stem rot, Charcoal rot, Fusarium wilt, Phytophthora root and stem rot, Sclerotinia stem rot (white mold), Seedling diseases (caused by a complex of organisms such as multiple species of *Fusarium*, *Pythium*, *Phomopsis*, and/or *Rhizoctonia solani*), Southern blight, Stem canker, and Sudden death syndrome.
<sup>f</sup> Includes: *Alfalfa mosaic virus*, *Bean pod mottle virus*, *Bean yellow mosaic virus*, *Peanut mottle virus*, *Soybean dwarf virus*, *Soybean mosaic virus*, *Soybean vein necrosis virus*, *Tobacco ringspot virus*, *Tobacco streak virus*, and *Tomato spotted wilt virus*.
<sup>g</sup> Includes: black root rot, Cercospora leaf blight, *Cylindrocladium parasticum* (red crown rot), green stem syndrome, Neocosmospora root rot, Pythium root rot, target spot, and Texas root rot.

**Supplementary table 12.**
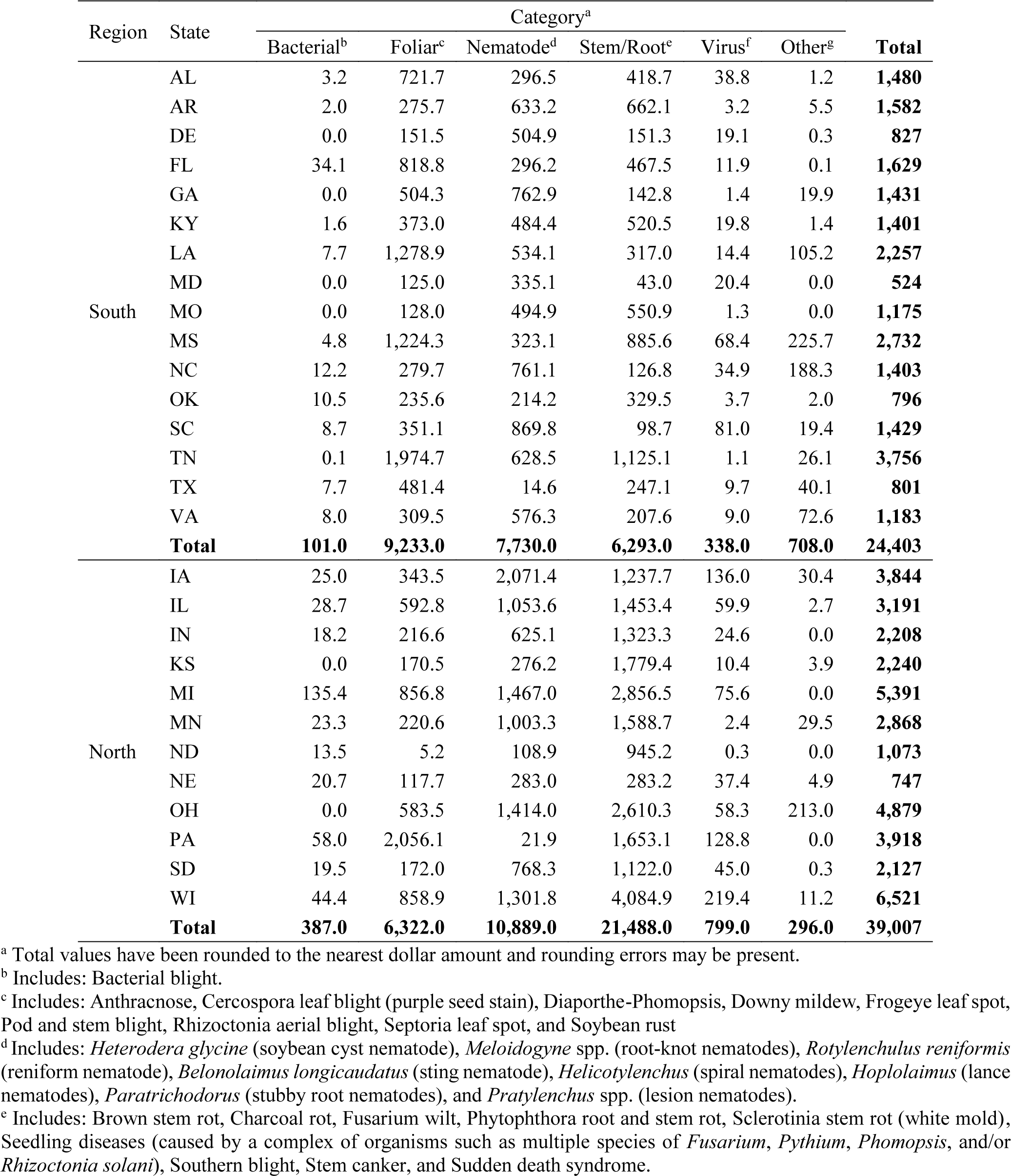
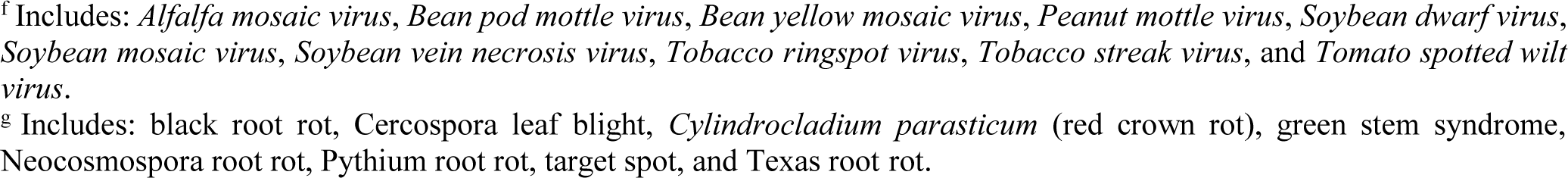
Estimated cumulative soybean economic losses from 1996 to 2016 (in U.S. ollars per hectare) due to disease categories in 16 southern and 12 northern states within the United tates.

| Region | State | Category<sup>a</sup> |  |  |  |  |  | Total |
| --- | --- | --- | --- | --- | --- | --- | --- | --- |
|  |  | Bacterial<sup>b</sup> | Foliar<sup>c</sup> | Nematode<sup>d</sup> | Stem/Root<sup>e</sup> | Virus<sup>f</sup> | Other<sup>g</sup> |  |
| South | AL | 3.2 | 721.7 | 296.5 | 418.7 | 38.8 | 1.2 | **1,480** |
|  | AR | 2.0 | 275.7 | 633.2 | 662.1 | 3.2 | 5.5 | **1,582** |
|  | DE | 0.0 | 151.5 | 504.9 | 151.3 | 19.1 | 0.3 | **827** |
|  | FL | 34.1 | 818.8 | 296.2 | 467.5 | 11.9 | 0.1 | **1,629** |
|  | GA | 0.0 | 504.3 | 762.9 | 142.8 | 1.4 | 19.9 | **1,431** |
|  | KY | 1.6 | 373.0 | 484.4 | 520.5 | 19.8 | 1.4 | **1,401** |
|  | LA | 7.7 | 1,278.9 | 534.1 | 317.0 | 14.4 | 105.2 | **2,257** |
|  | MD | 0.0 | 125.0 | 335.1 | 43.0 | 20.4 | 0.0 | **524** |
|  | MO | 0.0 | 128.0 | 494.9 | 550.9 | 1.3 | 0.0 | **1,175** |
|  | MS | 4.8 | 1,224.3 | 323.1 | 885.6 | 68.4 | 225.7 | **2,732** |
|  | NC | 12.2 | 279.7 | 761.1 | 126.8 | 34.9 | 188.3 | **1,403** |
|  | OK | 10.5 | 235.6 | 214.2 | 329.5 | 3.7 | 2.0 | **796** |
|  | SC | 8.7 | 351.1 | 869.8 | 98.7 | 81.0 | 19.4 | **1,429** |
|  | TN | 0.1 | 1,974.7 | 628.5 | 1,125.1 | 1.1 | 26.1 | **3,756** |
|  | TX | 7.7 | 481.4 | 14.6 | 247.1 | 9.7 | 40.1 | **801** |
|  | VA | 8.0 | 309.5 | 576.3 | 207.6 | 9.0 | 72.6 | **1,183** |
|  | **Total** | **101.0** | **9,233.0** | **7,730.0** | **6,293.0** | **338.0** | **708.0** | **24,403** |
| North | IA | 25.0 | 343.5 | 2,071.4 | 1,237.7 | 136.0 | 30.4 | **3,844** |
|  | IL | 28.7 | 592.8 | 1,053.6 | 1,453.4 | 59.9 | 2.7 | **3,191** |
|  | IN | 18.2 | 216.6 | 625.1 | 1,323.3 | 24.6 | 0.0 | **2,208** |
|  | KS | 0.0 | 170.5 | 276.2 | 1,779.4 | 10.4 | 3.9 | **2,240** |
|  | MI | 135.4 | 856.8 | 1,467.0 | 2,856.5 | 75.6 | 0.0 | **5,391** |
|  | MN | 23.3 | 220.6 | 1,003.3 | 1,588.7 | 2.4 | 29.5 | **2,868** |
|  | ND | 13.5 | 5.2 | 108.9 | 945.2 | 0.3 | 0.0 | **1,073** |
|  | NE | 20.7 | 117.7 | 283.0 | 283.2 | 37.4 | 4.9 | **747** |
|  | OH | 0.0 | 583.5 | 1,414.0 | 2,610.3 | 58.3 | 213.0 | **4,879** |
|  | PA | 58.0 | 2,056.1 | 21.9 | 1,653.1 | 128.8 | 0.0 | **3,918** |
|  | SD | 19.5 | 172.0 | 768.3 | 1,122.0 | 45.0 | 0.3 | **2,127** |
|  | WI | 44.4 | 858.9 | 1,301.8 | 4,084.9 | 219.4 | 11.2 | **6,521** |
|  | **Total** | **387.0** | **6,322.0** | **10,889.0** | **21,488.0** | **799.0** | **296.0** | **39,007** |

<sup>a</sup> Total values have been rounded to the nearest dollar amount and rounding errors may be present.
<sup>b</sup> Includes: Bacterial blight.
<sup>c</sup> Includes: Anthracnose, Cercospora leaf blight (purple seed stain), Diaporthe-Phomopsis, Downy mildew, Frogeye leaf spot, Pod and stem blight, Rhizoctonia aerial blight, Septoria leaf spot, and Soybean rust
<sup>d</sup> Includes: *Heterodera glycine* (soybean cyst nematode), *Meloidogyne* spp. (root-knot nematodes), *Rotylenchulus reniformis* (reniform nematode), *Belonolaimus longicaudatus* (sting nematode), *Helicotylenchus* (spiral nematodes), *Hoplolaimus* (lance nematodes), *Paratrichodorus* (stubby root nematodes), and *Pratylenchus* spp. (lesion nematodes).
<sup>e</sup> Includes: Brown stem rot, Charcoal rot, Fusarium wilt, Phytophthora root and stem rot, Sclerotinia stem rot (white mold), Seedling diseases (caused by a complex of organisms such as multiple species of *Fusarium*, *Pythium*, *Phomopsis*, and/or *Rhizoctonia solani*), Southern blight, Stem canker, and Sudden death syndrome.
<sup>f</sup> Includes: *Alfalfa mosaic virus*, *Bean pod mottle virus*, *Bean yellow mosaic virus*, *Peanut mottle virus*, *Soybean dwarf virus*, *Soybean mosaic virus*, *Soybean vein necrosis virus*, *Tobacco ringspot virus*, *Tobacco streak virus*, and *Tomato spotted wilt virus*.
<sup>g</sup> Includes: black root rot, *Cercospora* leaf blight, *Cylindrocladium parasticum* (red crown rot), green stem syndrome, *Neocosmospora* root rot, *Pythium* root rot, target spot, and Texas root rot.

**Supplementary table 13.** Estimated cumulative economic losses (from 1996 to 2016 in U.S. dollars per hectare) due to disease categories observed from a total of 28 states within each of two regions in the United States, pre- and post-discovery of soybean rust, harvest/yield/production zones.

| Category | Region |  | Rust discovery<sup>j</sup> |  | Harvest Zone<sup>k</sup> |  |  |  | Yield Zone<sup>l</sup> |  |  |  | Production Zone<sup>m</sup> |  |  |  |
| --- | --- | --- | --- | --- | --- | --- | --- | --- | --- | --- | --- | --- | --- | --- | --- | --- |
|  | North<sup>h</sup> | South<sup>i</sup> | Post | Pre | HZ1 | HZ2 | HZ3 | HZ4 | YZ1 | YZ2 | YZ3 | YZ4 | PZ1 | PZ2 | PZ3 | PZ4 |
| Bacterial<sup>a</sup> | 387.0 | 101.0 | 452.0 | 35.3 | 82.1 | 66.4 | 225.5 | 113.2 | 35.4 | 52.2 | 205.1 | 194.6 | 73.8 | 73.7 | 219.5 | 120.4 |
| Foliar<sup>b</sup> | 6,194.0 | 9,233.0 | 12,089.3 | 3,337.0 | 3,772.6 | 5,749.6 | 3,734.9 | 2,169.3 | 2,239.7 | 3,029.6 | 4,315.8 | 5,841.2 | 3,094.4 | 5,715.8 | 4,426.1 | 2,189.9 |
| Nematode<sup>c</sup> | 10,394.0 | 7,730.0 | 12,950.2 | 5,174.1 | 2,551.8 | 3,228.2 | 5,201.2 | 7,143.2 | 2,368.8 | 3,416.0 | 4,865.8 | 7,473.8 | 2,691.4 | 2,987.8 | 5,320.8 | 7,124.3 |
| Stem/Root<sup>d</sup> | 20,939.0 | 6,293.0 | 21,070.0 | 6,161.1 | 2,387.3 | 3,503.3 | 12,221.3 | 9,119.1 | 2,880.2 | 4,473.2 | 9,243.1 | 10,634.0 | 2,067.4 | 4,002.9 | 12,145.9 | 9,014.8 |
| Virus<sup>e</sup> | 798.0 | 338.0 | 737.6 | 398.7 | 127.7 | 278.3 | 411.5 | 318.8 | 170.1 | 95.0 | 354.1 | 517.1 | 144.5 | 275.0 | 399.9 | 317.0 |
| Other<sup>f</sup> | 296.0 | 708.0 | 768.0 | 235.7 | 70.4 | 367.8 | 288.9 | 276.7 | 105.4 | 224.4 | 158.4 | 515.5 | 81.6 | 259.9 | 387.4 | 274.9 |
| **Total<sup>g</sup>** | **39,008** | **24,403** | **48,067** | **15,342** | **8,992** | **13,194** | **22,083** | **19,140** | **7,800** | **11,290** | **19,142** | **25,176** | **8,153** | **13,315** | **22,900** | **19,041** |

<sup>a</sup> Includes: Bacterial blight.
<sup>b</sup> Includes: Anthracnose, Cercospora leaf blight (purple seed stain), Diaporthe-Phomopsis, Downy mildew, Frogeye leaf spot, Pod and stem blight, Rhizoctonia aerial blight, Septoria leaf spot, and Soybean rust
<sup>c</sup> Includes: *Heterodera glycine* (soybean cyst nematode), *Meloidogyne* spp. (root-knot nematodes), *Rotylenchulus reniformis* (reniform nematode), *Belonolaimus longicaudatus* (sting nematode), *Helicotylenchus* (spiral nematodes), *Hoplolaimus* (lance nematodes), *Paratrichodorus* (stubby root nematodes), and *Pratylenchus* spp. (lesion nematodes).
<sup>d</sup> Includes: Brown stem rot, Charcoal rot, Fusarium wilt, Phytophthora root and stem rot, Sclerotinia stem rot (white mold), Seedling diseases (caused by a complex of organisms such as multiple species of *Fusarium*, *Pythium*, *Phomopsis*, and/or *Rhizoctonia solani*), Southern blight, Stem canker, and Sudden death syndrome.
<sup>e</sup> Includes: *Alfalfa mosaic virus*, *Bean pod mottle virus*, *Bean yellow mosaic virus*, *Peanut mottle virus*, *Soybean dwarf virus*, *Soybean mosaic virus*, *Soybean vein necrosis virus*, *Tobacco ringspot virus*, *Tobacco streak virus*, and *Tomato spotted wilt virus*.
<sup>f</sup> Includes: black root rot, Cercospora leaf blight, *Cylindrocladium parasticum* (red crown rot), green stem syndrome, Neocosmospora root rot, Pythium root rot, target spot, and Texas root rot.
<sup>g</sup> Total values have been rounded to the nearest dollar amount and rounding errors may be present.
<sup>h</sup> Includes: Illinois, Indiana, Iowa, Kansas, Michigan, Minnesota, Nebraska, North Dakota, Ohio, Pennsylvania, South Dakota, and Wisconsin.
<sup>i</sup> Includes: Alabama, Arkansas, Delaware, Florida, Georgia, Kentucky, Louisiana, Maryland, Mississippi, Missouri, North Carolina, Oklahoma, South Carolina, Tennessee, Texas, and Virginia.
<sup>j</sup> For the purpose of this study, period of post-discovery of soybean rust spans from 2004 to 2016 while pre-discovery spans from 1996 to 2003.
<sup>k, l, m</sup> Represent four levels (zone 1-4) based on the quartiles within a data base containing 588 yield (kg/ha)/harvest area (ha)/production (MT) data points (588 = 21 years × 28 states). The data points within the minimum to first quartile are classified as zone 1. Similarly, data points from the first quartile to median, median to third quartile, and > third quartile were respectively classified as zones 2, 3, and 4.

